# Why Boolean network control tools disagree: a taxonomy of control problems

**DOI:** 10.64898/2026.03.01.703722

**Authors:** Célia Biane, Kyungduk Moon, Kangbok Lee, Loïc Paulevé

## Abstract

Boolean networks are discrete dynamical models that use Boolean states and logical functions to represent the dynamics of biological systems. A primary application of Boolean networks is to identify controls (e.g., genetic mutations or knockouts) that drive the system toward a desired phenotype. However, existing computational tools often produce inconsistent results because they rely on differing modeling assumptions. To better understand these differences, we survey existing tools and propose a taxonomy of control problems. Our taxonomy unveils hidden coverage relationships among their solutions that arise from these modeling assumptions. We provide a computational framework to empirically assess these relationships by comparing their predicted controls on a suite of artificial and biological models. Finally, we develop a coverage-consistent metric, the mutation co-occurrence score, to prioritize mutations based on their predicted impact on the phenotype. A case study on T-LGL leukemia highlights how an ensemble prediction of the score across multiple tools identifies key mutations associated with apoptosis.

**Author summary:** Boolean networks let us model gene and protein regulation with simple on/off logic. That simplicity makes them useful for asking a practical question: which controls (e.g., genetic interventions, knockouts, or forced activations) can guarantee a desired cellular phenotype such as survival or apoptosis. In practice, different software tools often return inconsistent control sets, creating a practical barrier to reproducibility and reliability of predictions. Here we provide a comprehensive survey to navigate this landscape.

We provide a comparison framework to assess coverage relationships, both theoretically and empirically, among the solutions of control tools. We classify the tools based on our new taxonomy that highlights the core differences among them. For many tools, we can theoretically determine their coverage relationships based on which components are fixed together to drive the phenotype. These new findings clearly explain subtle differences among solutions produced by various tools. We then develop a coverage-consistent metric for mutations called the mutation co-occurrence score. This new metric helps prioritize control targets based on their predicted impact on the phenotype. We also demonstrate that averaging scores from multiple tools gives reliable predictions. Our code is extensible to future tools and will facilitate the comparison of Boolean network control tools in biological research.

## 1 Introduction

A *Boolean network* (BN) is a qualitative dynamical model that uses Boolean states and logical functions to represent the dynamical behavior of a system [1, 2]. BNs are widely used to model biological systems, where each *component* represents a biological entity (e.g., a gene, protein, or RNA). The state (activity/presence) of each component is modeled as a binary value, and Boolean functions capture their interactions in a logical manner [3–6]. The state space of a BN with *n* components is the set of 2^*n*^ possible configurations, and its dynamics are represented by transitions between these states. A *phenotype* is then usually characterized as a set of states, which can be mathematically described by a Boolean function over the states. For example, in a cancer model, a phenotype may represent either the presence or absence of a signal to develop a tumor, corresponding to the activation state of a few components of a BN [7–11].

BNs are particularly useful for evaluating the impact of *controls* (i.e., perturbing the state of a few components) on the phenotype of *target states* of interest. For example, a control may represent a mutation in a gene that permanently inhibits its product and drives a cancerous phenotype. With a BN, such controls can be identified as a set of values fixed as 0 or 1 for the components. For instance, analyzing BNs under various controls enabled the identification of key genes whose mutations affect the survival of T-cell large granular lymphocyte (T-LGL) leukemia cells [7, 8], to assess how mutations in EGFR or FGFR3 genes affect the development of urinary bladder cancer [9], and the discovery of mutually exclusive or co-occurring controls involved in the development of bladder cancer [11]. These studies often manually selected controls based on biological knowledge, and the computational analysis was cross-validated with lab experiments and a literature review.

Over the past few decades, many software tools have been developed to identify biologically relevant controls [12–18]. These tools are distributed as open-source Python packages in the CoLoMoTo Docker distribution [19]. CoLoMoTo Docker has enabled researchers to easily access and compare various tools for analyzing BNs. While these tools often output almost the same sets of controls, some controls are highly inconsistent due to differing modeling assumptions. However, no study has clearly explained the reasons behind these differences. This calls for a framework to classify the tools and systematically determine relationships among their solutions according to their assumptions.

There are several difficulties in comparing controls predicted by tools in the literature. First, various tools actually solve different control problems, by considering different control targets (e.g., controlling steady states vs. controlling a larger portion of the state space) and different timing of control actions (e.g., permanent vs. temporary). The implementations may rely on approximate algorithms and usually output sparse solutions with respect to the specific control problem.

In this paper, we provide a theoretical and practical framework to classify and compare BN control tools. We focus on tools that identify controls ensuring that the BN dynamics always converge to a predetermined target set of states, independently of the initial state. We refer to this as *universal target control*.

Our framework compares tools based on the notion of solution *coverage*: we say a control *covers* another if it fixes overlapping components to the same values, and possibly fix more. Intuitively, a covering control affects a larger set of states to satisfy the phenotype property as more components are fixed. This notion of coverage is extended to tools: a tool covers another if each output control covers at least one of the output controls of the other tool.

First, we propose a taxonomy of control problems to distinguish among the different modeling assumptions in target states and control time span. Based on this taxonomy, we determine theoretical coverage relationships among different control problems, providing a rigorous explanation of differences among tools.

Then, we provide a computational framework to empirically derive coverage relationships among tools. The framework is implemented as an open-source Python package to apply the tools to a suite of BNs and compare their solutions. The framework is extensible to new tools and models, aiming to facilitate the comparison of additional and future methods.

Finally, we propose a *mutation co-occurrence score* (MCS) that quantifies the impact of mutations on a given phenotype. We measure it across multiple control tools, paving the way for a robust ensemble prediction with various methods and models. We demonstrate that this score can effectively prioritize biologically relevant mutations using a T-LGL leukemia model.

## 2 Methods

### 2.1 Formal definitions

#### 2.1.1 Boolean networks

We denote the set of Boolean values by 𝔹 := {0, 1}. A BN on *n* components is a map *f* : 𝔹^*n*^ → 𝔹^*n*^. Its *state* is a Boolean vector **x** ∈ 𝔹^*n*^ with assigned values for each of the *n* components. A *phenotype formula* (simply *phenotype* hereafter) is a Boolean function *φ* : 𝔹^*n*^ → 𝔹 that evaluates whether a given state **x** ∈ 𝔹^*n*^ satisfies the desired property or not. A *(Boolean) subspace* is specified by a vector **h** ∈ {0, 1,*\**} ^*n*^. A component *i* is *fixed* if **h**_*i*_ ∈ 𝔹 and *free* if **h**_*i*_ = *. A subspace forms a hypercube and corresponds to the set of states where free components can take any Boolean values while fixed components remain constant.

A *state transition graph* (STG) is a directed graph *G*(𝔹^*n*^, → *E*) where each node in 𝔹^*n*^ represents a state, and each arc **x**_1_ → **x**_2_ ∈ *E* represents a *transition* from state **x**_1_ to **x**_2_. There are various *update modes* to specify how to define the arc sets of the STG based on *f*. In this paper, we focus on two special types of update modes: *synchronous* and *fully asynchronous*. The *synchronous* update maps one state to the unique successor by simultaneously updating all components, i.e., there is a transition between any two different states **x** → **y** if and only if **y** = *f* (**x**). On the other hand, the *fully asynchronous* update maps each state to multiple successors where exactly one component is updated, i.e., there is a transition between any two different states **x** → **y** if and only if they differ on only one component *i*, and **y**_*i*_ = *f*_*i*_(**x**). In both cases, transitions are assumed to be only between different states. In this paper, we simply abbreviate the fully asynchronous update as *asynchronous*. Other update modes are summarized in [20].

#### 2.1.2 Target states

An important modeling assumption concerns the definition of *target states* to evaluate the phenotype. A common property employed in the literature is that target states are *trap sets*: no further transition escapes the set once entered [21]. A trap set is called a *trap space* if it is a subspace. We focus on the following five types of target states. A *fixed point* (FP) is a single state with no outgoing transition [12, 14]. This can be generalized to a *minimal trap space* (MTS), which is an inclusion-wise minimal trap space [15, 22]. An *attractor* is a mutually reachable trap set; once reached, its states are infinitely visited in the long-term transitions. Based on the update mode, we have a *synchronous attractor* (SA) or an *asynchronous attractor* (ASA) [13, 17]. A *value-propagated trap space* (VPTS) is a subspace obtained by iteratively fixing components in Boolean functions by substituting previously fixed components [18]. The formal definitions of these target states are given in “S1 Appendix”. When we say *target states of a certain type*, we indicate the set of all states included in at least one object of that type (e.g., union of all synchronous attractors). Notably, fixed points are special cases of synchronous attractors, asynchronous attractors, and minimal trap spaces involving only a single state.

**Example 1**. *Fig 1 shows a BN example with three components. Fig 1a presents the BN and the phenotype. Fig 1c and Fig 1d show the corresponding state transition graphs under the synchronous and asynchronous update, respectively. States satisfying the phenotype φ*(**x**) = **x**_2_ ∧ **x**_3_ *are colored red. This BN contains a single fixed point* 000, *and the corresponding target states are* {000}. *There are two synchronous attractors* {{000}, {111, 011, 101}} *and two asynchronous attractors* {{000}, {111, 011, 101, 001}}. *Therefore, we list their states to define target states as* {000,111, 011, 101} *and* {000, 111, 011, 101, 001}, *respectively. There are two minimal trap spaces:* 000 *and* ∗∗1. *The corresponding target states are* {000, 111, 011, 101, 001}, *which coincide with the asynchronous attractors in this example. The value-propagated trap space consists of all states* 𝔹^3^ *since no component is initially fixed*.

**Fig 1.**
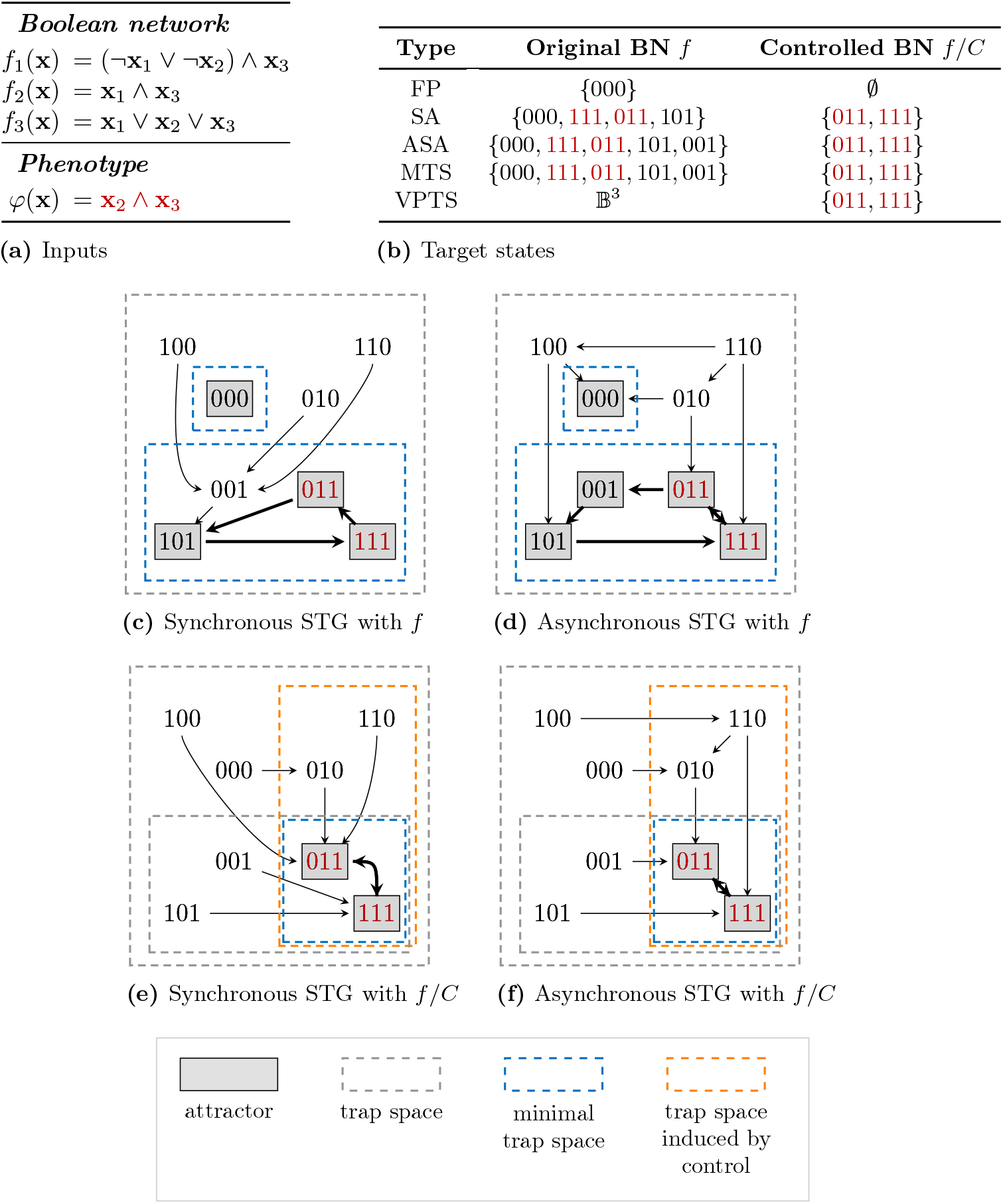
A Boolean network and the impact of control *C* = {(**x**_2_ = 1)}

#### 2.1.3 Control problem

A *control C* is a partial assignment of Boolean values to components, i.e., a subset of {(**x**_*i*_ = *b*) : *i* ∈ [1, *n*], *b* ∈ 𝔹}. For each component *i*, at most one assignment (**x**_*i*_ = *b*) can be included in *C*. Equivalently, a control can be represented as a subspace *C* ∈ {0, 1, ∗}^*n*^ where control assignments indicate fixed components and the other components are free. Fixing a component *i* to 1 or 0 represents a biological intervention that activates or inhibits the corresponding entity, respectively. The BN *f* controlled by *C*, denoted *f/C*, is defined as follows:

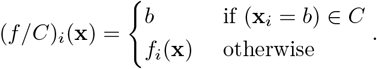

A control may have different *time spans* over which it is applied. The most common assumption is *permanent control* where the Boolean functions are fixed as constants during all transitions. Some tools apply the control temporarily, typically until reaching an attractor of *f/C*, after which the fixed components revert to their original forms [17, 23]. We term this *release control*. The time span can represent different biological scenarios. Permanent control may represent a genetic mutation that constantly activates or inhibits a gene, while release control may represent a temporary drug treatment that affects the gene expression for a limited time. Only the target states in the STG reachable from some initial state under the applied control are considered. It is important to note that the set of target states can differ significantly before and after applying a control, as well as between permanent and release controls.

We write 𝕋(*f, C*) for the target states of the chosen type and time span for BN *f* under control *C* (e.g., the set of fixed points of *f* or the set of states belonging to synchronous attractors of *f*). In this paper, we narrow down our focus to specific tools that consider a *universal target control problem*, formally defined as follows:

##### Definition 1

(The universal target control problem). *Let us consider n* ∈ ℕ *components, a BN f* : 𝔹^*n*^ → 𝔹^*n*^, *a phenotype formula φ* : 𝔹^*n*^ → 𝔹, *and a function* 𝕋 *returning the target states of interest of a given BN*.

*The set of all* valid controls *for the corresponding* universal target control problem *is the set of controls C* ∈ {0, 1, ∗}^*n*^ *where* any *target state in* 𝕋(*f, C*) *satisfies the phenotype formula:*

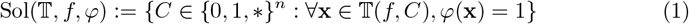

*A valid control C is* minimal *if there is no strictly smaller* valid *control C*^*′*^ ⊊ *C*.

In this paper, a *tool* is associated with one type of target states, and thus fully determines the 𝕋 function. A tool is called *exact* if, for any BN *f* and phenotype *φ*, its output only contains valid controls in Sol(𝕋, *f, φ*), including *all* the minimal valid controls.

**Example 2**. *Fig 1e-f illustrate the modified state transition graphs after applying control C* = {(**x**_2_ = 1)}. *The trap space corresponding to C is highlighted with an orange dashed box. From any state, a single transition now eventually leads to a state inside this subspace since* **x**_2_ *is fixed to 1. As the value propagation further fixes* **x**_3_ *to 1, the* VPTS *is now* ∗11 *and coincides with the unique* MTS.

*Fig 1b compares target states before and after applying the control. In the original BN, target states include at least one state that violates the phenotype. Therefore, an empty control* ∅ *(fixing no component) cannot be valid for the universal target control problem for any type of target states. On the other hand, after applying control C as a permanent control, all target states satisfy the phenotype regardless of the type. In this case, C is valid for the universal target control problem for all types of target states. If the phenotype represents the death of cells in a cancer model, we can interpret that the control activating the gene corresponding to component* **x**_2_ *may effectively eliminate the cancer in the long-term*.

*If C is applied as a release control maintained for a single transition, every state reaches a subspace* ∗1∗ *(colored in orange), and then the Boolean network reverts to its original form. For synchronous update mode, the transition from this subspace no longer reaches the attractor* { 000}, *which can also be effective in eliminating states violating the phenotype*.

*Another control C*^*′*^ = {(**x**_2_ = 1), (**x**_3_ = 1)} *leads to the same target states as C, and thus is also valid for the universal target control problem for all types of target states. However, it is not minimal since C is a subset of C*^*′*^.

*Notably, the target states can be an empty set after applying the control, as for the fixed point case. We consider it valid since there is no state violating the phenotype*.

### 2.2 Solution coverage between control tools

Given a BN *f* and a phenotype *φ*, a control tool outputs a control set. For two control sets ℂ_1_ and ℂ_2_, we say ℂ_1_ *covers* ℂ_2_ if for every control in ℂ_1_, there exists a control in ℂ_2_ that is a subset of it. Formally, this boils down to defining a binary relation ⇝ between control sets:

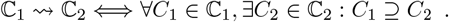

By convention, if ℂ_1_ = ∅, then ℂ_1_ ⇝ ℂ_2_ holds vacuously. Let us denote by 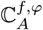 and 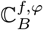 the control sets returned by tools *A* and *B* for the BN *f* and phenotype *φ*. We say that the tool *A* covers the tool *B* (*A* ⇝ *B*) if for any BN *f* and phenotype *φ*, 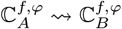.

While the theoretical coverage between tools may be characterized mathematically provided a formal characterization of their algorithm, an empirical coverage relationship between the implementations can be estimated over a fixed set of BNs and phenotypes.

Intuitively, when *A* ⇝ *B*, tool *A* can be viewed as more conservative than tool *B* as it always requires more simultaneous interventions to satisfy the same phenotype. Often *A* returns fewer controls than *B* since it is more difficult to satisfy the phenotype. This relationship can be represented as a *coverage graph*, where each node represents a tool and an arc *A* ⇝ *B* indicates that *A* covers *B*. There are several important properties of the coverage relation:

- **Transitivity**: If *A* ⇝ *B* and *B* ⇝ *C*, then *A* ⇝ *C*. Hence, we can omit transitive arcs (*A* ⇝ *C* in this case) for better visualization.
- **Minimality**: When comparing two control sets, it suffices to consider only their minimal controls. This property is useful since many tools only report minimal controls.
- **Equivalence**: If both *A* ⇝ *B* and *B* ⇝ *A* hold, then *A* and *B* have the same minimal control set for all given inputs. Therefore, those tools can be grouped into a single node, omitting the arcs among them.

**Example 3**. *Consider five control sets in Fig 2a from tools A, B, C, D, and E for a single input. The corresponding coverage graph for this restricted input is shown in Fig 2b, omitting all transitive arcs. First, the control* {(**x**_1_ = 1), (**x**_2_ = 0), (**x**_3_ = 1)} *of tool A is not minimal, and thus it can be ignored for determining the coverage. Although tool A finds more controls than B, they have the same ability to find minimal controls. Coverage relationships A* ⇝ *C and B* ⇝ *C can be easily checked by definition. Tools C and D do not cover one another because they fix the same component* **x**_1_ *to different values. Tool E returns* {∅} *as its only control (i*.*e*., *it fixes no component). Since* ∅ *is a subset of any control, tool E is covered by all other tools. Note that this example only illustrates the coverage based on a single input of BN and phenotype. In general, the coverage must be verified for all given inputs*.

**Fig 2.**
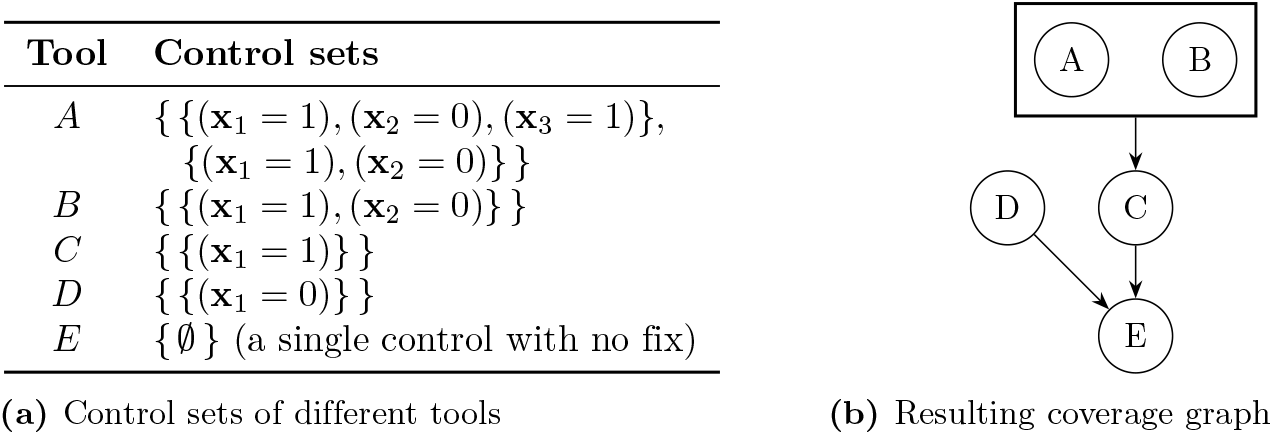
The coverage graph example for a single input

### 2.3 Taxonomy and theoretical coverage

Our taxonomy for classifying tools is based on two modeling assumptions for the universal target control problem: time span and type of target states. For time span, we consider permanent control (P) and release control (R). For target states, we consider five types: FP, SA, ASA, MTS, and VPTS. The set of target states is defined only when these two factors are given (see “S1 Appendix” for details).

Based on this taxonomy, we can theoretically determine the coverage relationships in Fig 3, where each node represents an exact tool for the corresponding universal target control problem. *Theoretical coverage* here means that the solution coverage holds for every input BN and phenotype, both synchronous and asynchronous update modes, and for any fixed release time. If no coverage exists from one setting to another, there is an input that contradicts the theoretical coverage. Therefore, Fig 3 completely characterizes the theoretical coverage relationships between all pairs of settings. All such findings are summarized in “S1 Appendix”. For instance, with a fixed time span, the exact tools VPTS cover those for SA, ASA, and MTS. However, there are no theoretical coverage relationships between permanent and release controls.

**Fig 3.**
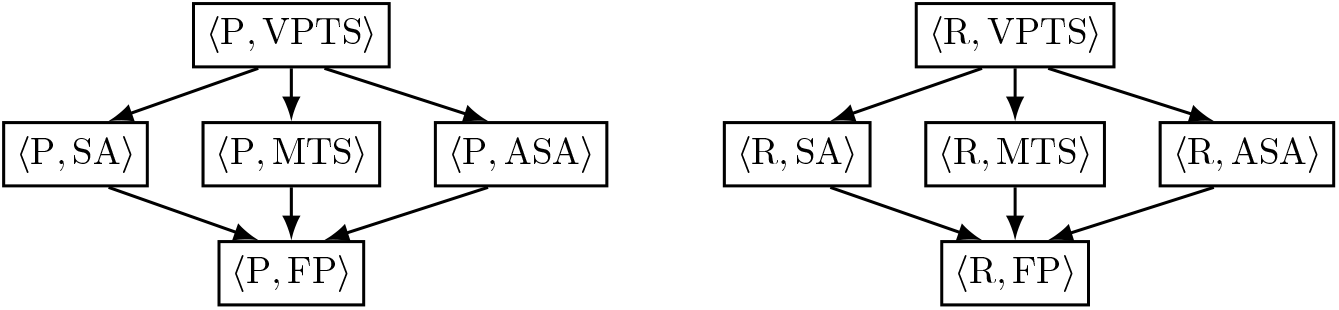
Theoretical coverage based on the type of target states and time span. Node labels are of the form ⟨Time span, Type of target states⟩. There is an arc from node A to B whenever A covers B. Transitive arcs are omitted.

Intuitively, the coverage relationships come from the inclusion relationships between target states. Given the same input, suppose there are two sets of target states with different types. By the definition of the universal target control problem, a control valid for a target set is valid for any other smaller target set. We observe the following inclusion relationships, coming from their formal definitions: VPTS target states always include the other four types of target states; FP target states are always included in the other four types of target states, as fixed points are a special case of attractors and trap spaces; there is no general inclusion between SA, ASA, and MTS target states. This leads to the coverage relationships in Fig 3 when the time span is fixed.

On the other hand, when the time span differs, no theoretical coverage relationship exists. This is because permanent and release control have different core abilities to change the target states, mostly coming from the fact that permanent control can drastically modify the attractors. Notably, permanent control can introduce new attractors while release control cannot, and it can partially remove some states from an existing attractor. In contrast, release control cannot modify attractors internally; it can only eliminate an attractor as a whole. Finally, release control can remove some transient states from target states, while permanent control cannot.

The notion of coverage relationships can clearly explain subtle differences in control sets from various tools, which has never been understood in previous works. Although control sets of tools might look completely different, they can be compared rigorously based on the coverage relationships derived from their modeling assumptions.

### 2.4 Mutation co-occurrence score for phenotype influence

In applications, the control of BNs is employed to uncover important genes whose mutation, possibly in synergy with the mutation of other genes, shows a strong influence on the studied phenotype.

In order to empirically compare and combine control sets of different tools, we propose the *Mutation Co-occurrence Score* (MCS) of a mutation *µ* in a given control set ℂ to assess its importance for the given phenotype. Denoted by MCS(*µ*, ℂ) where *µ* = {(**x**_*i*_ = *b*)} is a mutation of a single component, it relates the size of controls that include *µ*. Intuitively, a higher co-occurrence score indicates that a mutation is more likely to be sufficient to achieve the control objective on its own. On the other hand, a mutation which always has to be coupled with many other mutations has a low co-occurrence score. Thus, given two control sets, one covered by the other, the same mutation will have higher co-occurrence score in the covered one.

Our score function sums up the contribution from each minimal control. If *µ* itself is an element of the control set as a minimal control, the score is 1 as it is covered by all controls containing the mutation. However, if a minimal control *C* requires more co-occurring mutations for achieving the phenotype, its contribution to the score *w*(*C*) should be smaller. Hence, we recursively compute the score function *w*(*C*) as follows. Suppose there are two controls *C*_1_ and *C*_2_, both include the mutation *µ*, and they only differ by one element: |*C*_2_ \ *C*_1_| = 1. If a tool only reports minimal controls, then it should have discarded *C*_2_ since *C*_1_ is already sufficient to satisfy the phenotype. Therefore, a reasonable approach is to define the score of *C*_1_ as the sum of the scores of all such supersets *C*_2_:

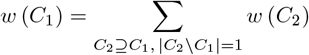

With equal weights, each of these supersets contributes 1*/*(2(*n* − |*C*_1_|)) to the score of *C*_1_ since there are *n* −|*C*_1_| possible components that can be additionally fixed, to either 0 or 1. This leads to a recursive definition

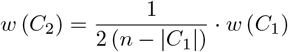

Solving this recursion with the base case *w*(*µ*) = 1, we obtain the score function for a single control in Equation (2). Before summing up the contributions, two cases need to be handled. Although a control *C* does not fix component **x**_*i*_, it still contributes to the score because *C* is covered by all controls in {*C*^*′*^ ∈ {0, 1, ∗}^*n*^ : (*C* ∪*µ*) ⊆*C*^*′*^}. Hence, we append *µ* to all such controls and compute the score in the same way as above. On the other hand, if a control *C* includes the contradictory mutation {(**x**_*i*_ = 1 −*b*)}, it should not contribute to the score since it cannot be covered by any control including the mutation *µ*. To handle these two cases, we use an augmented control set

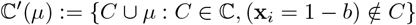

and recursively compute the score function as in Definition 2.

#### Definition 2

(Mutation co-occurrence score). *For each control C* ∈ ℂ^*′*^(*µ*) *in the augmented control set, its contribution to the mutation co-occurrence score is defined as*

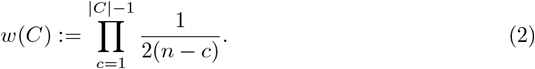

*The score for the entire control set is the sum of the score contributed by all minimal controls in the augmented set:*

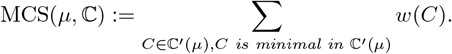

The mutation co-occurrence score has the following properties:

- **Normalization**: The score belongs to the range from 0 to 1. The maximum score is attained when ℂ contains ∅ or *µ*.
- **Additivity**: The score of a singleton control set ℂ = {*C*} is the sum of the scores of all controls that append one additional mutation to *C*.
- **Coverage**: Given two control sets ℂ_1_ and ℂ_2_, if ℂ_1_ ⇝ ℂ_2_, then MCS(*µ*, ℂ_1_) ≤ MCS(*µ*, ℂ_2_) for any mutation *µ*.

**Example 4** (Mutation co-occurrence score). *Consider a Boolean network with four components* {1, 2, 3, 4}. *Let the control set be* ℂ = {*C*_1_, *C*_2_, *C*_3_, *C*_4_, *C*_5_} *where*

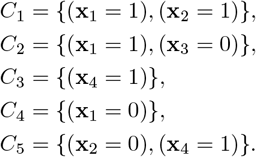

*Notably, C*_5_ *never contributes to any mutation score (it is always removed in the augmented-minimal filtering), but we keep it in the table for illustration*.

*The mutation* {(**x**_1_ = 1)} *appears in C*_1_ *and C*_2_, *and implicitly participates in C*_3_ *as* {(**x**_1_ = 1), (**x**_4_ = 1)} *after appending the mutation. C*_4_ *contradicts the mutation and thus it is ignored. Since all three participating controls are of size* 2,

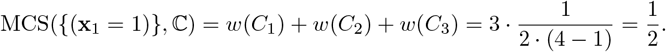

*The mutation* {(**x**_2_ = 1)} *appears in C*_1_. *After appending the mutation, it implicitly participates in C*_2_, *C*_3_, *and C*_4_ *as* {(**x**_1_ = 1), (**x**_3_ = 0), (**x**_2_ = 1)}, {(**x**_4_ = 1), (**x**_2_ = 1)}, *and* {(**x**_1_ = 0), (**x**_2_ = 1)} *respectively. However, C*_1_ *is covered by C*_2_ *after appending the mutation, and thus C*_2_ *is ignored. Since all three participating controls are of size* 2,

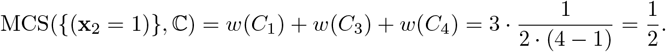

*The mutation* {(**x**_2_ = 0)} *has one contradicting control (C*_1_*), one contributing control with size 3 (C*_2_ *after appending the mutation), and two contributing controls with size 2 (C*_3_ *and C*_4_ *after appending the mutation). Therefore*,

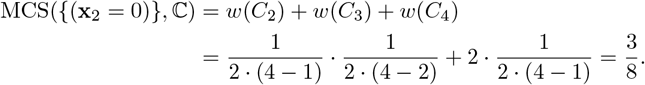

## 3 Results

### 3.1 The survey of software tools

Table 2 shows the list of tools reviewed in this survey. They are all distributed as part of the CoLoMoTo Docker distribution (up-to-date version: 2026-03-01) [19]. The column ‘Package name’ denotes the name of the package, which may support multiple control settings with various options in column ‘Option’. Abbreviations for the option are either taken from the original authors, or can be easily inferred from the reference. To denote a tool with a specific option, we use the notation ‘`Package name[Option]`’ (e.g., `PyBoolNet[ASA]`). For each tool, we indicated the closest settings following the taxonomy in Section 2.3. The meanings of symbols are explained in the caption. Brief information on each tool is summarized as follows.

**Table 1.**
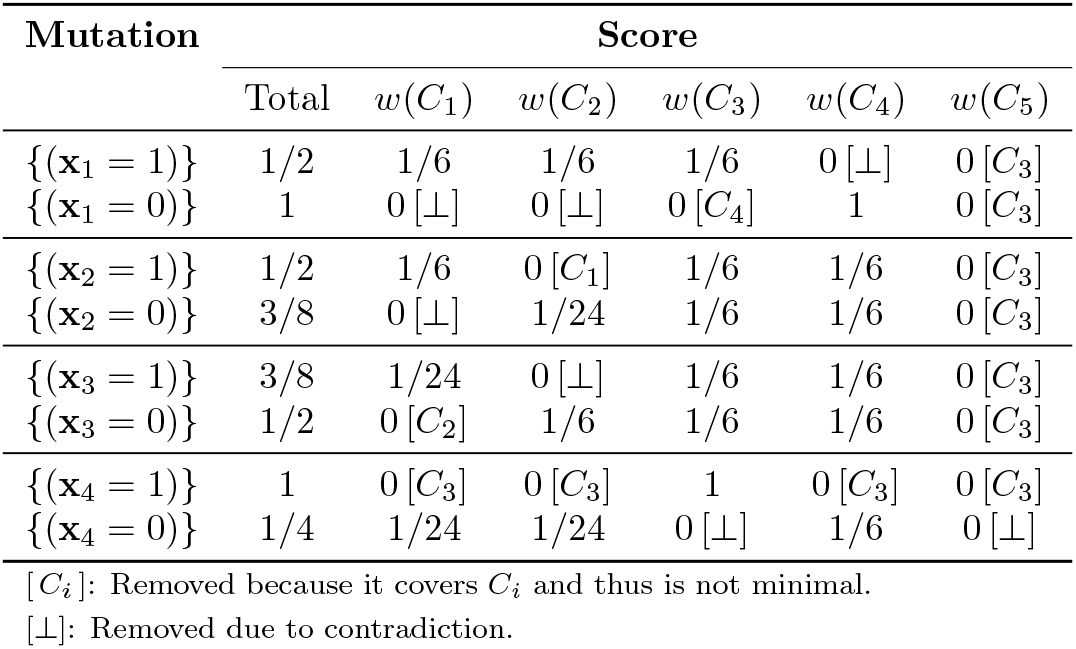
Mutation co-occurrence scores for Example 4.

| Mutation | Score |  |  |  |  |  |
| --- | --- | --- | --- | --- | --- | --- |
| | Total | $w(C_1)$ | $w(C_2)$ | $w(C_3)$ | $w(C_4)$ | $w(C_5)$ |
| $\{(\mathbf{x}_1 = 1)\}$ | 1/2 | 1/6 | 1/6 | 1/6 | 0 $[\perp]$ | 0 $[C_3]$ |
| $\{(\mathbf{x}_1 = 0)\}$ | 1 | 0 $[\perp]$ | 0 $[\perp]$ | 0 $[C_4]$ | 1 | 0 $[C_3]$ |
| $\{(\mathbf{x}_2 = 1)\}$ | 1/2 | 1/6 | 0 $[C_1]$ | 1/6 | 1/6 | 0 $[C_3]$ |
| $\{(\mathbf{x}_2 = 0)\}$ | 3/8 | 0 $[\perp]$ | 1/24 | 1/6 | 1/6 | 0 $[C_3]$ |
| $\{(\mathbf{x}_3 = 1)\}$ | 3/8 | 1/24 | 0 $[\perp]$ | 1/6 | 1/6 | 0 $[C_3]$ |
| $\{(\mathbf{x}_3 = 0)\}$ | 1/2 | 0 $[C_2]$ | 1/6 | 1/6 | 1/6 | 0 $[C_3]$ |
| $\{(\mathbf{x}_4 = 1)\}$ | 1 | 0 $[C_3]$ | 0 $[C_3]$ | 1 | 0 $[C_3]$ | 0 $[C_3]$ |
| $\{(\mathbf{x}_4 = 0)\}$ | 1/4 | 1/24 | 1/24 | 0 $[\perp]$ | 1/6 | 0 $[\perp]$ |

$[C_i]$ : Removed because it covers $C_i$ and thus is not minimal.
$[\perp]$ : Removed due to contradiction.

**Table 2.** The list of software tools classified according to universal target control problems.

| Software tools |  |  | Time span | Type of target states |  |  |  |  |
| --- | --- | --- | --- | --- | --- | --- | --- | --- |
| Package name | Option | Ref. |  | FP | SA | ASA | MTS | VPTS |
| ActoNet | - | [12] | P | ■ |  |  |  |  |
| optboolnet<br>(optbn) | FP | [14] | P | ■ |  |  |  |  |
|  | SA | [24] | P | □ | ■ |  |  |  |
| BoNesis | FP | [15] | P | ■ |  |  |  |  |
|  | MTS | [22] | P | □ |  |  | ■ |  |
| PyBoolNet<br>(PBN) | SA | [13] | P | □ | ■ |  |  |  |
|  | ASA | [13] | P | □ |  | ■ |  |  |
|  | percolation | [25] | P |  |  |  |  | △ |
|  | trap_spaces | [26] | P |  |  |  |  | △ |
| Caspo | - | [18] | P | □ | □ | □ | □ | ■ |
| CABEAN | ITC | [17] | R |  |  | △ |  |  |
|  | TTC | [17] | R |  |  | △ |  |  |
|  | PTC | [17] | R&P |  |  | △ |  |  |
| pystablemotifs<br>(SM) | brute-force | [16] | R&P |  |  |  |  | △ |
|  | internal | [16] | R&P |  |  |  |  | △ |
|  | minimal | [16] | R&P |  |  |  |  | △ |

■ (Exactness): Enumerates all minimal controls satisfying the phenotype on all target states.
□ (Validity): Guarantees that the phenotype is satisfied on all target states.
△ (Heuristic): Conceptually employs target states with no underlying universal target control problem.
R&P: Uses both BNs before and after release control, not perfectly fitting into the taxonomy.

- `ActoNet` (ver. 1.0): One of the earliest tools by [12] that enumerates control sets for fixed points. Originally designed as a binary decision diagram approach, it was later implemented using logic programming with Answer Set Programming (ASP).
- `optboolnet` (ver. 1.0.1): A bilevel optimization approach for efficiently enumerating control sets with two options for target states `FP` [14] and `SA` [24]. The implemented version for the FP control is slightly different from the original paper [14] since it follows a more recent approach by [24].
- `BoNesis` (ver. 0.6.9): logic programming (ASP) approach for the synthesis (inference) of BNs from prior knowledge and constraints on the dynamics [27]. As the synthesis problem generalizes the control problem, it provides two options for target states: `FP` and `MTS` [15, 22].
- `PyBoolNet` (ver. 3.0.16): A Python package providing multiple analysis tools for BNs. Options for `SA` and `ASA` are the implementation of a model checking approach by [13]. The option `percolation` is a preliminary version by [25] that employs a VPTS-based heuristic approach. The option `trap_spaces` is another heuristic based on ASP by [26].
- `Caspo` (ver. 4.0.1): A Python package for construction, analysis, and control of BNs based on VPTS approaches [18]. The core idea has been advanced by [28–30].
- `CABEAN` (ver. 2.0.1, binary file): A software collection for solving release control problems [17, 31]. Supports several target control options based on the release time and method to filter attractors: Instantaneous (`ITC`), Temporary (`TTC`), and Permanent (`PTC`) [32, 33].
- `pystablemotifs` (ver. 3.0.6): A Python package for finding control sets based on special types of trap spaces called *stable motifs* [34]. The core idea was proposed by [35], further developed by [16]. `pystablemotifs` provides options for control problems, including an exact method `brute-force` along with several heuristics. Among heuristic options represented as a tuple ‘(target method, driver method)’, we selected two options ‘(merge, internal)’ and ‘(merge, minimal)’, denoted as `internal` and `minimal`, respectively.

Table 2 shows that most packages provide at least one exact approach to specific universal target control problems, often with heuristics for computational efficiency. For example, `PBN` aims to control SA or ASA while providing VPTS-based heuristics: `percolation` and `trap_spaces`. These approaches can be justified because SA and ASA are covered by VPTS. Another finding is that most tools focus on permanent control because it is easier to handle than the release control. An important exception is `CABEAN`: both implications above fail because it explicitly removes ASAs that are not reachable from given source attractors. As the control problem can be formally described as a logic programming problem, answer set programming and formal methods are the most popular approaches used by `ActoNet`, `BoNesis`, `PyBoolNet`, and `Caspo`.

### 3.2 Coverage between selected control tools

We compare the tools on Table 2 using coverage testing to identify counterexamples under a selected set of input and output settings: On the input side, we selected eleven BNs listed in Table 3. Three BNs A1-A3 are from biological case studies collected from the online repository of `GINsim` [36]. Proper phenotypes were defined based on the original papers. The other eight BNs B1-B8 are manually designed small BNs to show counterexamples. For B6-B8, we added an auxiliary component *PHENOTYPE* to BNs in [37] with *f*_*PHENOTYPE*_ representing the original phenotype. The new phenotype is *φ*(**x**) = **x**_*PHENOTYPE*_. On the output side, we enforced an option to limit the number of fixed components in a control to be two or less. All tools support this option to speed up computation, except `CABEAN` and `PBN`. To apply this constraint across tools, we applied a post-processing step to filter out controls with more than two fixed components. As some tools do not allow fixing components involved in the phenotype, we also removed such controls in post-processing. For the bladder cancer (A1), we initially fixed the following three components and applied value propagation: *GrowthInhibitors* = 1, *EGFR stimulus* = 1, *FGFR3 stimulus* = 1.

**Table 3.** Characteristics of each problem input. Group A: Biological case studies from the GINsim repository [36]. Group B: Manually designed small BNs to show counterexamples.

| No. | Size | Remarks |
| --- | --- | --- |
| A1 | 35 | Bladder cancer model by [11] |
| A2 | 60 | T-LGL model by [8] |
| A3 | 32 | Tumor invasion model by [10] |
| B1 | 4 | One long ASA spanning all states (no FP exists) |
| B2 | 4 | $\langle P, ASA \rangle \not\rightsquigarrow \langle P, MTS \rangle$ |
| B3 | 4 | $CABEAN[TTC] \not\rightsquigarrow CABEAN[ITC]$ |
| B4 | 3 | $\langle P, ASA \rangle \not\rightsquigarrow \langle R, ASA \rangle$ |
| B5 | 3 | $\langle R, ASA \rangle \not\rightsquigarrow \langle P, ASA \rangle$ |
| B6 | 4 | Fig 4a in [37]: $\langle P, ASA \rangle \not\rightsquigarrow \langle P, SA \rangle$, $\langle P, ASA \rangle \not\rightsquigarrow \langle P, MTS \rangle$ |
| B7 | 4 | Fig 4b in [37]: $\langle P, SA \rangle \not\rightsquigarrow \langle P, ASA \rangle$, $\langle P, SA \rangle \not\rightsquigarrow \langle P, MTS \rangle$ |
| B8 | 4 | Fig 4c in [37]: $\langle P, MTS \rangle \not\rightsquigarrow \langle P, SA \rangle$, $\langle P, MTS \rangle \not\rightsquigarrow \langle P, ASA \rangle$ |

All experiments were done using CoLoMoTo Docker version 2025-12-01 on a Windows™ computer equipped with Intel™ CPU (i7-14700KF) and 64GB RAM. See code availability section and “S2 File” for details. `CABEAN` terminated with errors in a few instances with no control set returned. In those cases, we mapped it to a control set following the definition of the universal target control problem: (i) *Memory limit exceeded*. We treated the output as an empty control set ℂ =∅ (A3, all options); (ii) *Only one attractor*. If all states in the unique attractor satisfy the phenotype, we accepted ℂ = {∅} as a valid control set; otherwise we set ℂ = ∅ (B1, B2, B6, B7 for `TTC` and `PTC`); (iii) *No attractor within the phenotype*. We set ℂ = ∅ (B1, B4, B7 for `ITC`; B4 for `TTC` and `PTC`). These exceptions did not affect the coverage graph since other counterexamples were found in the same settings. Notably, optional parameters may affect the results. We used the default settings if possible, except for the following:

- `ActoNet`: ‘existential’=False (to allow the empty set as target states);
- `optbn[SA]`: ‘max_attractor length’=45 (large enough to capture all attractors);
- `BoNesis[MTS]`: ‘at_least_one’=False (to allow the empty set as target states);
- `PBN[percolation]` and `PBN[trap_spaces]`: ‘intervention type’=‘node’;
- `CABEAN`: ‘compositional’=2, provided ‘-tmarker’ option for target control.

Fig 4 shows the coverage graph and counterexamples identified with the eleven inputs in Table 3. Fig 4a omits transitive arcs for visibility. The tools are colored based on their time spans. Equivalent tools are grouped as labeled clusters. The matrix in Fig 4b shows the first counterexample input in Table 3 for the corresponding coverage comparison. The matrix with the full set of counterexamples is available in “S2 File”.

**Fig 4.**
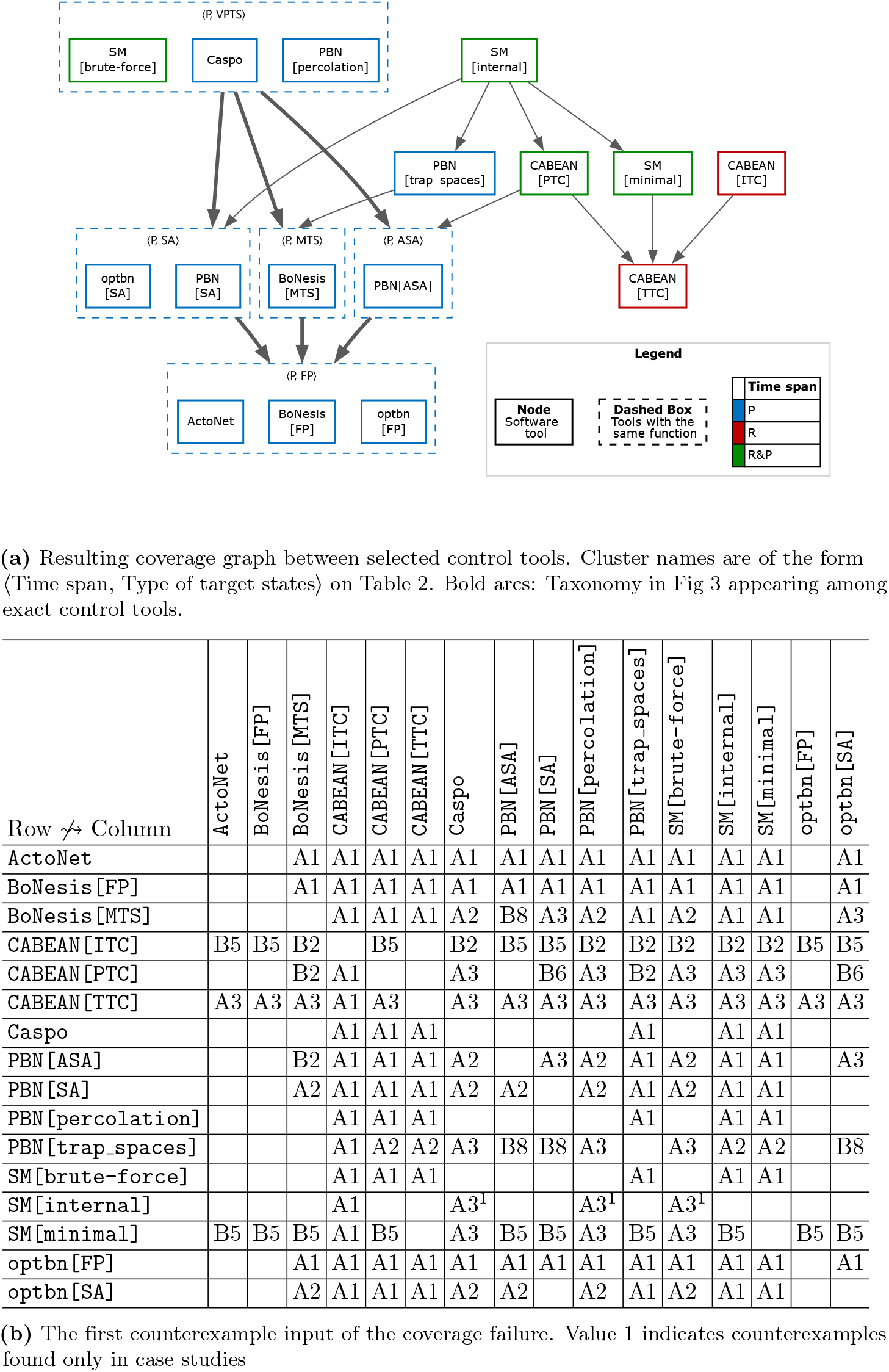
Empirical coverage between selected control tools

The coverage graph completely separates permanent control tools from release control tools as expected. Tools with R&P have arcs to both types of tools, especially with `CABEAN[PTC]` that bridges them with direct arcs. The perment control tools are mostly exact, and thus the six bold arcs are consistent with theoretical arcs in Fig 3. Based on these observations, several conjectures on coverage relationships were derived for future theoretical investigations:

1. Will the tools with R&P be separated from permanent or release control tools?
2. Is the empirical coverage between `pystablemotifs` and `CABEAN` actually general? If so, what are the underlying theoretical reasons?
3. Will the three tools with VPTS as target states (`PBN[percolation]`, `SM[brute-force]`, and `Caspo`) remain equivalent in general? If so, can we prove their equivalence theoretically?

The counterexample matrix Fig 4b highlights several implications for testing new tools and inputs. First, inputs from biological case studies may not capture all counterexamples, especially for `CABEAN[ITC]` and `SM[minimal]`. Even when a counterexample is found among case studies, a smaller manually designed counterexample is often available. Hence, a new tool should be tested on small inputs first for computational efficiency. The only exception is `SM[internal]`, for which A3 is the only counterexample observed.

### 3.3 Mutation co-occurrence score for T-LGL model

To illustrate the use of the mutation co-occurrence score, we analyzed the T-LGL leukemia model by [8] (A2 in Table 3) using 16 different tools listed in Table 2. The T-LGL model has 60 components and models the survival signaling network of T-LGL leukemia cells. The phenotype is *φ*(**x**) := **x**_*Apoptosis*_, which is an auxiliary component representing the apoptotic death of T-LGL leukemia cells. Hence, each computed control is intended to satisfy apoptosis on all target states (under the control). The full data are available in “S3 File”.

Fig 5 illustrates various patterns of the MCS scores for three representative components: *Caspase, CD45*, and *S1P*. Upward blue bars and downward red bars represent scores for the mutations that fix components to 1 and 0, respectively. Fixing to 1 and 0 correspond to activation and inhibition, respectively. The last six tools from `SM[internal]` to `CABEAN[ITC]` predicted near-zero scores for all components. Therefore, we focus on the first ten tools from `ActoNet` to `PBN[percolation]`. For *Caspase*, most tools yield similar scores overall. On the other hand, for *CD45*, the predicted scores vary: the tools with target states FP predicted high scores for both activation and inhibition, while the others did not. For *S1P*, predicted scores for both activation and inhibition are high. The scores for inhibition are slightly higher than those for activation, but the difference is not as significant. This suggests that a component may influence phenotype satisfaction under both activation and inhibition.

**Fig 5.**
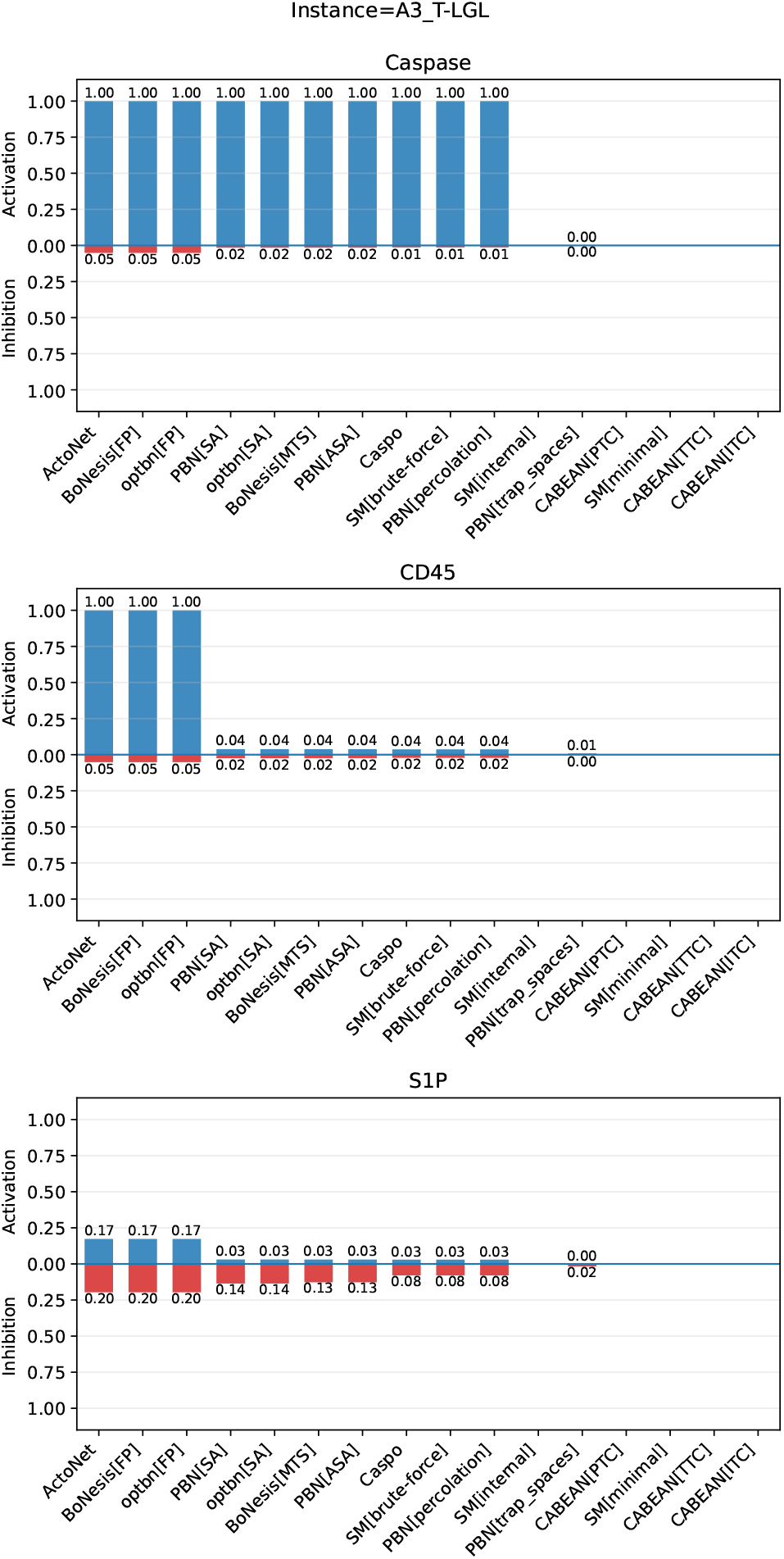
MCS scores for three representative components in the T-LGL model

We computed the MCS scores for the BN proposed by [8] for T-LGL leukemia by averaging the scores from different groups of tools based on their target states and time span settings. Table 4 lists the top components ranked by the average MCS scores across all 16 tools on Table 2. We only evaluated the scores for inhibition as it is often more practical and relevant in therapeutics. Each column corresponds to a specific group of tools, indicating ⟨Time span, Type of target states⟩ on Table 2. If multiple tools match the column, their MCS scores are averaged, and the ranks are determined based on the averaged scores. The table only shows the top 16 components; the remaining components have MCS scores significantly lower than those listed as the row for “Not listed” indicates.

**Table 4.**
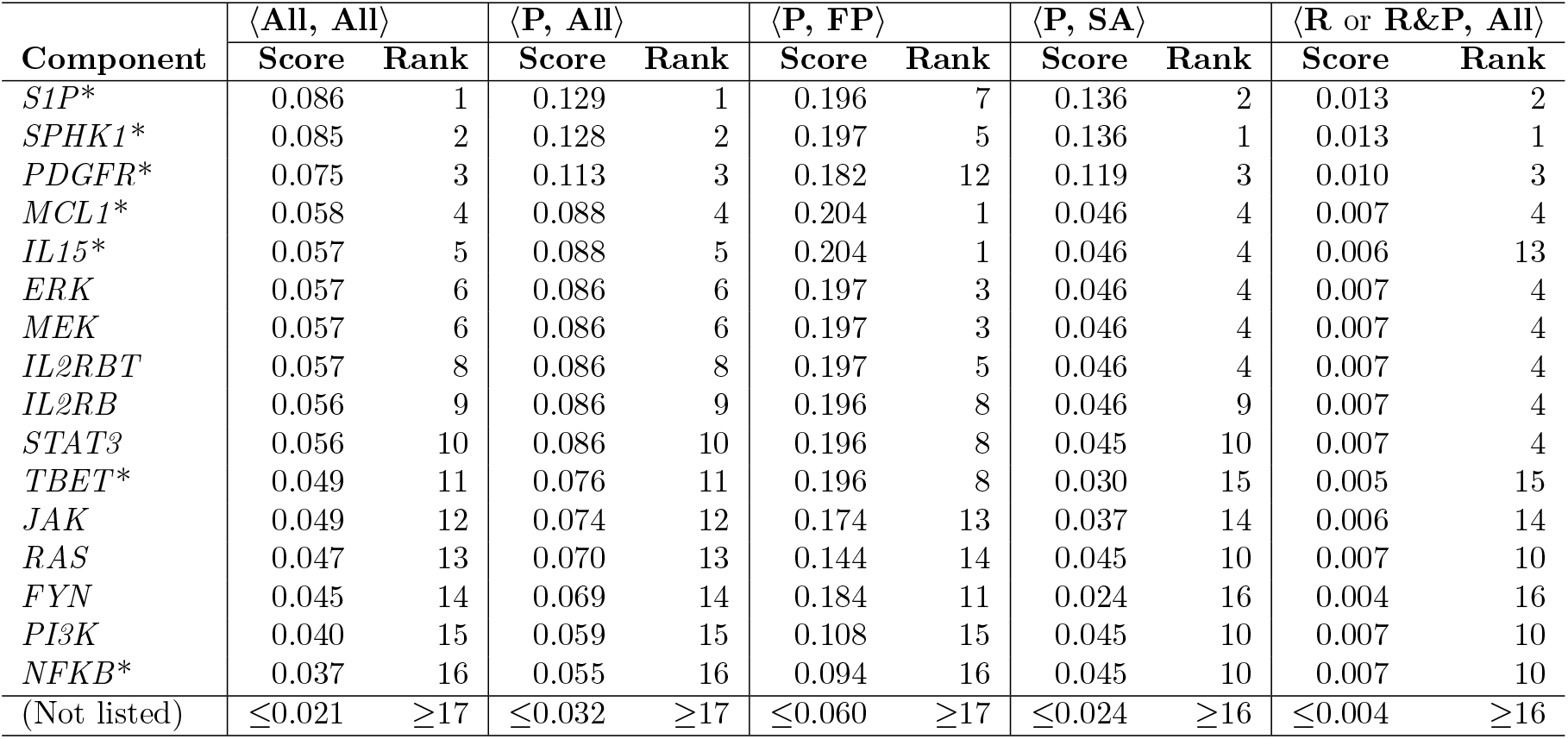
Components with top MCS scores for the apoptotic phenotype via inhibition. The first row indicates ⟨Time span, Type of target states⟩ on Table 2 for classifying tools. If multiple tools match the column, their MCS scores are averaged. Asterisks (*) indicate 7 key components (out of 60) identified by [7] as critical for apoptosis.

The ranks from all 16 tools ⟨All, All⟩ are highly consistent with those from permanent control tools ⟨P, All⟩ as they recorded higher MCS scores overall. However, if we select a specific target state, some of the components may be overlooked. For example, if we only considered FP as target states (i.e., ⟨P, FP⟩), *PDGFR* would be underrated from rank 3 to rank 12. Similarly, if we only considered SA as target states (i.e., ⟨P, SA⟩), *IL15* would drop from rank 5 to rank 13. By including more tools, we can reduce the risk of missing critical components. Moreover, fewer components have tie scores when averaging across tools, making it easier to prioritize components for experimental validation. For release control tools (i.e., ⟨R or R&P, All⟩), the MCS scores are generally lower, and thus the ranks are less reliable.

We observe that most key components identified by [7] as critical for apoptosis are also highly ranked in our analysis. The BN we used was a version slightly modified in [8] from the original in [7], but the difference is insignificant. Our top-ranked inhibition targets in Table 4 align closely with the key survival mechanisms highlighted by [7]. The sphingolipid pathway was predicted by [7] as essential for long-term survival, specifically *SPHK1* (sphingosine kinase 1) and *S1P* (sphingosine-1-phosphate). These two components have top MCS ranks, and thus inhibiting them may be an effective route to enforce the apoptosis phenotype. [7] also identified persistent *IL15* (interleukin 15) and *PDGF* (platelet-derived growth factor) signaling as sufficient to reproduce the leukemic survival phenotype. They reported that disrupting *PDGF* signaling induced apoptosis. Correspondingly, *PDGFR* (platelet-derived growth factor receptor) recorded a high MCS score for apoptosis by inhibition, consistent with *PDGF* /*PDGFR* acting as an upstream survival driver. Downstream, [7] emphasized the central role of *NFKB* (NF-*κ*B; nuclear factor kappa B) in sustaining survival via maintaining expression of *MCL1* (myeloid cell leukemia 1), with *NFKB* functioning downstream of *PI3K* (phosphoinositide 3-kinase). This result is consistent with the fact that *NFKB, MCL1*, and *PI3K* are among our high-scoring inhibition targets. Finally, [7] predicted that *TBET* (T-bet; T-box 21) activation must accompany *NFKB* activation to reproduce the leukemic phenotype. If *TBET* activation is required for survival, inhibiting *TBET* should disrupt survival, which is consistent with its high MCS score.

For other two BN models in Table 3, we also found that the top-ranked components were also highlighted as critical in the original papers (full MCS scores are presented in “S3 File”). [11] identified co-occurrence of *FGFR3* activation with activation of *RAS* or *PIK3CA* as a critical driver of bladder cancer progression. Our analysis shows that inhibition of *RAS, FGFR3*, and *PIK3CA* all have high MCS scores (rank 1, 2, 4) for the apoptotic phenotype (*φ*(**x**) = *Apoptosis b1* ∧ *RB1*), consistent with their findings. In the tumor invasion model (A3, 32 components) [10], *p53* and *SNAI1* are ranked 2 and 3 for inhibiting the metastatic phenotype. This is consistent with their finding that *p53* activity strongly reduces metastasis, especially for *SNAI1* loss of function mutants. Moreover, [10] identifies *SNAI2* and *AKT2* gain-of-function as metastasis amplifiers, matching their high activation MCS ranks in our analysis (1 and 4, respectively).

## 4 Discussion

### 4.1 Extension of the proposed taxonomy

The proposed taxonomy only deals with universal target control problems, but it can be generalized in several ways. The first extension is an *existential target control problem* [12] where a control is valid if at least one of the target states satisfies the phenotype:

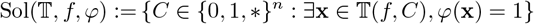

If all other settings remain, every solution for a universal target control problem is also a solution for an existential target control problem if target states are not empty (e.g., SA, MTS, ASA, and VPTS). Therefore, a control set for an existential target control problem must be covered by a universal target control problem for such cases. For the existential variant, the phenotype is easier to satisfy for a larger set of target states, which is the opposite of the universal target control problem. Hence, the theoretical coverage relationships in Section 2.3 hold with the arcs reversed.

The second extension is the *source-target control problem* [17] where a subset of target states reachable from a given set of source states 𝕊 in the STG is considered when evaluating the phenotype:

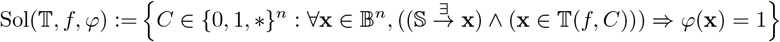

The universal target control problem is a special case of this with 𝕊 = 𝔹^*n*^ and ignoring the reachability condition 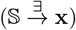. For the source-target variant, we can expand the taxonomy according to the inclusion relationship of the source states: the phenotype is easier to satisfy for smaller source states. Source-target control problems have a close connection to the release control because the set of states visited at release can be regarded as the source states [31].

Last, the current definition of control fixing the value of a component can be relaxed as *edgetic control* [38] that selectively fixes components in particular Boolean functions, providing more flexibility in control design. For instance, if an edge from component *i* to component *j* is fixed to 0, the Boolean function of component *j* is modified by substituting *x*_*i*_ with 0 (i.e., 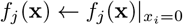. Other terms in *f*_*j*_ are not affected, which is the core difference from the current definition of control. Edgetic control can be interpreted as an intervention on interactions among biological entities rather than on the entities themselves.

### 4.2 Discovering synthetic lethal or viable pairs via mutation co-occurrence score

Currently, the MCS score is defined for a singleton mutation. It can be naturally extended to any control that fixes multiple components. For control *µ* ∈ {0, 1, ∗} ^*n*^ to evaluate the score, we can redefine the augmented control set by adding *µ* to every control in the original control set while ensuring no conflicting mutations exist:

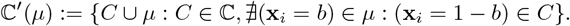

Then, the contribution of each control *C* ∈ ℂ′(*µ*) to the MCS score can be redefined as

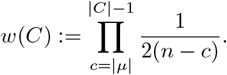

This leads to the same score function Definition 2 with all properties preserved. One application in cancer research is to discover synthetic lethal or synthetic viable pairs, which must be simultaneously controlled to achieve apoptotic or cancerous phenotypes of cells [39–41]. By comparing the MCS scores among all pairs of components, we can prioritize candidate pairs for experimental validation. Currently, we take the average of MCS scores from different tools, but more sophisticated aggregation methods can be considered for reliable scoring. For example, different weights may be provided to tools based on the user’s prior knowledge.

### 4.3 Computational complexity analysis

Recently, [42] reviewed the computational complexity results for universal target control problems, providing theoretical insights into the practical performance of each tool. They focused on verification problems (given a control, checking if phenotype is satisfied at all target states) and search problems (finding a single control). The verification and search problems of permanent control for FP are coNP-complete and 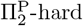, respectively [14, 43]. The verification and search problems of permanent control for MTS are 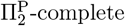 [44] and 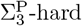, respectively [42]. There are no known results for SA and ASA, but they are conjectured to be PSPACE-hard as checking whether a state belongs to an attractor is PSPACE-complete [45]. For SA with the release control, [46, 47] showed that the search problem can be PSPACE-hard, assuming that the initial and the final states are given and the control can be applied more than once during transitions. The results for other settings are still limited, and they are left as important future work.

### 4.4 Future work

Several tools outside the CoLoMoTo Docker are omitted due to integration difficulty, but they are worth mentioning. `AEON.py` [48] provides both methods for permanent and release control. `AEON.py` can even handle partially specified BNs where several candidate Boolean functions are available [49, 50]. [51] proposed an algebraic approach and network reduction method for permanent control of SAs. These tools would be the next candidates to be added to the coverage graph. Our framework is limited to controls applied only once, while some tools compute a sequence of controls applied at different time steps [17, 46, 47, 52]. Such tools cannot be directly compared using our framework, which remains an open challenge.

Another interesting future work is finding specific types of BNs for which theoretical coverage holds in addition to Fig 3. For example, [53] discovered a family of BNs where MTS ⇝ ASA holds. For such BNs, more efficient approaches can be used.

## 5 Conclusion

Boolean network control is often presented as a single task, but current software tools have different assumptions and capabilities. They vary in which target states are required to satisfy the phenotype and in how long the control is applied. In this paper, we made these assumptions explicit as universal target control problems, using them as the core framework to compare existing tools.

Our main technical contribution is a solution-coverage framework that supports tool-to-tool comparison even when outputs are filtered by minimality. We provided an open-source implementation for empirical coverage testing on benchmark inputs. Building on this, we proposed a taxonomy of universal target control problems that explains a large portion of the coverage relationships observed in practice. Finally, we introduced the mutation co-occurrence score to quantify how strongly single-component perturbations influence a phenotype. If multiple tools are available, the score can be aggregated across tools to provide robust predictions. The validation on the T-LGL leukemia model confirmed previously known biological insights for apoptosis-related targets.

Looking forward, our taxonomy naturally extends beyond universal target control problems, including existential control, source-target control, and partially specified Boolean networks. We expect that making problem assumptions explicit, and validating them through coverage-based comparisons, will help researchers choose appropriate tools, reconcile conflicting predictions, and report control results with clearer biological interpretation.

## Software and data availability

All code and data underlying the findings reported in this manuscript are publicly available without restriction. The full analysis pipeline (scripts to run tool comparisons, compute coverage relationships, and reproduce all figures and tables) and the minimal dataset needed to replicate the reported results are available in the project GitHub repository (https://github.com/kalebmoon07/BN-control-taxonomy) and are permanently archived in the Software Heritage archive (https://archive.softwareheritage.org/swh:1:dir:9af25fb7d8e04509c59b091d8806af389c24bcd0).

In addition, the Supporting Information includes all derived results: “S2 File” provides benchmark control sets and the full list of counterexamples (including counterexample full match.csv), and “S3 File” provides the full data and figures for the mutation co-occurrence score analysis on the T-LGL leukemia model.

## Supporting information

**S1 Appendix**. Formal definitions and proofs. This appendix provides formal definitions required to define the universal target control problems and the proofs for the theoretical coverage relationships in Section 2.3.

**S2 File**. Results for solution coverage analysis. This file provides the control sets of each software tool on benchmark inputs. The full list of counterexamples identified in the solution coverage analysis in Section 3.2 can be found in “counterexample full match.csv”.

**S3 File**. Results for mutation co-occurrence score analysis. This file provides the full data and figures for the mutation co-occurrence score analysis on the T-LGL leukemia model in Section 3.3. The results for bladder cancer and tumor invasion in Table 3 are also included.

## Acknowledgments

KL and KM acknowledge funding from the National Research Foundation of Korea (NRF), the Ministry of Science and ICT (MSIT), and the Ministry of Education (MOE). This work was supported by the National Research Foundation of Korea (NRF) grant funded by the Korean government (MSIT) (No. NRF-2022R1F1A107414011), supported under the framework of international cooperation program managed by National Research Foundation of Korea (grant number: RS-2023-00259481), and supported by Basic Science Research Program through the National Research Foundation of Korea (NRF) funded by the Ministry of Education (No. RS-2025-25433718).

CB and LP acknowledge funding from French Agence Nationale pour la Recherche (ANR) in the scope of the project RD2BOOL (ANR-23-CE45-0014), REPAIRNET (ANR-25-IABT-0003), and by CampusFrance in the scope of the PHC STAR 2024 project 50147NM.

The funders had no role in study design, data collection and analysis, decision to publish, or preparation of the manuscript.

