## Supplementary material for "Why Boolean network control tools disagree: a taxonomy of control problems": S1. Appendix

### S1 Appendix. Formal definitions and proofs in “Why Boolean network control tools disagree: a taxonomy of control problems”

Célia Biane[1], Kyungduk Moon[2*], Kangbok Lee[2], Loïc Paulevé[1]

**1** Univ. Bordeaux, CNRS, Bordeaux INP, LaBRI, UMR 5800 F-33400 Talence, France
**2** Pohang University of Science and Technology, Pohang, 37673, Korea

*

### 1 Formal definitions related to the universal target control problem

#### 1.1 Boolean networks and state transition graphs

We denote the set of Boolean values by $\mathbb{B} := \{0, 1\}$. A Boolean network on $n$ components is a map $f : \mathbb{B}^n \to \mathbb{B}^n$ and its *state* is a Boolean vector $\mathbf{x} \in \mathbb{B}^n$ with assigned values. A *phenotype formula* (simply *phenotype* hereafter) is a Boolean function $\varphi : \mathbb{B}^n \to \mathbb{B}$ that evaluates a given state $\mathbf{x} \in \mathbb{B}^n$ whether it satisfies the desired property or not. A *(Boolean) subspace* is specified by a vector $\mathbf{h} \in \{0, 1, *\}^n$. A component $i$ is *fixed* if $\mathbf{h}_i \in \mathbb{B}$ and *free* if $\mathbf{h}_i = *$. A subspace corresponds to the set of states where free components can take any Boolean values while fixed components remain constant.

$$s(\mathbf{h}) := \{\mathbf{x} \in \mathbb{B}^n : \forall i, (\mathbf{h}_i \in \mathbb{B}) \Rightarrow (\mathbf{x}_i = \mathbf{h}_i)\}.$$

If $\mathbf{x} \in s(\mathbf{h})$, we say $\mathbf{h}$ *contains* $\mathbf{x}$ and $\mathbf{x}$ *belongs to* $\mathbf{h}$.

A *state transition graph* (STG) is a directed graph $G(\mathbb{B}^n, E)$ where each node in $\mathbb{B}^n$ represents a state, and each arc $(\mathbf{x}_1, \mathbf{x}_2) \in E$ represents a *transition* from state $\mathbf{x}_1$ to $\mathbf{x}_2$. There are various *update modes* to specify how to construct the STG based on $f$. In this paper, we focus on two special types of update modes: *synchronous* and *fully asynchronous*. The *synchronous* update maps one state to the unique successor by simultaneously updating all components. In other words, $(\mathbf{x}, \mathbf{y}) \in E \iff ((\mathbf{y} = f(\mathbf{x})) \wedge (\mathbf{x} \neq \mathbf{y}))$. On the other hand, the *fully asynchronous* update maps each state to multiple successors where any single component is updated. In other words, $(\mathbf{x}, \mathbf{y}) \in E \iff \exists i \in [1, n] : (\mathbf{y}_i = f_i(\mathbf{x})) \wedge (\mathbf{x} \neq \mathbf{y}) \wedge (\forall j \neq i : \mathbf{y}_j = \mathbf{x}_j)$. In this study, we simply abbreviate the fully asynchronous update as *asynchronous*. Other update modes are summarized by [1]. A state $\mathbf{y}$ is *reachable* from another state $\mathbf{x}$ if there exists a directed path from $\mathbf{x}$ to $\mathbf{y}$ in $G$. If we just say a state $\mathbf{x}$ is reachable, it means it is reachable from some initial state in a set provided by the context.

#### 1.2 Control problem and biological interpretation

A *control* $C$ is a partial assignment of Boolean values to components given as a subset of $\{(\mathbf{x}_i = b) : i \in [1, n], b \in \mathbb{B}\}$. For each $i$, at most one assignment $(\mathbf{x}_i = b)$ can be included in $C$. Fixing a component $i$ to 1 or 0 represents a biological intervention that activates or inhibits the corresponding entity, respectively. The BN $f$ controlled by $C$ is denoted as $f/C$ and it must satisfy

$$(f/C)_i(\mathbf{x}) = \begin{cases} b & \text{if } (\mathbf{x}_i = b) \in C \\ f_i(\mathbf{x}) & \text{otherwise} \end{cases}.$$

A control can also be represented as a subspace $C \in \{0,1,*\}^n$ where control assignments indicate fixed components and the other components are free. Hence, the set of all controls can also be represented as $\{0,1,*\}^n$. A control may have different *time spans* to apply: *permanent* and *release*. The permanent control fixes components during all transitions. For the release control, components are fixed for a limited number of transitions. We are particularly interested in the reachable states after releasing the control. *Release time* is the number of transitions after which the BN reverts to its original form.

We have several notes for release control. The update mode and release time may differ by tools, and it affects the set of reachable states. Release control can change the set of reachable target states significantly. We assume that release time is a deterministic, positive number independent of the type of target states. We clarify under which conditions our proofs hold, if necessary. Otherwise, our proofs hold for any release time and both synchronous and asynchronous update modes.

##### 1.3 Five types of target states

*Target states* are the states that we desire to satisfy the phenotype after applying control. One common property for target states is that they are trap sets (see Definition 1); no further transition escapes the set once entered [2], restricting the states visited in future transitions. We focus on the following types of target states in Definition 2-5: *fixed points* (FP), *synchronous attractors* (SA), *asynchronous attractors* (ASA), *minimal trap spaces* (MTS), and *value-propagated trap spaces* (VPTS). We use $\delta$ to denote a type of target states, i.e., $\delta \in \{\text{FP}, \text{SA}, \text{ASA}, \text{MTS}, \text{VPTS}\}$. If an input BN is $f$, the corresponding set of target states can be denoted as $\mathcal{T}_f^\delta$. If we consider the BN under control $C$, the resulting set of target states becomes $\mathcal{T}_{f/C}^\delta$.

**Definition 1** (Trap set)**.** *Given a state transition graph $G(\mathbb{B}^n, E)$, a set of states $\mathcal{T}$ is a* trap set *if for each pair of states $\mathbf{x} \in \mathcal{T}$ and $\mathbf{y} \notin \mathcal{T}$, $(\mathbf{x},\mathbf{y}) \notin E$.*

**Definition 2** (Fixed point)**.** *A* fixed point *(FP) is a state $\mathbf{x} \in \mathbb{B}^n$ satisfying $f(\mathbf{x}) = \mathbf{x}$. We denote the set of all fixed points as $\mathcal{T}_f^{\text{FP}}$.*

**Definition 3** (Attractor)**.** *An attractor $\mathcal{A} \subseteq \mathbb{B}^n$ is a set of states with the following properties:*

1. ***Terminal**: For all $\mathbf{x} \in \mathcal{A}$, if there exists a transition $(\mathbf{x},\mathbf{y}) \in E$, then $\mathbf{y} \in \mathcal{A}$.*
2. ***Strongly connected**: All pairs of states $\mathbf{x},\mathbf{y} \in \mathcal{A}$ are mutually reachable (i.e., there exists a path in $G$ from $\mathbf{x}$ to $\mathbf{y}$ and vice versa).*

*Based on the update mode, $\mathcal{A}$ is either called a* synchronous attractor *(SA) or an* asynchronous attractor *(ASA). We denote the union of all states belonging to attractors as $\mathcal{T}_f^\delta$ for $\delta \in \{\text{SA}, \text{ASA}\}$.*

**Definition 4** (Minimal trap space)**.** *A Boolean subspace $\mathbf{h} \in \{0,1,*\}^n$ is called a* trap space *if every transition from any state in the subspace remains in the subspace (i.e., $\forall \mathbf{x} \in s(\mathbf{h}) : f(\mathbf{x}) \in s(\mathbf{h})$). A trap space is a* minimal trap space *(MTS) if no proper subset of $s(\mathbf{h})$ is a trap space (i.e., $\forall \mathbf{h}' \in \{0,1,*\}^n, s(\mathbf{h}') \subsetneq s(\mathbf{h}) : (\exists \mathbf{x} \in s(\mathbf{h}') : f(\mathbf{x}) \notin s(\mathbf{h}'))$). We denote the union of all states belonging to minimal trap spaces as $\mathcal{T}_f^{\text{MTS}}$.*

**Table 1.** Target states with different types and time spans

| **Time span** ($\lambda$) | **Type of target states** ($\delta$) | **Target states** ($\mathbb{T}(f, C)$) |
|---|---|---|
| Permanent (P) | $\delta \in \{\mathrm{FP}, \mathrm{SA}, \mathrm{ASA}, \mathrm{MTS}, \mathrm{VPTS}\}$ | $\mathcal{T}^{\delta}_{f/C}$ |
| Release (R) | $\delta \in \{\mathrm{FP}, \mathrm{SA}, \mathrm{ASA}, \mathrm{MTS}, \mathrm{VPTS}\}$ | States in $\mathcal{T}^{\delta}_{f}$ reachable† from some initial state in $\mathbb{B}^n$, while control $C$ is applied first for a given release time‡ |

†An update mode must be given. SA and ASA apply only to synchronous and asynchronous update, respectively. The state visited at the release time is also considered reachable
‡Release time must be a deterministic, positive number independent of the type of target states.

**Definition 5** (Value-propagated trap space)**.** *The* value propagation $\Phi$ *is a map* $\Phi : \{0, 1, *\}^n \to \{0, 1, *\}^n$ *that associates subspace* $\mathbf{h}$ *to the minimal subspace containing the* $\{f(\mathbf{x}) : \mathbf{x} \in s(\mathbf{h})\}$ *(the image of all states in the subspace* $\mathbf{h}$*). In other words,*

$$\forall i \in \{1, 2, \ldots, n\},\ \Phi_i(\mathbf{h}) = \begin{cases} b & \text{if } \forall \mathbf{x} \in s(\mathbf{h}) : f_i(\mathbf{x}) = b \\ * & \text{otherwise} \end{cases}$$

*The* value-propagated trap space *(VPTS) is* $\Phi^n(*^n)$*, which is the unique trap space obtained by applying value propagation n times from the all-free subspace* $*^n$ *[3]. We denote the set of states belonging to the* VPTS *as* $\mathcal{T}^{\mathrm{VPTS}}_f$*.*

##### 1.4 The universal target control problem

**Definition 6** (The universal target control problem)**.** *We are given* $n \in \mathbb{N}$ *components, a Boolean network* $f : \mathbb{B}^n \to \mathbb{B}^n$*, and a phenotype* $\varphi : \mathbb{B}^n \to \mathbb{B}$*. Suppose we have predefined target states* $\mathbb{T}$ *that returns a set of states under control* $C$*, denoted as* $\mathbb{T}(f, C)$*. The set of all* valid controls *to the corresponding* universal target control problem *is*

$$\mathrm{Sol}(\mathbb{T}, f, \varphi) := \{C \in \{0, 1, *\}^n : \forall \mathbf{x} \in \mathbb{T}(f, C), \varphi(\mathbf{x}) = 1\} \quad (1)$$

*where all target states satisfy the phenotype. A tool is* exact *to this setting if its output control set only contains valid controls in* $\mathrm{Sol}(\mathbb{T}, f, \varphi)$ *and contains all minimal valid controls in it for every input* $(f, \varphi)$*.*

Suppose we are given inputs $(f, \varphi)$, the time span $\lambda \in \{\mathrm{R}, \mathrm{P}\}$, and the type of target states $\delta \in \{\mathrm{FP}, \mathrm{SA}, \mathrm{ASA}, \mathrm{MTS}, \mathrm{VPTS}\}$. The concrete definition of the target states $\mathbb{T}$ in the universal target control problem is summarized in Table 1, following the definitions in Definition 2-5. Hence, we denote a problem setting by a pair ⟨**Time span, Type of target states**⟩, and also a tool exact to the corresponding universal target control problem interchangeably.

#### 2 Theoretical coverage relationships

##### 2.1 Overview

Here, we provide formal statements and proofs for the theoretical coverage relationships. The definition of coverage is given in Definition 7. *Theoretical coverage* here means that the coverage holds for all the following conditions:

- Every input BN and phenotype

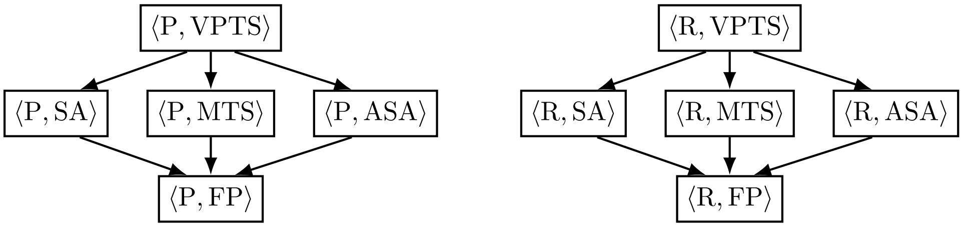


**Fig 1.** Theoretical coverage based on the type of target states and time span. Node labels are of the form ⟨Time span, Type of target states⟩. There is an arc from node A to B whenever A covers B. Transitive arcs are omitted.

- Both synchronous and asynchronous update modes
- Any positive release time, applied the same to all types of target states

Whenever there is a counterexample showing one of these conditions does not hold, we say there is no theoretical coverage relationship. The main result of this appendix is summarized in Fig 1. It shows the theoretical coverage relationships among all ten problem settings based on the type of target states and time span.

Theorem 1 states a theoretical coverage relationship for universal target control problems due to the inclusion relationship of target states. Based on this theorem, Theorem 2 provides coverage relationships based on the type of target states. Theorem 5 states that there is no theoretical coverage relationship between different time spans of control. All counterexamples are provided accordingly.

**Definition 7** (Solution coverage)**.** *Given two control sets $\mathbb{C}_1$ and $\mathbb{C}_2$, we say $\mathbb{C}_1$* covers *$\mathbb{C}_2$ if for every control in $\mathbb{C}_1$, there exists a control in $\mathbb{C}_2$ that is a subset of it:*

$$\mathbb{C}_1 \rightsquigarrow \mathbb{C}_2 \iff \forall C \in \mathbb{C}_1, \exists C' \in \mathbb{C}_2 : C \supseteq C' \ .$$

*Fix a set of inputs $\mathcal{P}$. Let $\mathbb{C}_t^{f,\varphi}$ denote the control sets returned by tool $t$ for input $(f, \varphi) \in \mathcal{P}$. We say tool $A$* covers *tool $B$ if for every input in $\mathcal{P}$, the control set of $A$ covers that of $B$:*

$$(A \rightsquigarrow B) := (\forall (f, \varphi) \in \mathcal{P} : \mathbb{C}_A^{f,\varphi} \rightsquigarrow \mathbb{C}_B^{f,\varphi}).$$

**Theorem 1** (Theoretical coverage based on target states inclusion)**.** *Let $A$ and $B$ be two tools exact to the universal target control problem in Definition 6 with target states $\mathbb{T}_A$ and $\mathbb{T}_B$, respectively. If $\mathbb{T}_A(f, C) \supseteq \mathbb{T}_B(f, C)$ for every input $(f, \varphi) \in \mathcal{P}$ and control $C \in \{0, 1, *\}^n$, then $A \rightsquigarrow B$ holds.*

*Proof.* Intuitively, the phenotype is easier to satisfy for smaller target states since the goal is to satisfy the phenotype in *all* target states. In other words, $(\mathbf{x} \in \mathbb{T}_A(f, C)) \Rightarrow \varphi(\mathbf{x})$ implies $(\mathbf{x} \in \mathbb{T}_B(f, C)) \Rightarrow \varphi(\mathbf{x})$. Therefore, $\mathrm{Sol}(\mathbb{T}_A, f, \varphi) \subseteq \mathrm{Sol}(\mathbb{T}_B, f, \varphi)$ for every input $(f, \varphi) \in \mathcal{P}$. For every control $C_A \in \mathrm{Sol}(\mathbb{T}_A, f, \varphi)$, it is also contained in $\mathrm{Sol}(\mathbb{T}_B, f, \varphi)$, implying that there exists a control $C_B \in \mathrm{Sol}(\mathbb{T}_B, f, \varphi)$ such that $C_B \subseteq C_A$ since $B$ is exact. This concludes that $A \rightsquigarrow B$ by Definition 7. □

#### 2.2 Theoretical coverage by types of target states

As illustrated in Fig 1, the same coverage relationships hold for both permanent and release control regarding the type of target states. Theorem 2 formalizes these relationships based on the inclusion relationships among the five types of target states given in Theorem 1. If there is no theoretical coverage relationship, we provide counterexamples accordingly (See Examples 3 and 4).

**Theorem 2** (Coverage based on the type of target states)**.** *For a fixed time span $\lambda \in \{\mathrm{R}, \mathrm{P}\}$, consider two types of target states $\delta_A$ and $\delta_B$. Then, the following coverage relationships hold:*

1. *If $\delta_A = \mathrm{VPTS}$ and $\delta_B \in \{\mathrm{SA}, \mathrm{ASA}, \mathrm{MTS}\}$, $\langle \lambda, \delta_A \rangle \rightsquigarrow \langle \lambda, \delta_B \rangle$ holds, but there are counterexamples showing $\langle \lambda, \delta_B \rangle \not\rightsquigarrow \langle \lambda, \delta_A \rangle$.*

2. *If $\delta_A \in \{\mathrm{SA}, \mathrm{ASA}, \mathrm{MTS}\}$ and $\delta_B = \mathrm{FP}$, $\langle \lambda, \delta_A \rangle \rightsquigarrow \langle \lambda, \delta_B \rangle$ holds, but there are counterexamples showing $\langle \lambda, \delta_B \rangle \not\rightsquigarrow \langle \lambda, \delta_A \rangle$.*

3. *If $\delta_A$ and $\delta_B$ are distinct elements in $\{\mathrm{SA}, \mathrm{ASA}, \mathrm{MTS}\}$, there are counterexamples showing $\langle \lambda, \delta_A \rangle \not\rightsquigarrow \langle \lambda, \delta_B \rangle$ and $\langle \lambda, \delta_B \rangle \not\rightsquigarrow \langle \lambda, \delta_A \rangle$.*

*By the transitivity of coverage, these results completely characterize the coverage relationships among all five types of target states.*

*Proof.* The proof for the coverage is based on Theorem 1 as below. The same result would hold if (i) $f$ is replaced by $f/C$ for permanent control and (ii) the target states are restricted to reachable ones after the same release time.

1. Since the value propagation fixes values when all transitions lead to the same value after one step, all attractors must be consistent with it. This implies that every SA, ASA, or MTS is contained in a VPTS. Therefore, $\mathcal{T}_f^{\delta} \subseteq \mathcal{T}_f^{\mathrm{VPTS}}$ for $\delta \in \{\mathrm{SA}, \mathrm{ASA}, \mathrm{MTS}\}$. However, the reverse inclusion does not hold in general due to [3] (See Example 4).

2. By definition, a FP is a special case of SA and ASA where only one state is present. In addition, each FP itself as a subspace forms a MTS. Therefore, $\mathcal{T}_f^{\mathrm{FP}} \subseteq \mathcal{T}_f^{\delta}$ for $\delta \in \{\mathrm{SA}, \mathrm{ASA}, \mathrm{MTS}\}$. The reverse inclusion does not hold in general because an attractor or a trap space may contain multiple states. A remarkable note is that a BN always has at least one SA, ASA, and MTS while it may have no FP. In such cases, $\mathcal{T}_f^{\mathrm{FP}} = \emptyset$, and every control is valid for the FP target states. This does not contradict the coverage since $\mathcal{T}_f^{\mathrm{FP}} \subseteq \mathcal{T}_f^{\delta}$ still holds, and $\emptyset$ is a subset of every set.

3. Counterexamples in [3] show that $\mathcal{T}_f^{\mathrm{SA}}$ and $\mathcal{T}_f^{\mathrm{ASA}}$ do not include $\mathcal{T}_f^{\mathrm{MTS}}$ and vice versa for general BNs. Although they did not mention it, the same BNs also work as counterexamples for the inclusion between SA and ASA (See Example 3).

□

**Example 3** (No theoretical coverage among SA, ASA, and MTS)**.** *The counterexamples are illustrated in Fig 2 with the same phenotype $\varphi = \neg \mathbf{x}_1$. As we use empty control set $\emptyset$, the claim holds for both permanent and release control.*

- $\langle \lambda, \mathrm{ASA} \rangle \not\rightsquigarrow \langle \lambda, \mathrm{SA} \rangle$ *and* $\langle \lambda, \mathrm{ASA} \rangle \not\rightsquigarrow \langle \lambda, \mathrm{MTS} \rangle$*: The BN in Fig 2a has one* SA $\{010, 101\}$, *one* ASA $\{011, 001, 000\}$, *and a minimal trap space* $\mathbb{B}^3$ *(See Fig 2d and Fig 2e). An empty control $\emptyset$ is a minimal control for* ASA, *but not for* SA *and* MTS.

- $\langle \lambda, \mathrm{SA} \rangle \not\rightsquigarrow \langle \lambda, \mathrm{ASA} \rangle$ *and* $\langle \lambda, \mathrm{SA} \rangle \not\rightsquigarrow \langle \lambda, \mathrm{MTS} \rangle$*: The BN in Fig 2b has one* SA $\{010, 001\}$, *one* ASA *spanning* $\mathbb{B}^3$, *and a minimal trap space* $\mathbb{B}^3$ *(See Fig 2f and Fig 2g). An empty control $\emptyset$ is a minimal control for* SA, *but not for* ASA *and* MTS.

$$\begin{aligned} f_1^a(\mathbf{x}) &= \mathbf{x}_2 \wedge \neg \mathbf{x}_3 \\ f_2^a(\mathbf{x}) &= \neg \mathbf{x}_2 \wedge \mathbf{x}_3 \\ f_3^a(\mathbf{x}) &= \mathbf{x}_2 \vee \neg \mathbf{x}_3 \end{aligned}$$

**(a)** BN in Figure 4a in [3]

$$\begin{aligned} f_1^b(\mathbf{x}) &= \mathbf{x}_2 \wedge \mathbf{x}_3 \\ f_2^b(\mathbf{x}) &= \mathbf{x}_3 \\ f_3^b(\mathbf{x}) &= \neg \mathbf{x}_3 \end{aligned}$$

**(b)** BN in Figure 4b in [3]

$$\begin{aligned} f_1^c(\mathbf{x}) &= (\mathbf{x}_1 \vee \mathbf{x}_2) \wedge \neg \mathbf{x}_3 \\ f_2^c(\mathbf{x}) &= \neg \mathbf{x}_1 \wedge (\mathbf{x}_2 \vee \mathbf{x}_3) \\ f_3^c(\mathbf{x}) &= \neg \mathbf{x}_2 \wedge (\mathbf{x}_1 \vee \mathbf{x}_3) \end{aligned}$$

**(c)** BN in Figure 4c in [3]

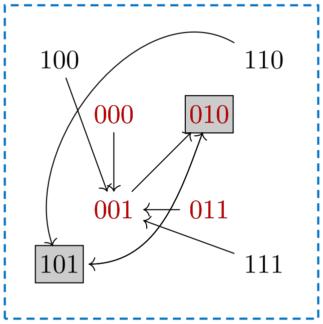


**(d)** Synchronous STG of $f^a$

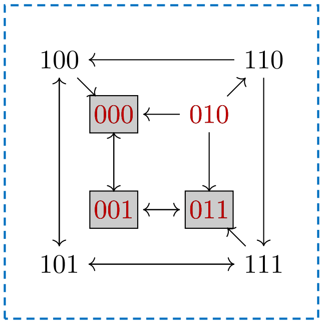


**(e)** Asynchronous STG of $f^a$

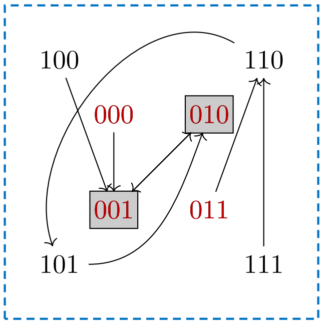


**(f)** Synchronous STG of $f^b$

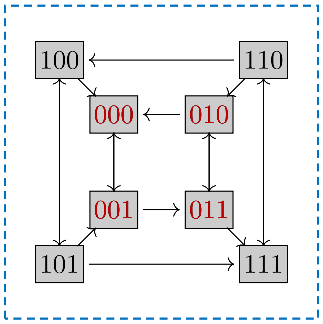


**(g)** Asynchronous STG of $f^b$

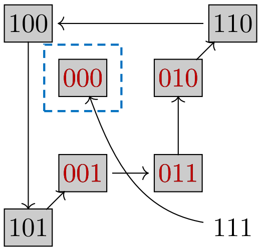


**(h)** Synchronous STG of $f^c$

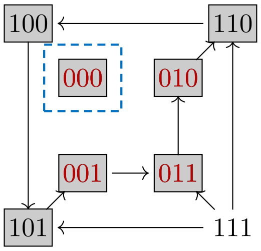


**(i)** Asynchronous STG of $f^c$

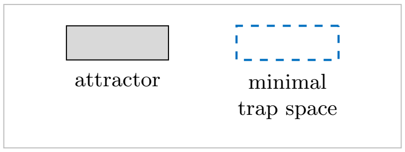


**Fig 2.** BNs adopted from Figure 4 [3] with phenotype $\varphi = \neg \mathbf{x}_1$ (colored red).

**Table 2.** Theoretical coverage between permanent and release control ($\langle \lambda_A, \delta_A \rangle \not\rightsquigarrow \langle \lambda_B, \delta_B \rangle$).

| Proof | Update mode | Release time | Tool A | | | Tool B | |
|---|---|---|---|---|---|---|---|
| | | | $\lambda_A$ | $\delta_A$ | | $\lambda_B$ | $\delta_B$ |
| ***Counterexamples*** | | | | | | | |
| Fig 3a | Synchronous | $\geq 1$ | P | $\in \{\mathrm{FP}, \mathrm{SA}, \mathrm{MTS}, \mathrm{VPTS}\}$ | $\not\rightsquigarrow$ | R | $\in \{\mathrm{FP}, \mathrm{SA}, \mathrm{MTS}, \mathrm{VPTS}\}$ |
| Fig 3a | Asynchronous | $\geq 1$ | P | $\in \{\mathrm{FP}, \mathrm{ASA}, \mathrm{MTS}, \mathrm{VPTS}\}$ | $\not\rightsquigarrow$ | R | $\in \{\mathrm{FP}, \mathrm{ASA}, \mathrm{MTS}, \mathrm{VPTS}\}$ |
| Fig 3a | Synchronous | $\geq 1$ | R | $\in \{\mathrm{FP}, \mathrm{SA}, \mathrm{MTS}, \mathrm{VPTS}\}$ | $\not\rightsquigarrow$ | P | $\in \{\mathrm{FP}, \mathrm{SA}, \mathrm{MTS}, \mathrm{VPTS}\}$ |
| Fig 3a† | Asynchronous | $\geq 2$ | R | $\in \{\mathrm{FP}, \mathrm{ASA}, \mathrm{MTS}, \mathrm{VPTS}\}$ | $\not\rightsquigarrow$ | P | $\in \{\mathrm{FP}, \mathrm{ASA}, \mathrm{MTS}, \mathrm{VPTS}\}$ |
| Fig 3e† | Asynchronous | $= 1$ | R | $\in \{\mathrm{FP}, \mathrm{ASA}, \mathrm{MTS}\}$ | $\not\rightsquigarrow$ | P | $\in \{\mathrm{FP}, \mathrm{ASA}, \mathrm{MTS}, \mathrm{VPTS}\}$ |
| Fig 3h† | Asynchronous | $= 1$ | R | $\in \{\mathrm{VPTS}\}$ | $\not\rightsquigarrow$ | P | $\in \{\mathrm{MTS}, \mathrm{VPTS}\}$ |
| ***Theoretical coverage (special case)*** | | | | | | | |
| Theorem 7† | Asynchronous | $= 1$ | R | $\in \{\mathrm{VPTS}\}$ | $\rightsquigarrow$ | P | $\in \{\mathrm{FP}, \mathrm{ASA}\}$ |

†Together characterize all possible combinations for asynchronous update mode and release time $\geq 1$.

- $\langle \lambda, \mathrm{MTS} \rangle \not\rightsquigarrow \langle \lambda, \mathrm{SA} \rangle$ *and* $\langle \lambda, \mathrm{MTS} \rangle \not\rightsquigarrow \langle \lambda, \mathrm{ASA} \rangle$*: The BN in Fig 2c has two* SA*s* $\{\{000\}, \{011, 010, 110, 100, 101, 001\}\}$*, which also constitute* ASA*s (See Fig 2h and Fig 2i). The* FP $\{000\}$ *is a unique* MTS *satisfying the phenotype. An empty control* $\emptyset$ *is a minimal control for* MTS*, but not for* SA *and* ASA*.*

**Example 4** (No theoretical coverage from SA, ASA, or MTS to VPTS)**.** *The counterexamples are illustrated in Fig 2 with the same phenotype* $\varphi = \neg \mathbf{x}_1$*. For all three BNs, the* VPTS *is* $\mathbb{B}^3$ *under an empty control* $\emptyset$*. As we use empty control set* $\emptyset$*, the claim holds for both permanent and release control.*

- $\langle \lambda, \mathrm{ASA} \rangle \not\rightsquigarrow \langle \lambda, \mathrm{VPTS} \rangle$*: For BN in Fig 2a, an empty control* $\emptyset$ *is a minimal control for* ASA*, but not for* VPTS*.*
- $\langle \lambda, \mathrm{SA} \rangle \not\rightsquigarrow \langle \lambda, \mathrm{VPTS} \rangle$*: For BN in Fig 2b, an empty control* $\emptyset$ *is a minimal control for* SA*, but not for* VPTS*.*
- $\langle \lambda, \mathrm{MTS} \rangle \not\rightsquigarrow \langle \lambda, \mathrm{VPTS} \rangle$*: For BN in Fig 2c, an empty control* $\emptyset$ *is a minimal control for* MTS*, but not for* VPTS*.*

#### 2.3 Theoretical coverage by time span

As illustrated in Fig 1, there is no theoretical coverage between permanent and release control. We now formally state this result in Theorem 5 and provide counterexamples in Example 6. The counterexamples are summarized in Table 2.

**Theorem 5** (No theoretical coverage between permanent and release control)**.** *Suppose an update mode is given as either synchronous or asynchronous. For any combination of* $\delta_A, \delta_B \in \{\mathrm{FP}, \mathrm{SA}, \mathrm{ASA}, \mathrm{MTS}, \mathrm{VPTS}\}$*, there are counterexamples with a positive release time showing that* $\langle \mathrm{P}, \delta_A \rangle \not\rightsquigarrow \langle \mathrm{R}, \delta_B \rangle$ *and* $\langle \mathrm{R}, \delta_A \rangle \not\rightsquigarrow \langle \mathrm{P}, \delta_B \rangle$*.*

*Proof.* The counterexamples are presented in Example 6. They correspond to the cases for Fig 3a in Table 2. □

**Example 6** (No theoretical coverage between permanent and release control)**.** *The counterexamples are illustrated from Figs 3a to 3d. The synchronous and asynchronous STGs are shown in Fig 3b and Fig 3c, respectively. We use two phenotypes* $\varphi_1 = \neg \mathbf{x}_1 \wedge \mathbf{x}_2$ *(red) and* $\varphi_2 = \mathbf{x}_1 \wedge \mathbf{x}_2$ *(blue). Regardless of the type, the target states*

***Boolean network***

$f_1^a(\mathbf{x}) = 1$

$f_2^a(\mathbf{x}) = \neg\mathbf{x}_1 \vee \mathbf{x}_2$

***Phenotype***

$\varphi_1(\mathbf{x}) = \neg\mathbf{x}_1 \wedge \mathbf{x}_2$

$\varphi_2(\mathbf{x}) = \mathbf{x}_1 \wedge \mathbf{x}_2$

**(a)** Inputs with $f^a$

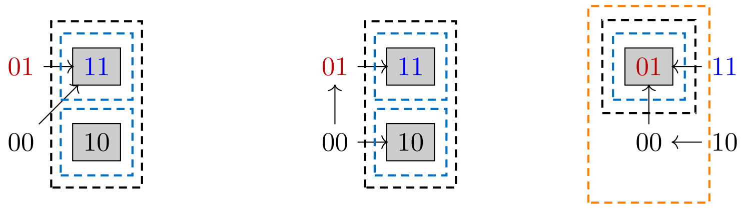


**(b)** Synch. STG of $f^a$ **(c)** Asynch. STG of $f^a$ **(d)** (A)synch. STG of $f^a/C$

***Boolean network***

$f_1^b(\mathbf{x}) = \mathbf{x}_1 \vee \mathbf{x}_2$

$f_2^b(\mathbf{x}) = \neg\mathbf{x}_1 \vee \mathbf{x}_2$

***Phenotype***

$\varphi_2(\mathbf{x}) = \mathbf{x}_1 \wedge \mathbf{x}_2$

**(e)** Inputs with $f^b$

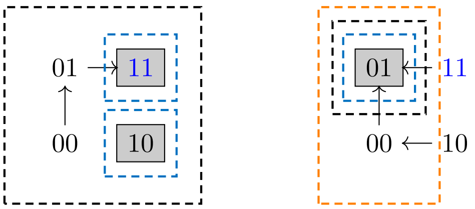


**(f)** (A)synch. STG of $f^b$ **(g)** (A)synch. STG of $f^b/C$

***Boolean network***

$f_1^c(\mathbf{x}) = \neg\mathbf{x}_3$

$f_2^c(\mathbf{x}) = (\neg\mathbf{x}_1 \wedge \mathbf{x}_2) \vee (\neg\mathbf{x}_1 \wedge \mathbf{x}_3) \vee (\mathbf{x}_2 \wedge \mathbf{x}_3)$

$f_3^c(\mathbf{x}) = (\mathbf{x}_1 \wedge \neg\mathbf{x}_2) \vee (\mathbf{x}_1 \wedge \mathbf{x}_3) \vee (\neg\mathbf{x}_2 \wedge \mathbf{x}_3)$

***Phenotype***

$\varphi_3(\mathbf{x}) = (\neg\mathbf{x}_1 \vee \neg\mathbf{x}_2 \vee \neg\mathbf{x}_3) \wedge (\mathbf{x}_1 \vee \mathbf{x}_2 \vee \mathbf{x}_3)$

**(h)** Inputs with $f^c$

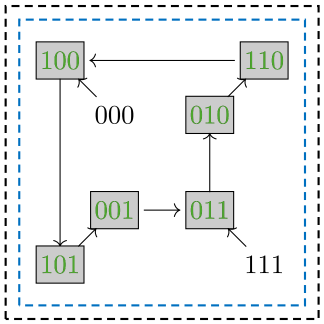


**(i)** (A)synchronous STG of $f^c$

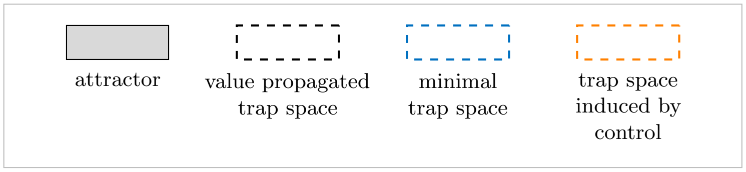


**Fig 3.** A counterexample for coverage between permanent and release control $C = \{(\mathbf{x}_1 = 0)\}$.

*under an empty control* $\emptyset$ *are* 11 *and* 10*. Since* 10 *does not satisfy either phenotype, empty control is not valid for any time span, phenotype, and type of target states. Hence, any valid control fixing one component is minimal (See Fig 3).*

*We consider control* $C = \{(\mathbf{x}_1 = 0)\}$ *both as permanent and release control. Applying control* $C$ *permanently,* 01 *is the unique* FP, SA, ASA, MTS, *and* VPTS *(See Fig 3d). It satisfies the phenotype* $\varphi_1$*, but not* $\varphi_2$*. Hence,* $C$ *is a minimal valid permanent control for* $\varphi_1$ *but not for* $\varphi_2$*.*

*Now consider control* $C$ *as a release control. For synchronous update,* 11 *is the unique reachable* FP, SA, MTS, *and* VPTS *after release. Start from* 10 *under control (See Fig 3d),* 01 *is reached after one transition. After release,* 10 *is no longer reachable in the original synchronous STG (See Fig 3b). Therefore,* 11 *is the unique reachable* FP, SA, MTS, *and* VPTS *under synchronous update. It satisfies the phenotype* $\varphi_2$*, but not* $\varphi_1$*.*

*For the asynchronous update, however,* 10 *is still reachable after releasing the control at* 00 *in the original asynchronous STG (See Fig 3c). Therefore, there is a target state* 10 *that does not satisfy either phenotype if release time is one. If we consider a release time of at least two,* 10 *is no longer reachable because the control is released at* 01 *(See Fig 3d). Thus, for asynchronous update with release time at least two,* 11 *is the unique reachable* FP, ASA, MTS, *and* VPTS *after release. It satisfies the phenotype* $\varphi_2$*, but not* $\varphi_1$*, just like the synchronous case.*

*Using the above observations, we can show the following counterexamples:*

- $\langle \mathrm{P}, \delta_A \rangle \not\rightsquigarrow \langle \mathrm{R}, \delta_B \rangle$ *for all* $\delta_A, \delta_B \in \{\mathrm{FP}, \mathrm{SA}, \mathrm{ASA}, \mathrm{MTS}, \mathrm{VPTS}\}$*: Control* $C$ *is a valid permanent control for phenotype* $\varphi_1$ *with all five types of target states and it is minimal. However, it is not a valid release control for both update modes, regardless of the release time.*
- $\langle \mathrm{R}, \delta_A \rangle \not\rightsquigarrow \langle \mathrm{P}, \delta_B \rangle$ *for all* $\delta_A, \delta_B \in \{\mathrm{FP}, \mathrm{SA}, \mathrm{ASA}, \mathrm{MTS}, \mathrm{VPTS}\}$*: Consider one of the following two cases:*
  - *For synchronous update with any positive release time*
  - *For asynchronous update with release time at least two*

  *Then, control* $C$ *is a valid release control for phenotype* $\varphi_2$ *with all five types of target states and it is minimal. However, it is not a valid permanent control.*

##### 2.4 Special case: Asynchronous update with release time of one

The previous counterexample suffices to disprove theoretical coverage, but a question still remains for asynchronous update with release time equal to one: Will there be theoretical coverage for this special case?

Surprisingly, $\langle \mathrm{R}, \mathrm{VPTS} \rangle \rightsquigarrow \langle \mathrm{P}, \mathrm{ASA} \rangle$ holds as we show in Theorem 7. The intuition is that a minimal valid release control to VPTS should not fix any previously fixed component in the VPTS to the opposite value. If not, one step transition can lead back to a state in VPTS that does not satisfy the phenotype. As a result, all ASA under permanent control must also belong to the VPTS under no control. By transitivity, this result extends to $\langle \mathrm{R}, \mathrm{VPTS} \rangle \rightsquigarrow \langle \mathrm{P}, \mathrm{FP} \rangle$ as well.

For the other combinations of the types of target states, we provide counterexamples in Example 8 and Example 9. Example 8 illustrates that a release control can make attractors that do not satisfy the phenotype no longer reachable. Example 9 illustrates that a release control can make transient states that do not satisfy the phenotype no longer reachable. Permanent control cannot achieve the same effect, and thus is not valid for the same phenotype.

**Theorem 7** (A special case where $\langle \mathrm{R}, \mathrm{VPTS} \rangle \rightsquigarrow \langle \mathrm{P}, \mathrm{ASA} \rangle$ holds)**.** *Consider the asynchronous update mode and release time equal to one. Given any Boolean network $f : \mathbb{B}^n \to \mathbb{B}^n$ and phenotype $\varphi : \mathbb{B}^n \to \mathbb{B}$, suppose $C \in \{0, 1, *\}^n$ is a valid minimal release control to $\langle \mathrm{R}, \mathrm{VPTS} \rangle$. Then, $C$ is also a valid permanent control for $\langle \mathrm{P}, \mathrm{ASA} \rangle$. Therefore, $\langle \mathrm{R}, \mathrm{VPTS} \rangle \rightsquigarrow \langle \mathrm{P}, \mathrm{ASA} \rangle$ holds.*

*Proof.* If $C = \emptyset$, the statement holds trivially. Therefore, consider $C \neq \emptyset$. Let $\mathbf{h}_f^{\mathrm{VPTS}}$ be the unique VPTS under no control, given as a subspace. We define $\mathcal{I}^{\mathrm{fixed}} \subseteq \{1, \ldots, n\}$ as the set of components fixed in $\mathbf{h}_f^{\mathrm{VPTS}}$. Since $C$ is not an empty control, there must be at least one state $\hat{\mathbf{x}} \in s(\mathbf{h}_f^{\mathrm{VPTS}})$ such that $\varphi(\hat{\mathbf{x}})$ does not hold.

We first show that a release control $C$ must fix all components $i \in \mathcal{I}^{\mathrm{fixed}}$ to the same value as $\mathbf{h}_{f,i}^{\mathrm{VPTS}}$. To make contradiction, suppose there is a component $i$ where $C_i \neq \mathbf{h}_{f,i}^{\mathrm{VPTS}}$. Then, $C_i$ is either $1 - \mathbf{h}_{f,i}^{\mathrm{VPTS}}$ or $*$, and we divide the case into these two subcases.

**Case (a): If** $(C_i = 1 - \mathbf{h}_{f,i}^{\mathrm{VPTS}})$**.** Applying control $C$, another state

$$\mathbf{x}' := \begin{cases} \hat{\mathbf{x}}_j & \text{if } j \neq i, \\ (1 - \mathbf{h}_{f,i}^{\mathrm{VPTS}}) & \text{if } j = i, \end{cases}$$

can be reached from $\hat{\mathbf{x}}$ after one transition under the asynchronous update. Since value propagation fixes component $i$ to $\mathbf{h}_{f,i}^{\mathrm{VPTS}}$, there must be a step in the value propagation that updates component $i$ from other fixed components. As those fixed components are also fixed the same in $\hat{\mathbf{x}}$, an arc from $\mathbf{x}'$ to $\hat{\mathbf{x}}$ must be present in the original asynchronous STG. Therefore, releasing the control at $\mathbf{x}'$, $\hat{\mathbf{x}}$ is still reachable in one transition in the original STG. As this contradicts the validity assumption of $C$ for $\langle \mathrm{R}, \mathrm{VPTS} \rangle$, no component in $\mathcal{I}^{\mathrm{fixed}}$ can be fixed differently in the control. This further implies $s(C) \subseteq s(\mathbf{h}_f^{\mathrm{VPTS}})$.

**Case (b): If** $(C_i = *)$**.** Starting from $\hat{\mathbf{x}}$, $\hat{\mathbf{x}}$ is reachable after one transition under control $C$ because $C_i$ does not fix component $i$, but the value propagation fixed it. After releasing the control at $\hat{\mathbf{x}}$, $\hat{\mathbf{x}}$ is still reachable in one transition in the original STG. Similar to subcase **Case (a)**, this contradicts the validity assumption of $C$, implying $s(C) \subseteq s(\mathbf{h}_f^{\mathrm{VPTS}})$.

From the previous argument, any state belonging to ASA under permanent control $C$ must also belong to VPTS under no control. Let $\tilde{\mathbf{x}} \in \mathcal{T}_{f/C}^{\mathrm{ASA}}$ be such a state in the ASA. As it belongs to an attractor, it should be reachable by one transition under control $C$. Therefore, $\tilde{\mathbf{x}}$ is still reachable after releasing the control at $\tilde{\mathbf{x}}$, and it also belongs to VPTS under no control. From the validity of $C$ for $\langle \mathrm{R}, \mathrm{VPTS} \rangle$, $\varphi(\tilde{\mathbf{x}})$ holds. Since $\tilde{\mathbf{x}}$ is an arbitrary state in ASA, $C$ is also a valid control for $\langle \mathrm{P}, \mathrm{ASA} \rangle$. This completes the proof. □

**Example 8** (Counterexamples: Attractors removed by release control)**.** *The counterexamples for $\langle \mathrm{R}, \delta_A \rangle \not\rightsquigarrow \langle \mathrm{P}, \delta_B \rangle$ for $\delta_A \in \{\mathrm{FP}, \mathrm{ASA}, \mathrm{MTS}\}$ and $\delta_B \in \{\mathrm{FP}, \mathrm{ASA}, \mathrm{MTS}, \mathrm{VPTS}\}$ are illustrated in Figs 3e to 3g. Consider the Boolean network $f^b$ with phenotype $\varphi_2 = \mathbf{x}_1 \wedge \mathbf{x}_2$. For this example, STGs are the same for both synchronous and asynchronous update modes. Under an empty control $\emptyset$, the (a)synchronous STG has two* FP*s:* 11 *and* 10 *(See Fig 3f). They are also the only* ASA*s and* MTS*s, and thus the target states for* FP, ASA, *and* MTS *include both* 11 *and* 10*. Since* 10 *does not satisfy $\varphi_2$, empty control is not valid for any type of target states; any valid control fixing one component is minimal.*

*Let $C = \{(\mathbf{x}_1 = 0)\}$ (as in Fig 3g) and interpret it as a* release *control with release time equal to one. Under $C$, every state reaches a state with $\mathbf{x}_1 = 0$ after one (a)synchronous transition (See Fig 3g). After releasing $C$, state* 10 *is no longer reachable in the original STG (See Fig 3f), so* 11 *is the unique reachable* FP, ASA*, and*

MTS. *Since* 11 *satisfies* $\varphi_2$, $C$ *is a minimal valid release control for* $\langle \mathrm{R}, \delta_A \rangle, \delta_A \in \{\mathrm{FP}, \mathrm{ASA}, \mathrm{MTS}\}$.

*However, applying the same* $C$ *permanently yields* 01 *as the unique* FP, ASA, MTS, *and* VPTS *(See Fig 3g). As* 01 *does not satisfy* $\varphi_2$, $C$ *is not a valid permanent control for* $\langle \mathrm{P}, \delta_B \rangle, \delta_B \in \{\mathrm{FP}, \mathrm{ASA}, \mathrm{MTS}, \mathrm{VPTS}\}$. *This provides a counterexample to the claimed coverage.*

**Example 9** (Counterexamples: Transient states removed by release control)**.** *The counterexample for* $\langle \mathrm{R}, \mathrm{VPTS} \rangle \not\rightarrow \langle \mathrm{P}, \delta_B \rangle, \delta_B \in \{\mathrm{MTS}, \mathrm{VPTS}\}$ *is illustrated in Figs 3h and 3i. Consider the Boolean network* $f^c$ *with phenotype* $\varphi_3 = (\neg \mathbf{x}_1 \vee \neg \mathbf{x}_2 \vee \neg \mathbf{x}_3) \wedge (\mathbf{x}_1 \vee \mathbf{x}_2 \vee \mathbf{x}_3)$. *For this example, STGs are the same for both synchronous and asynchronous update modes. Both the* VPTS *and the* MTS *coincide with the entire state space* $\mathbb{B}^3$ *(See Fig 3i). There are two states* 000 *and* 111 *that violate* $\varphi_3$ *in them. Hence, an empty control* $\emptyset$ *is not a valid permanent control for* MTS *and* VPTS.

*Consider an empty control* $\emptyset$ *as a release control with release time equal to one. As shown in the STG, the states* 000 *and* 111 *do not have incoming transitions and are not reachable after one transition (See Fig 3i). Hence, all reachable states satisfy* $\varphi_3$ *after release, and* $\emptyset$ *is a valid (and minimal) release control for both* MTS *and* VPTS. *This provides a counterexample to the claimed coverage.*
