## Supplementary figures and images for "Why Boolean network control tools disagree: a taxonomy of control problems"

### _graph.png

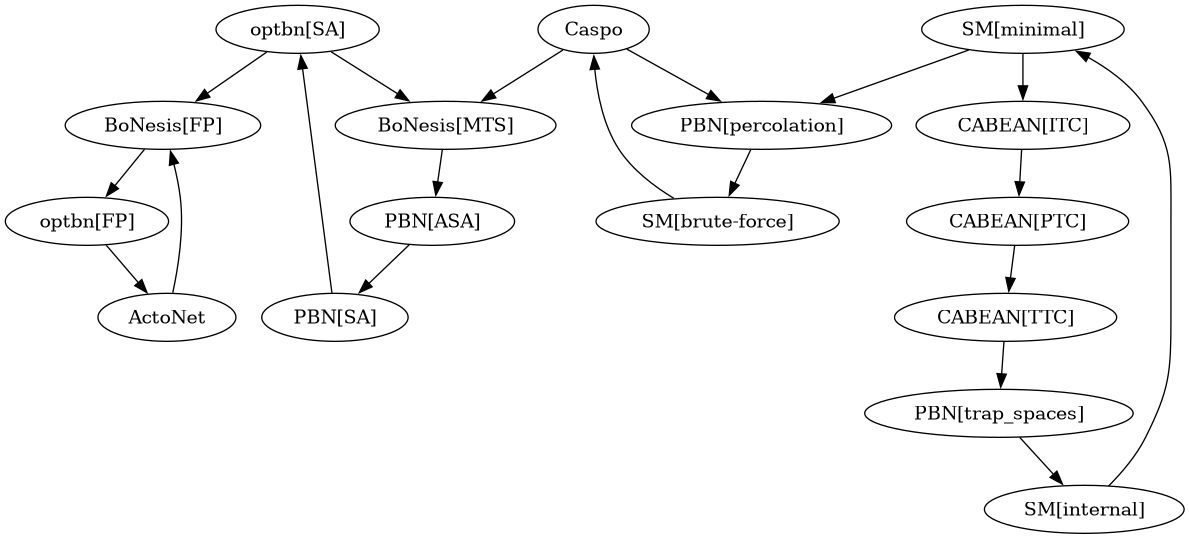

### _graph.png

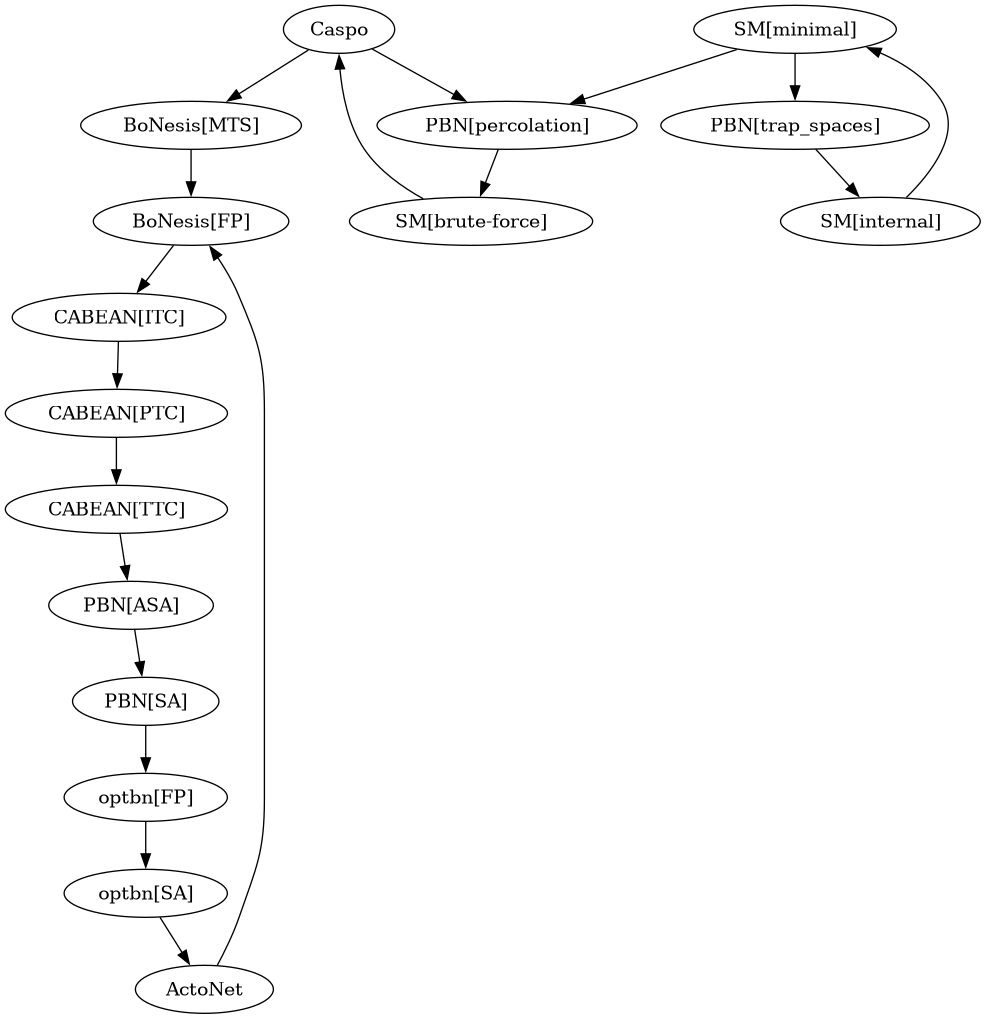

### _graph.png

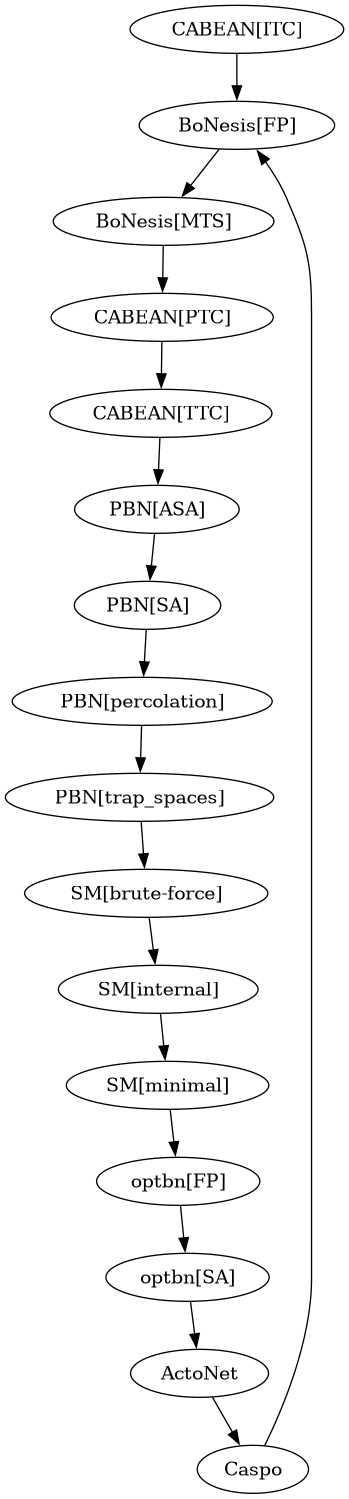

### _graph_tred.png

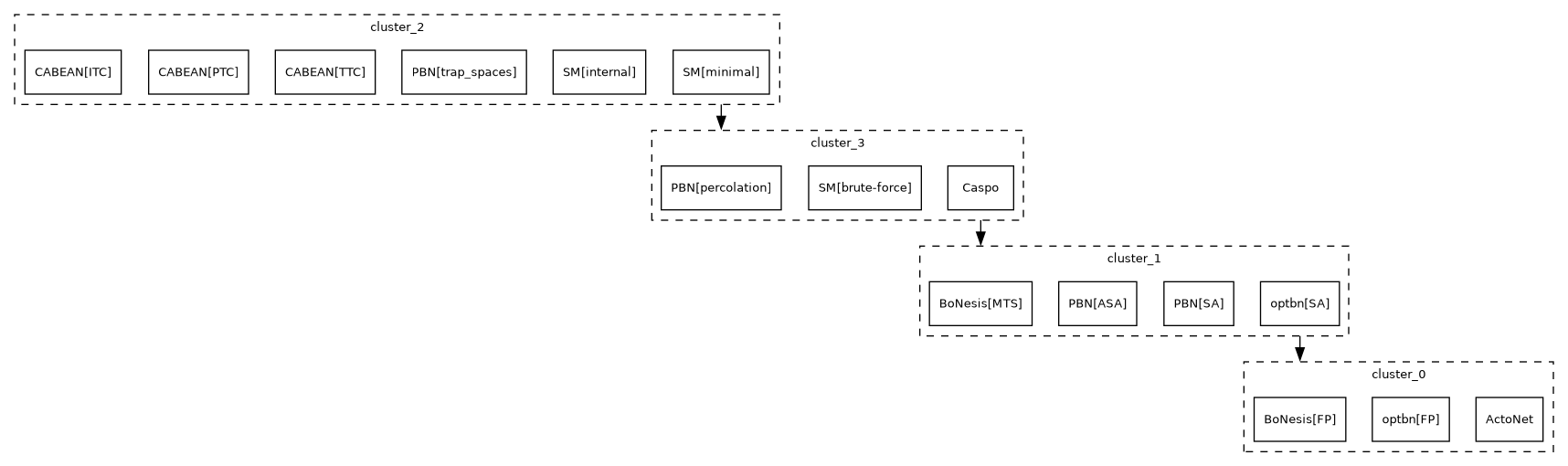

### _graph_tred.png

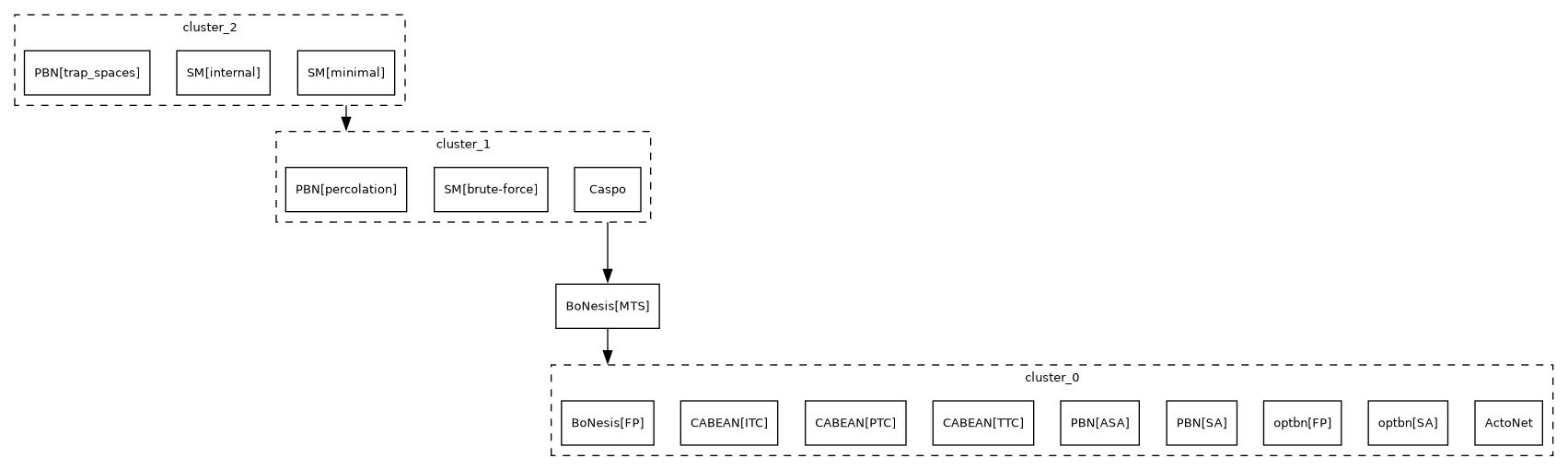

### _graph_tred.png

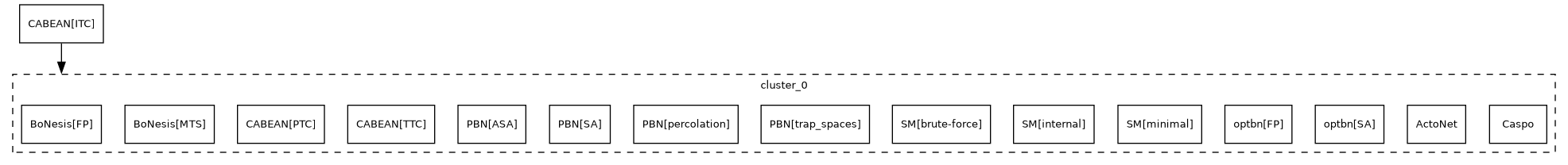

### _score_full.pdf

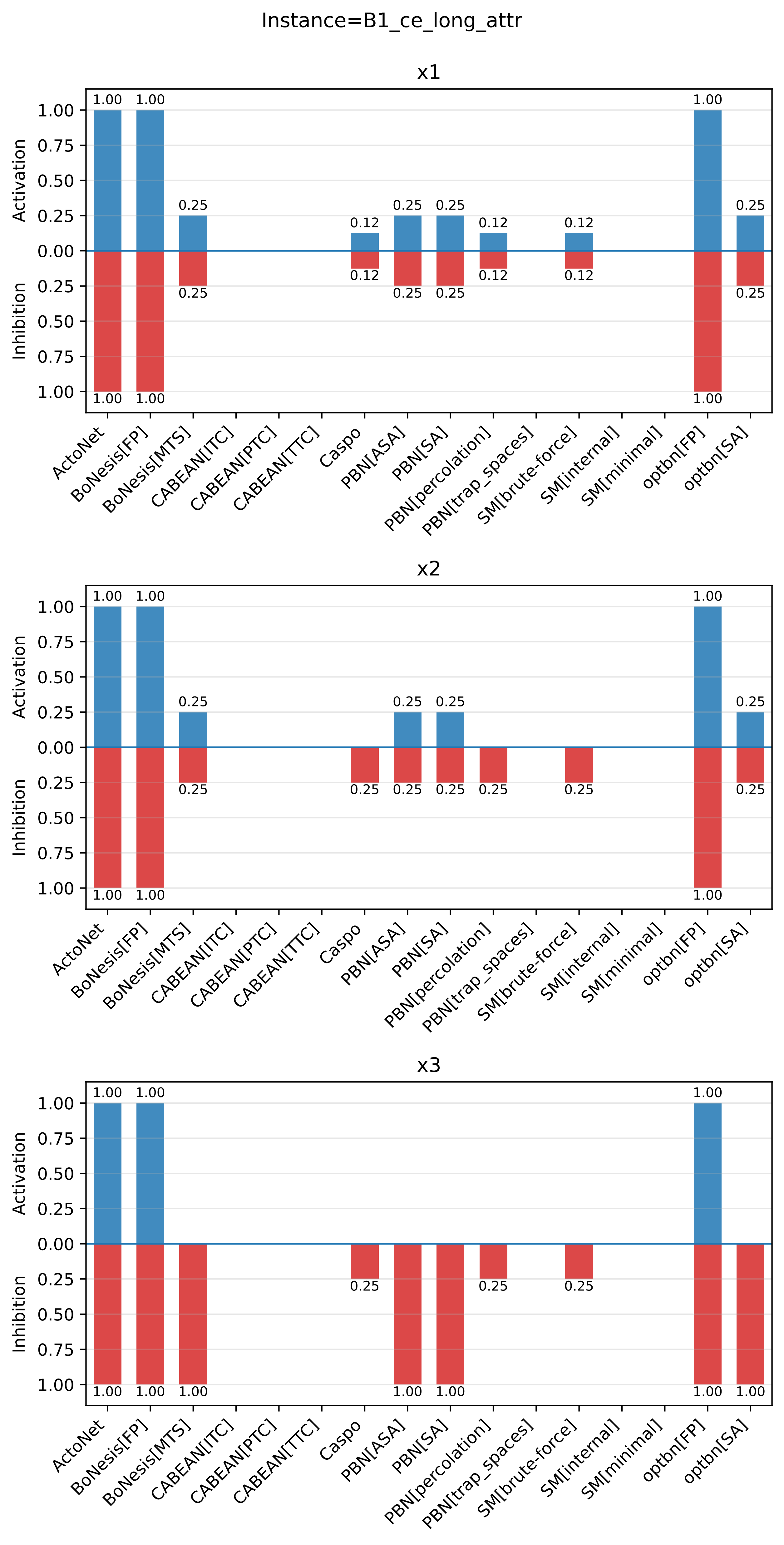

### _score_full.pdf

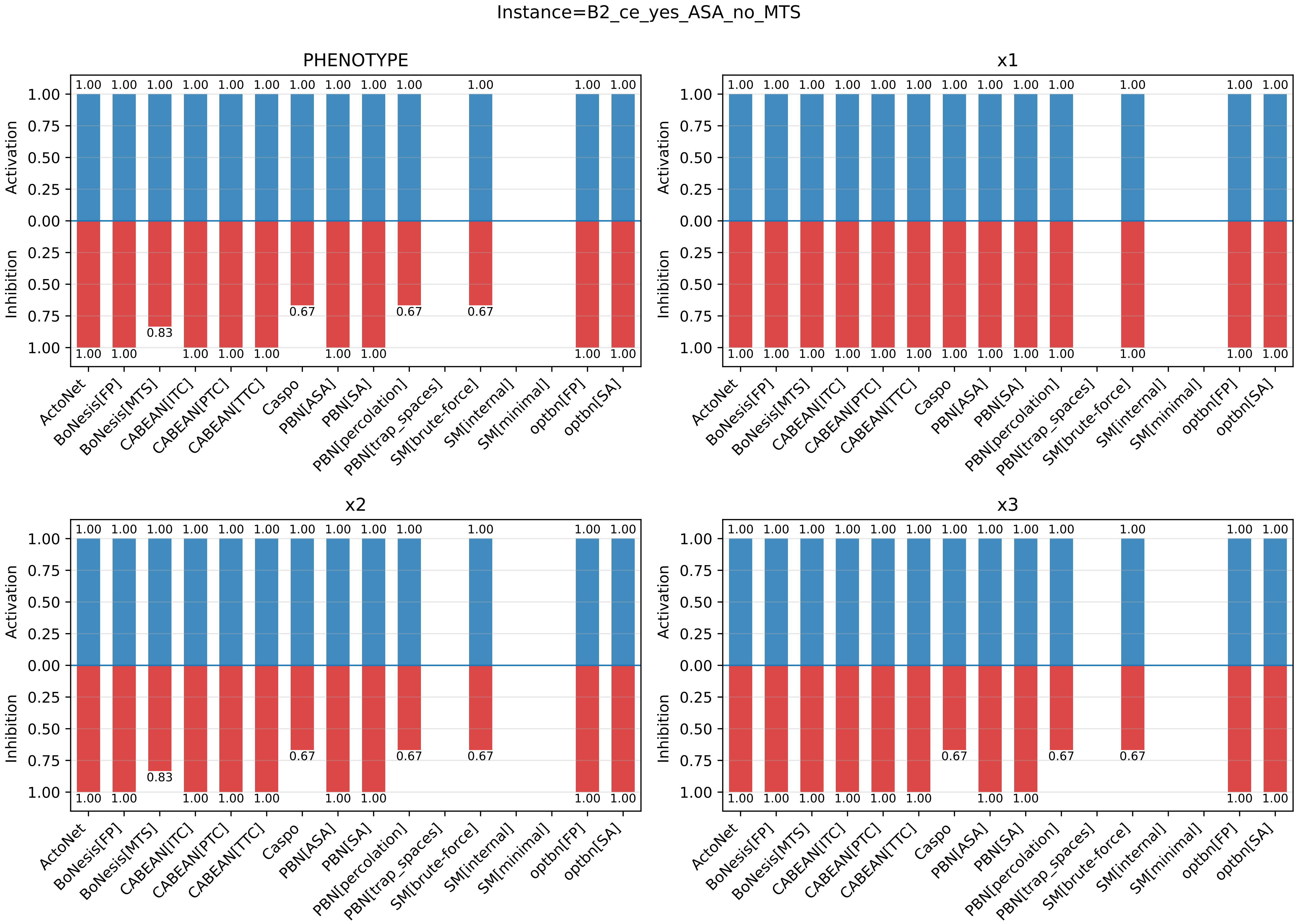

### _score_full.pdf

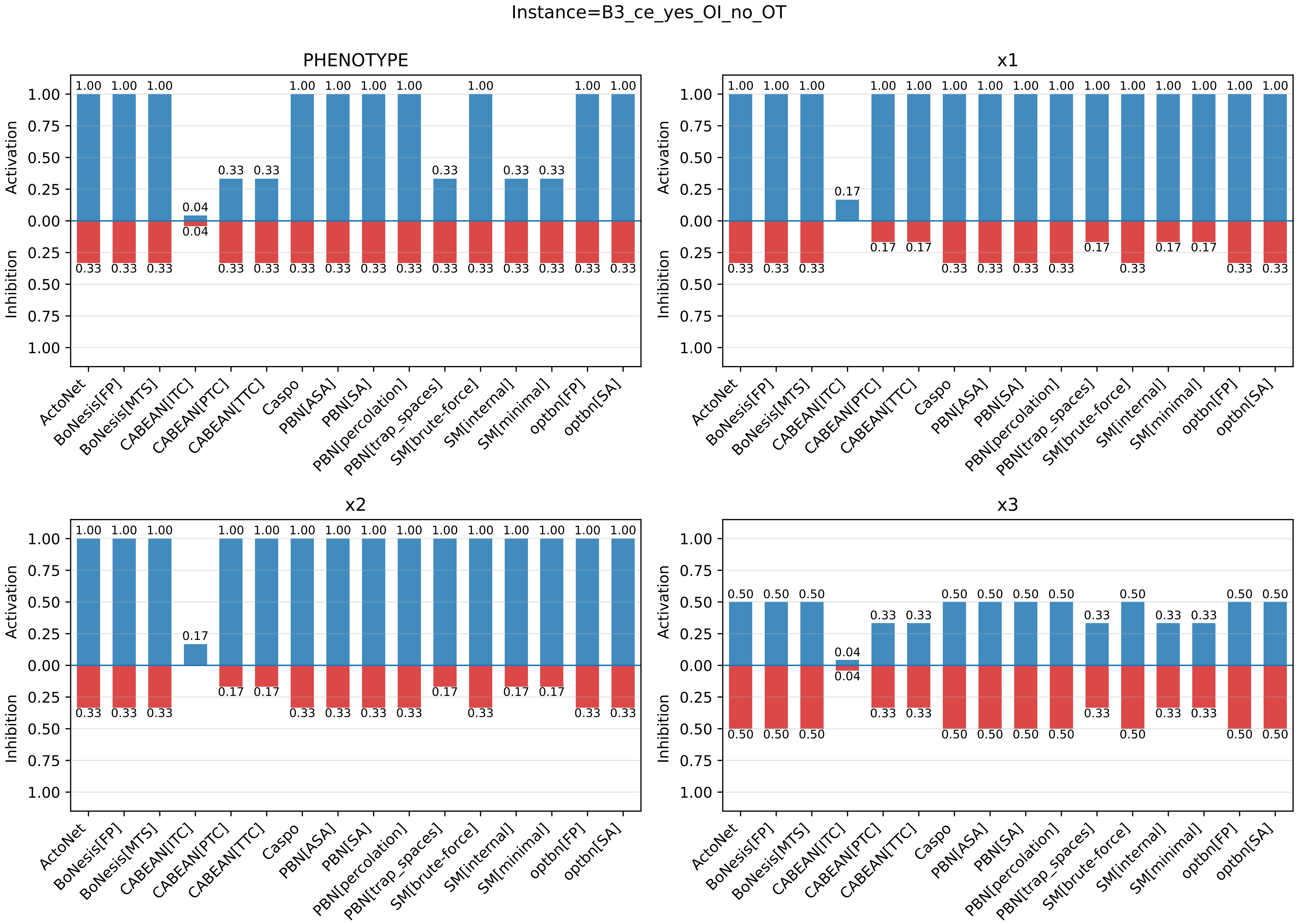

### _score_full.pdf

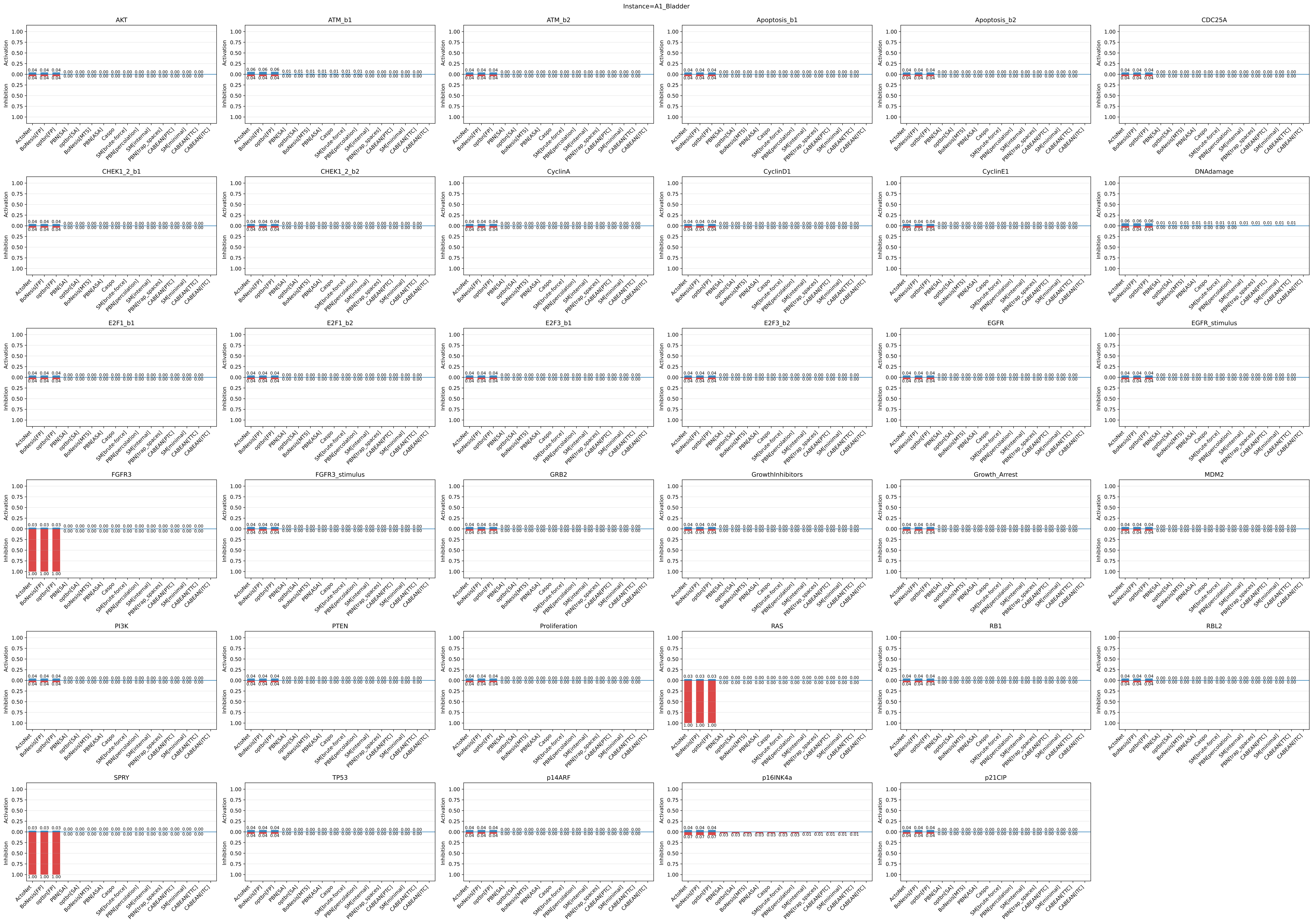

### _score_full.pdf

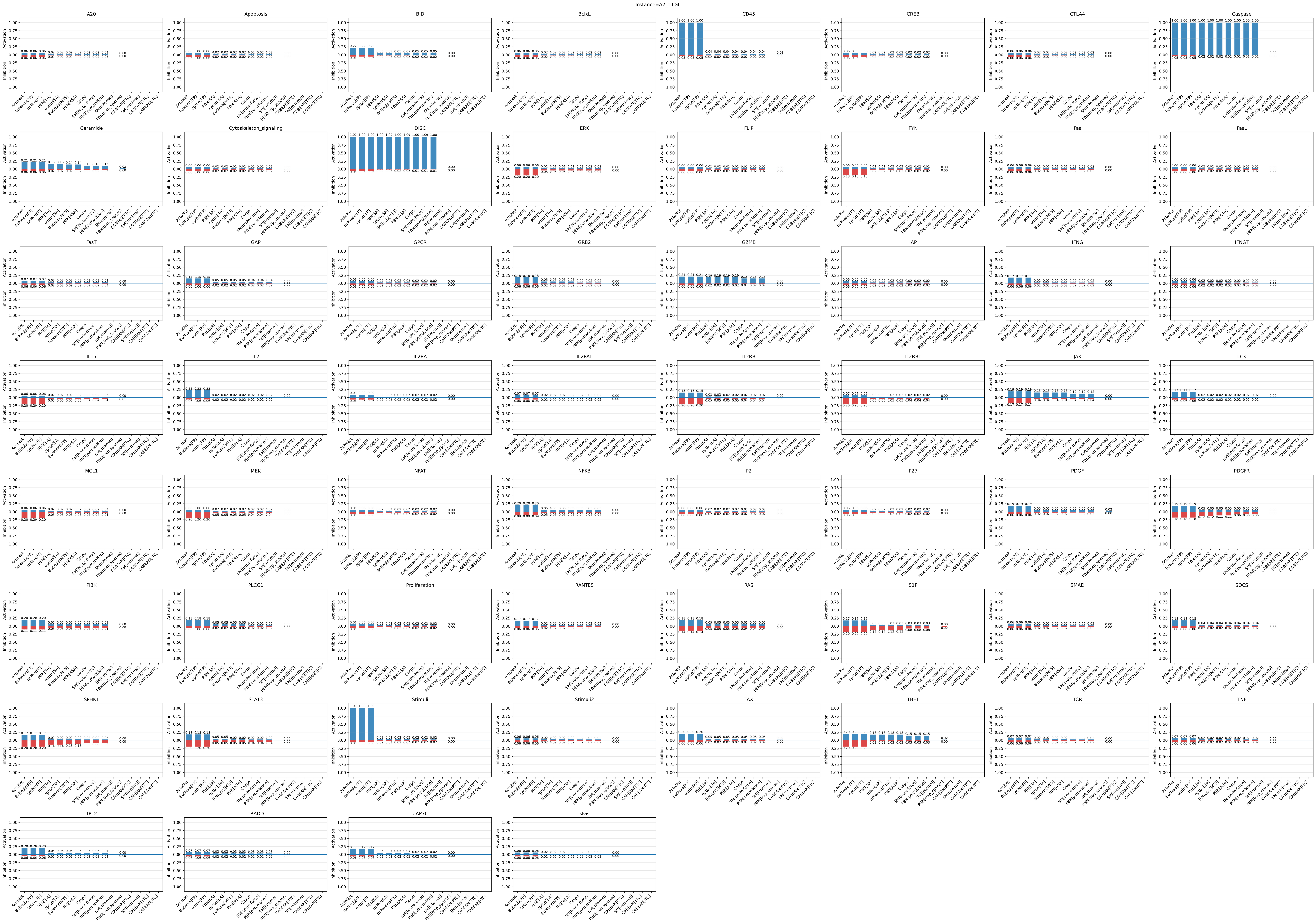

### _score_full.pdf

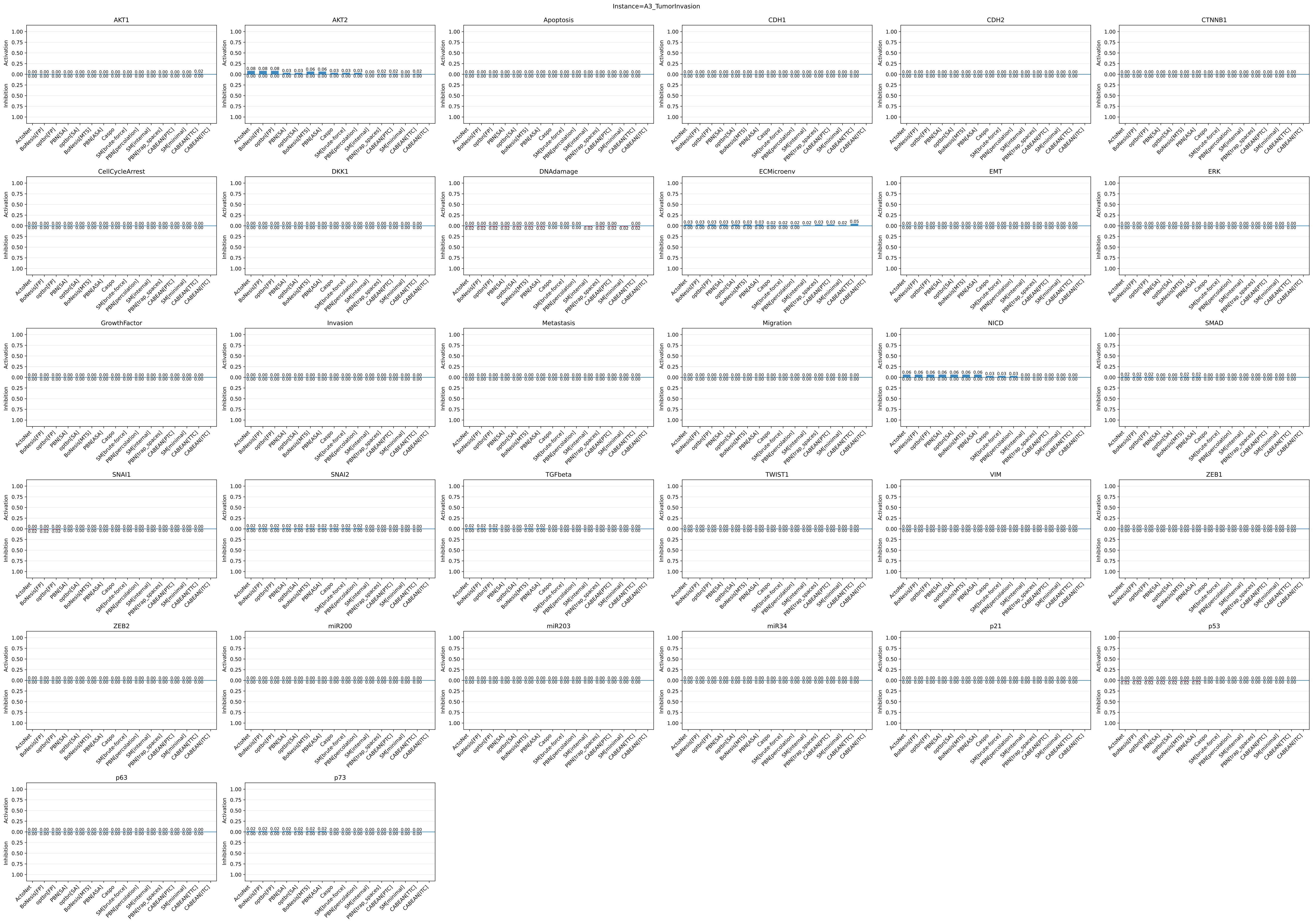

### _score_full.pdf

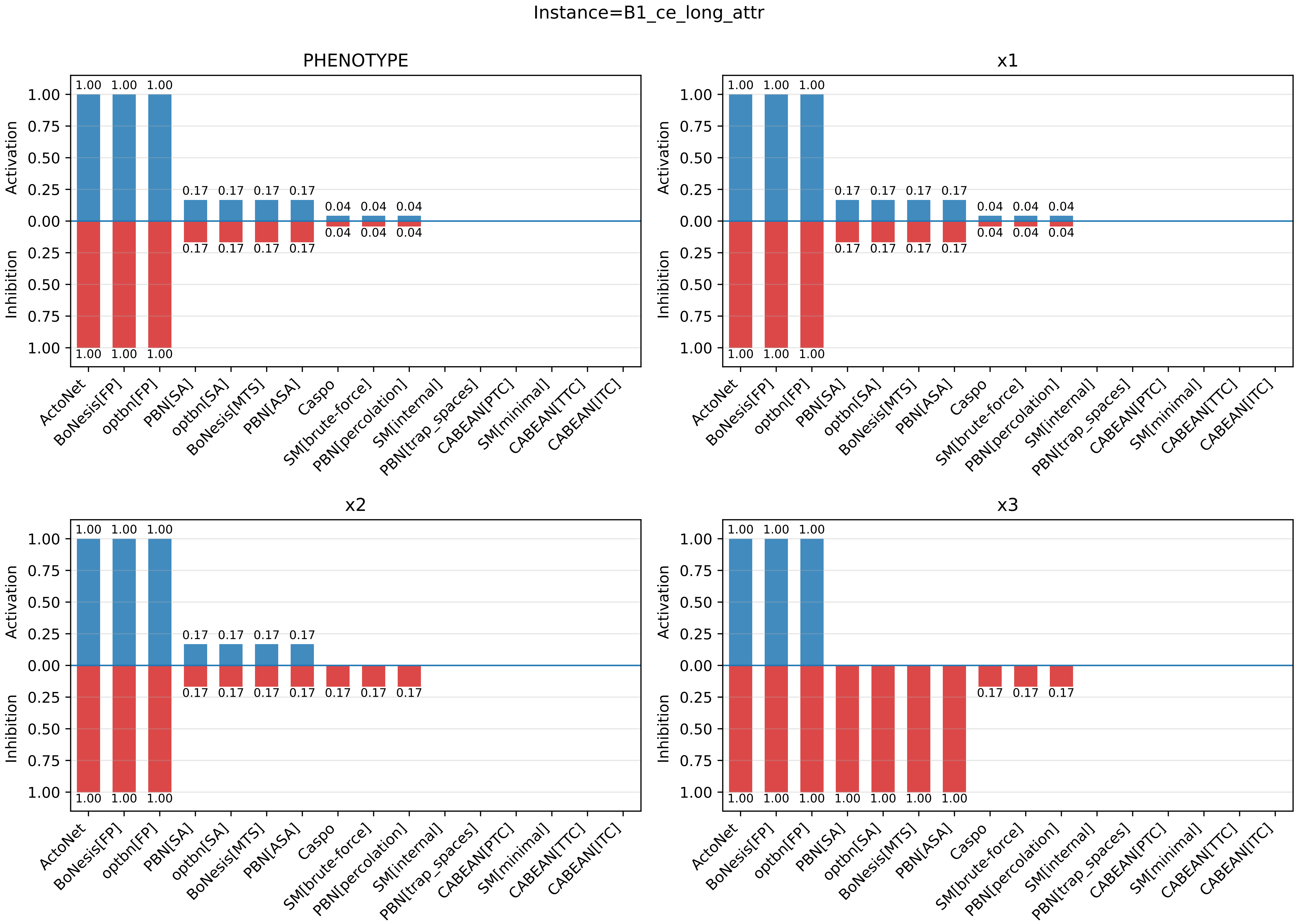

### _score_full.pdf

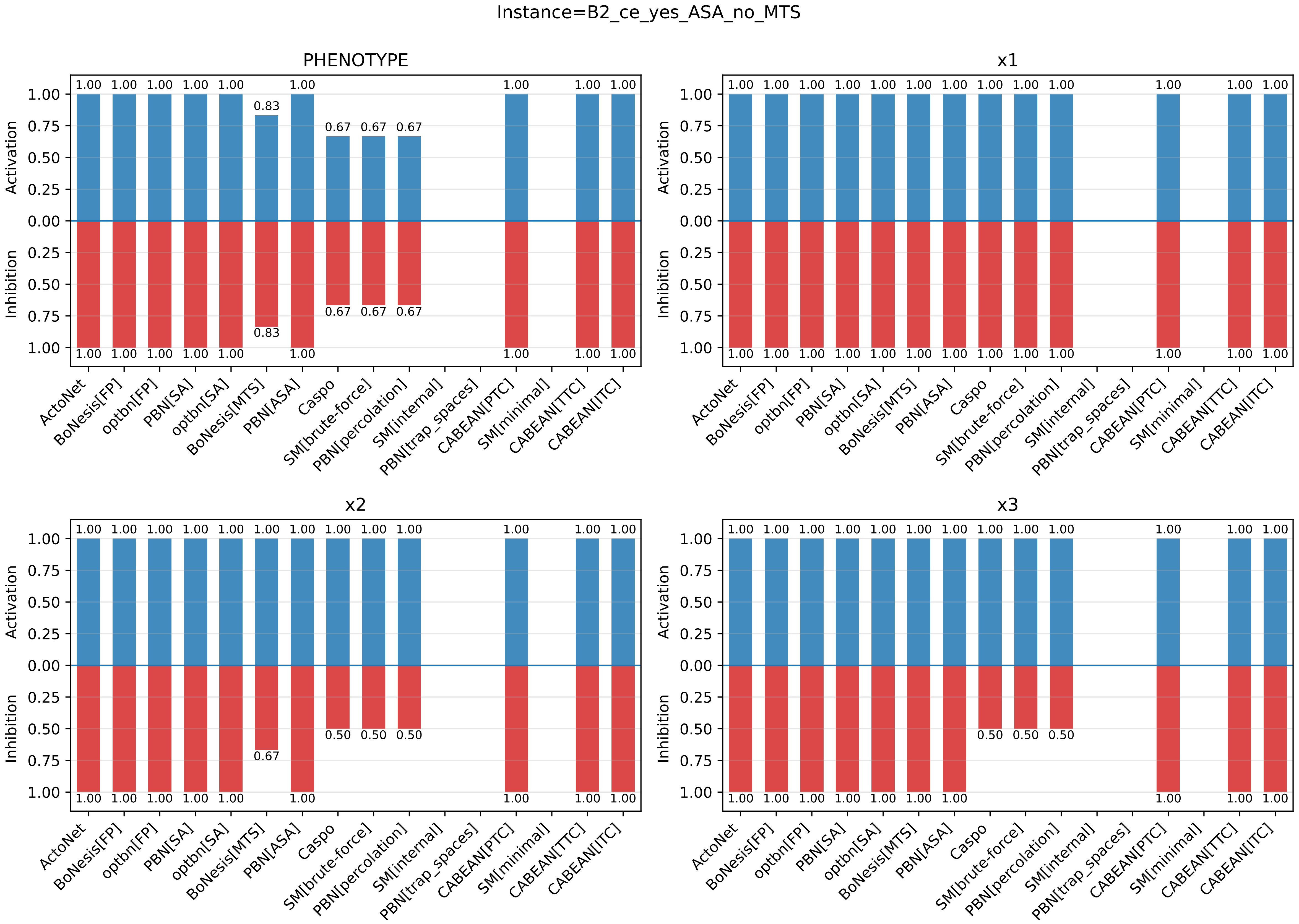

### _score_full.pdf

Instance=B3_ce_yes_OI_no_OT

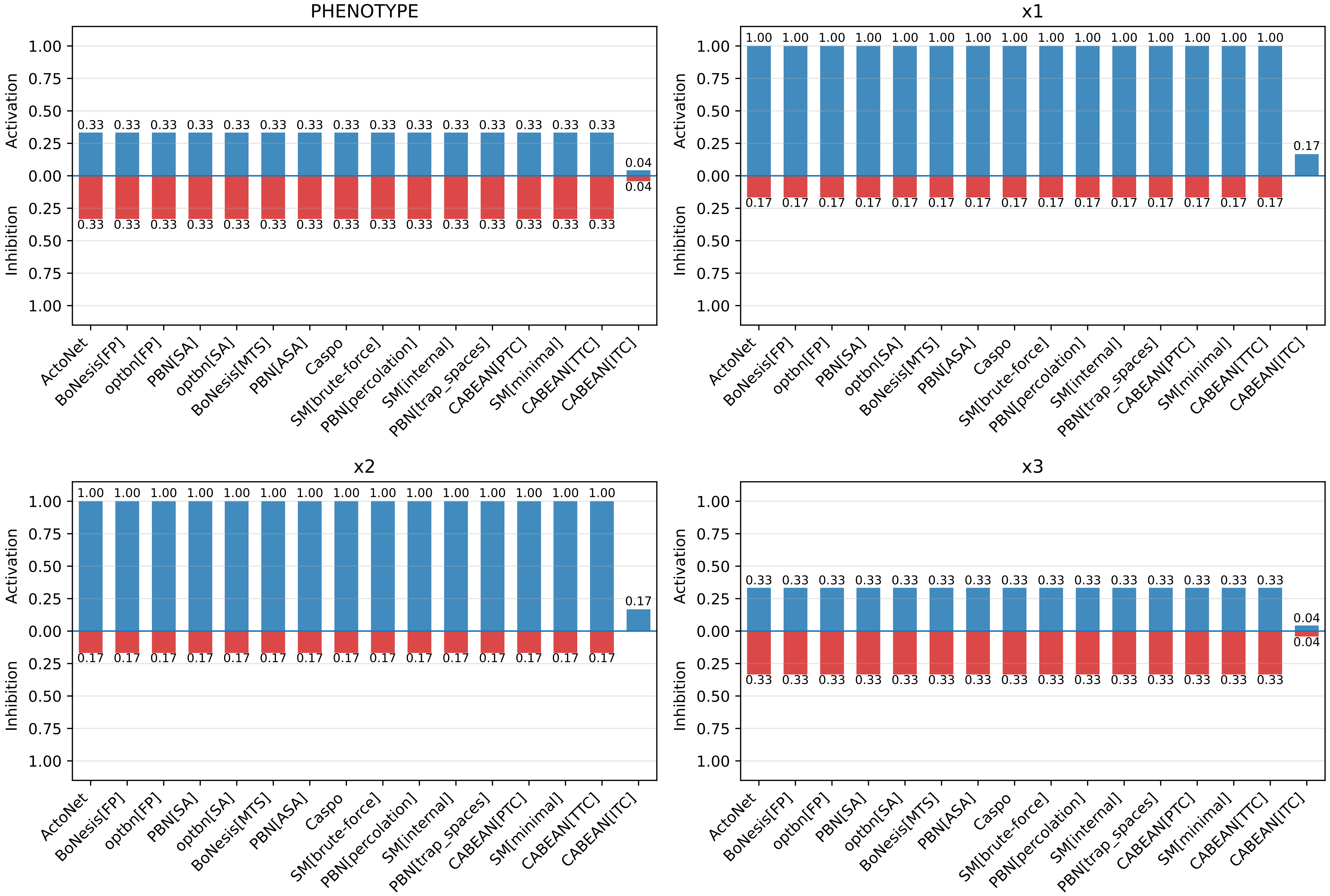

### _score_full.pdf

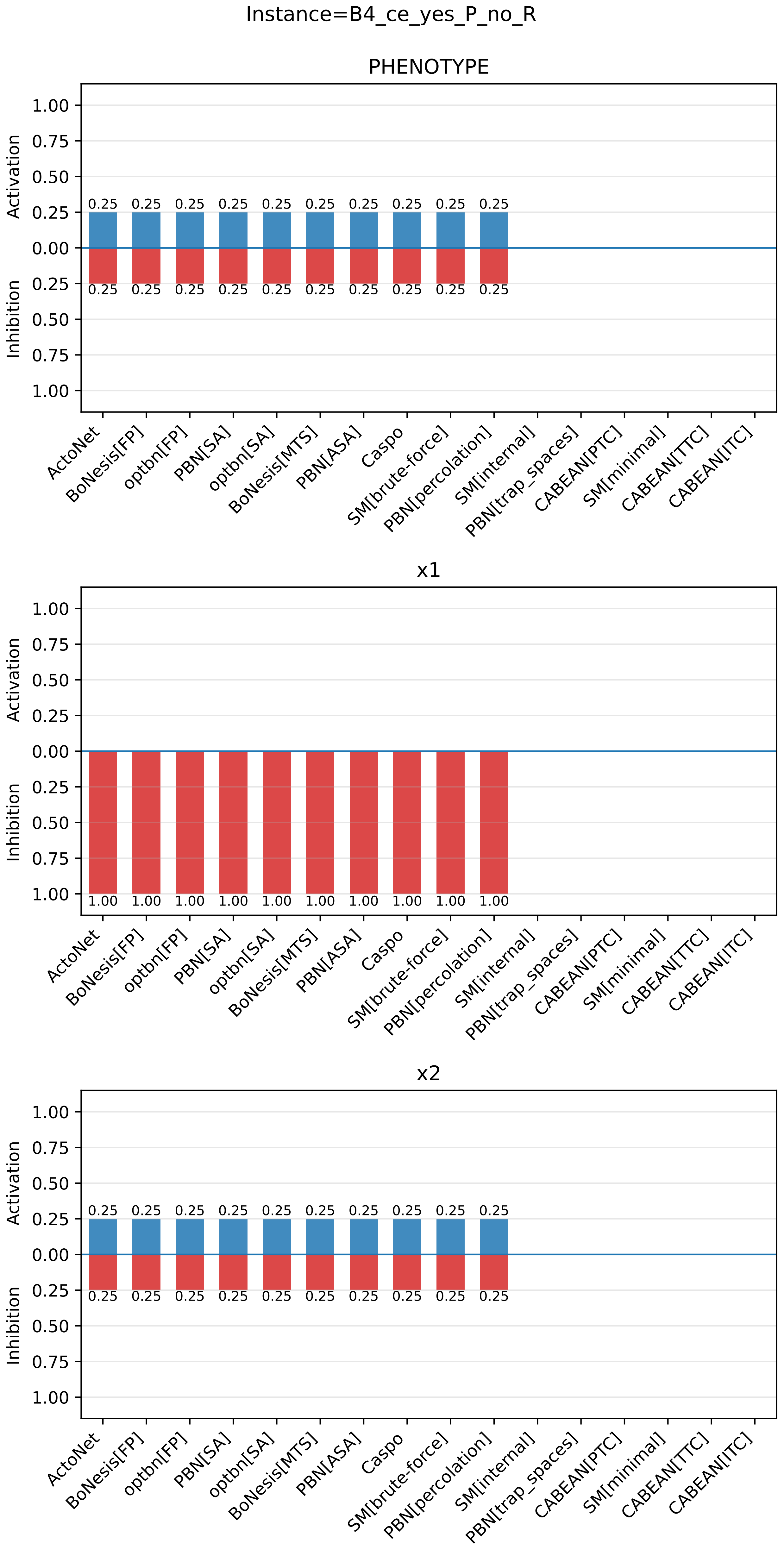

### _score_full.pdf

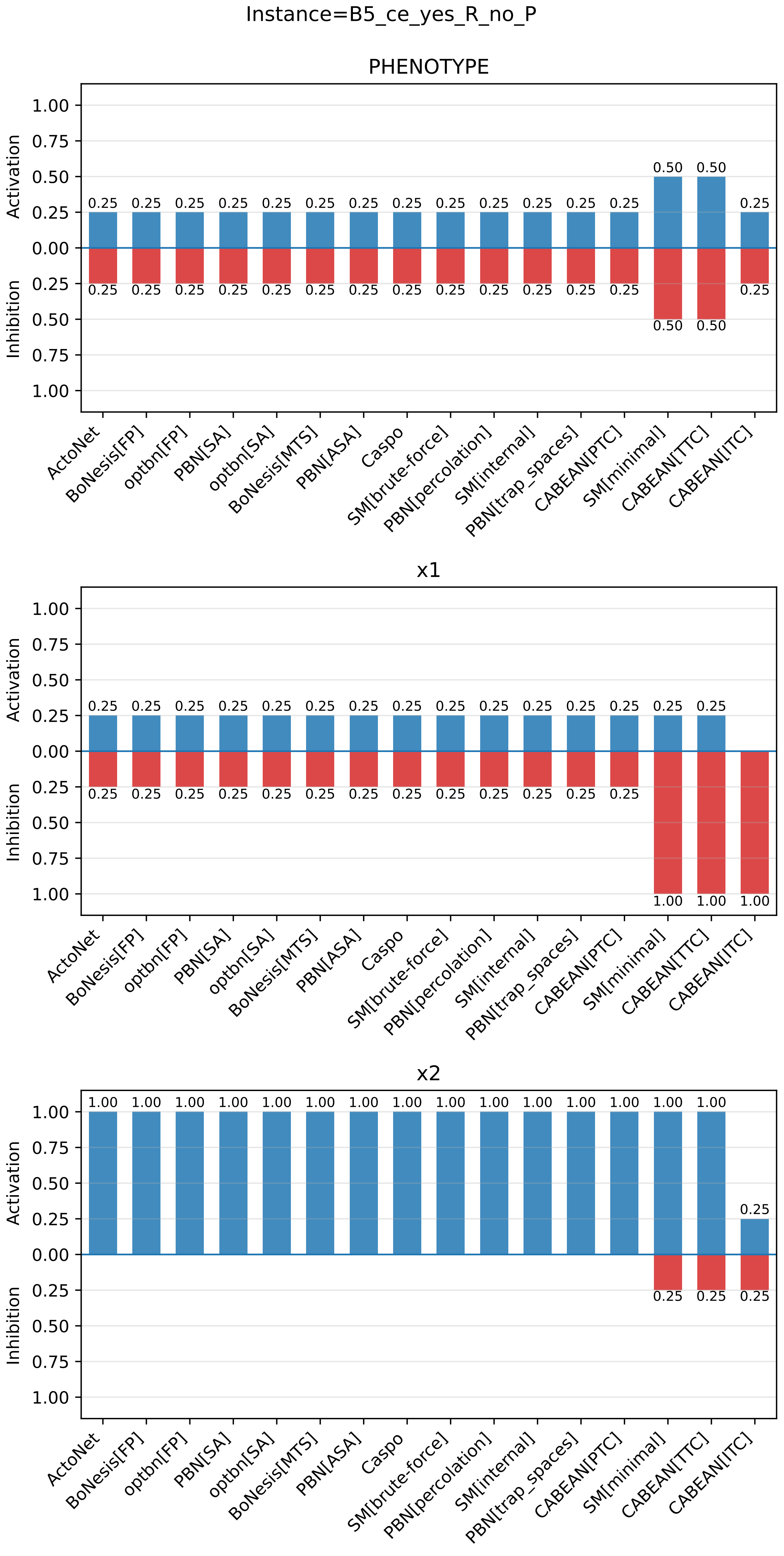

### _score_full.pdf

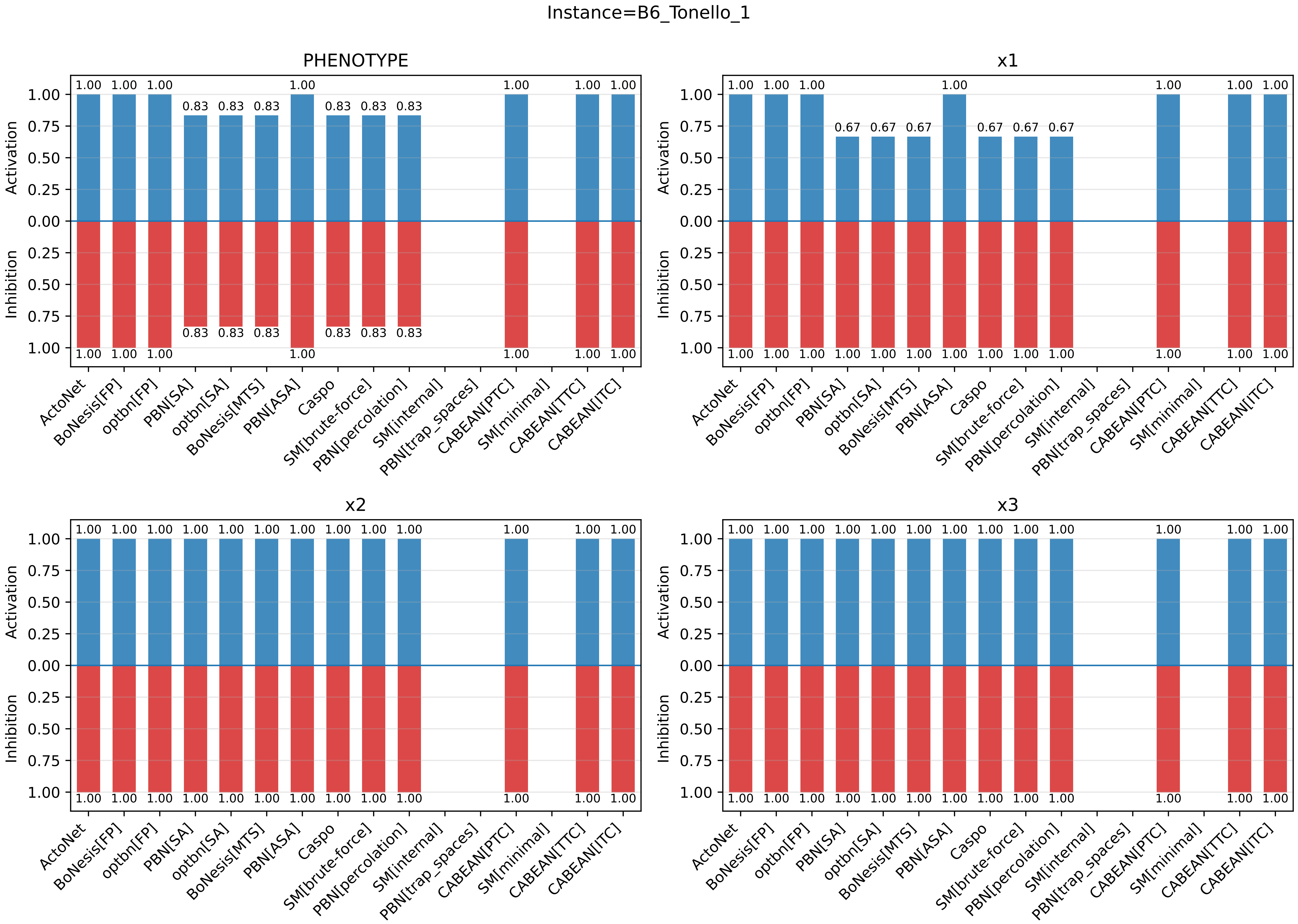

### _score_full.pdf

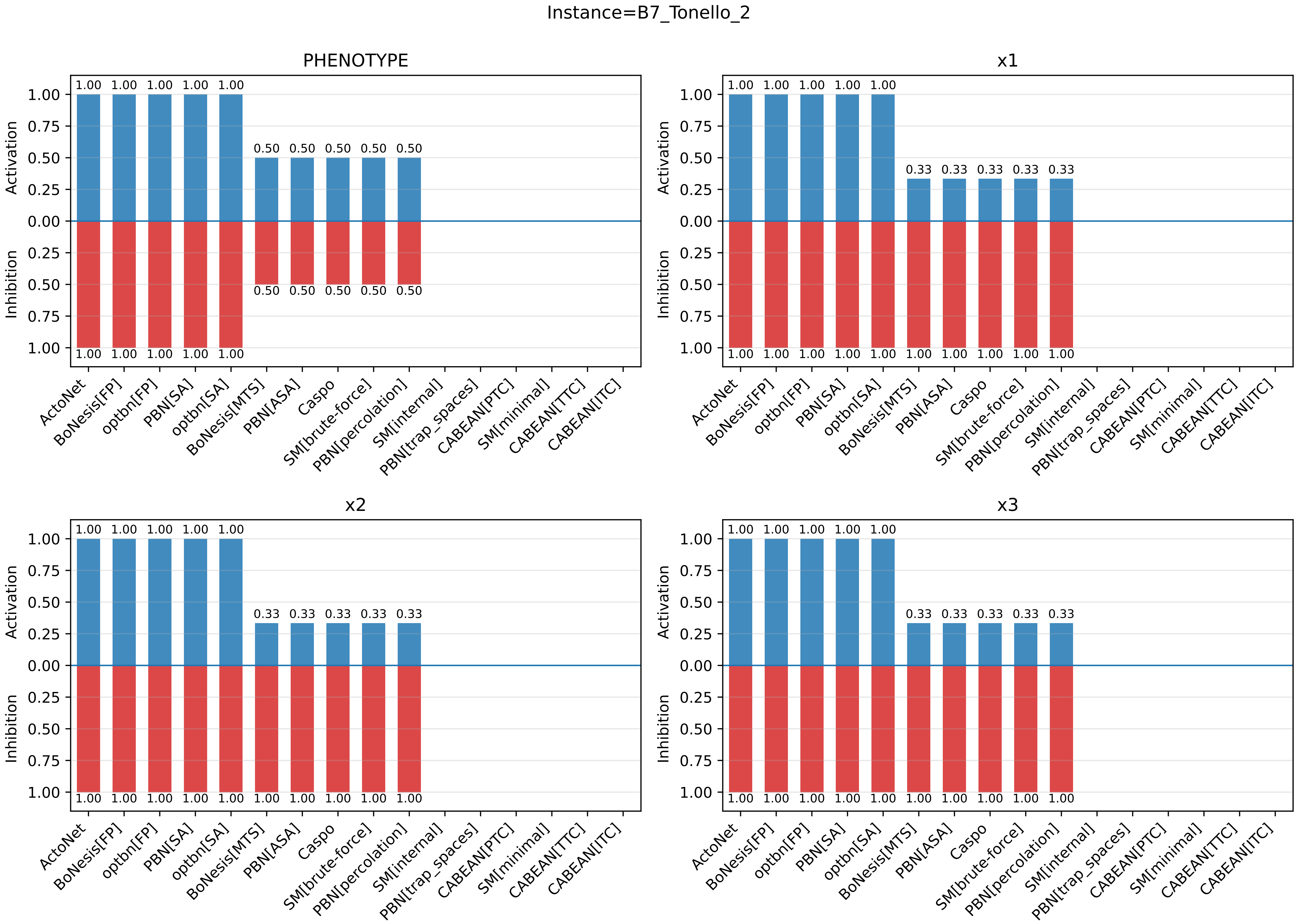

### _score_full.pdf

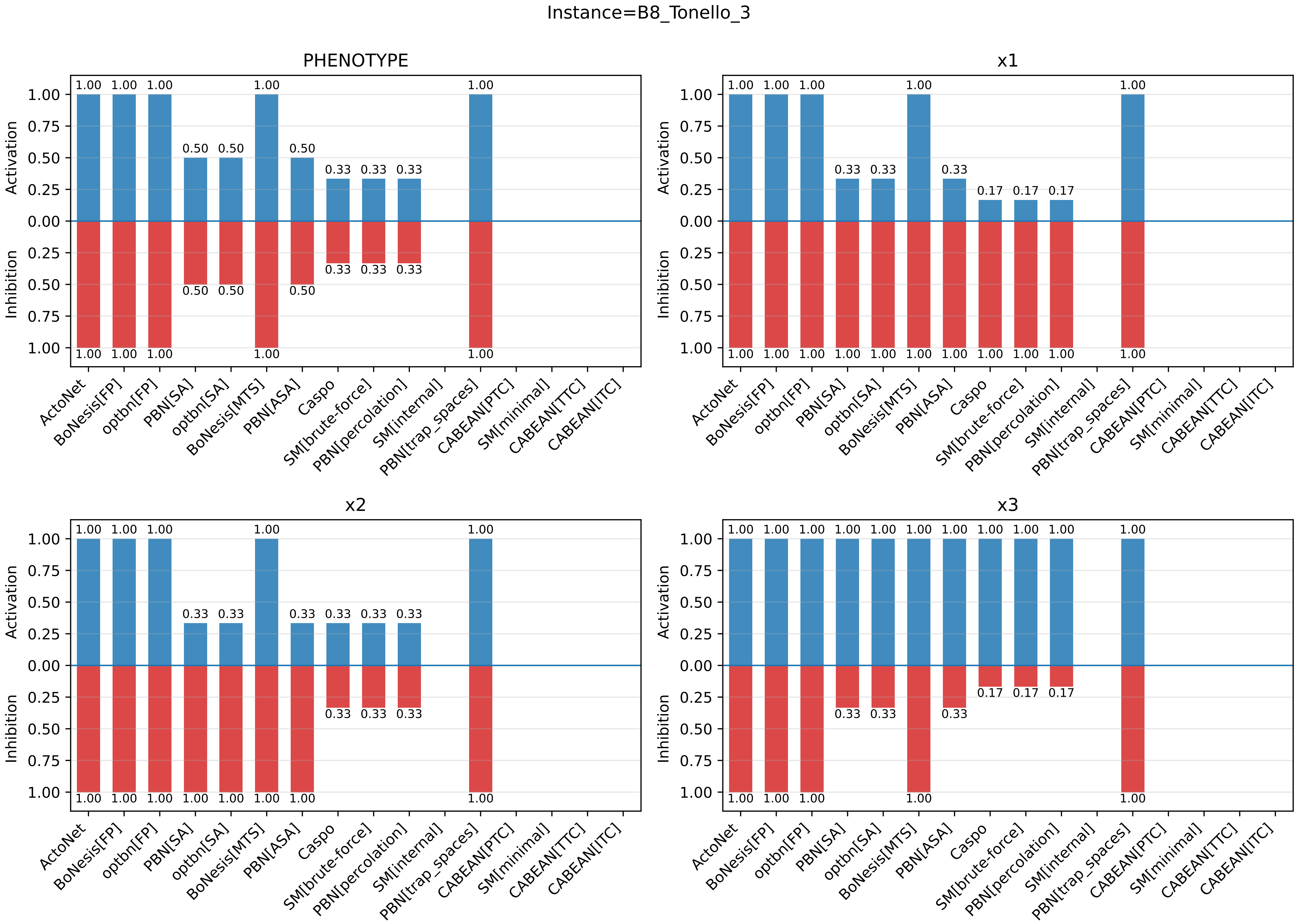

### _score_histogram.pdf

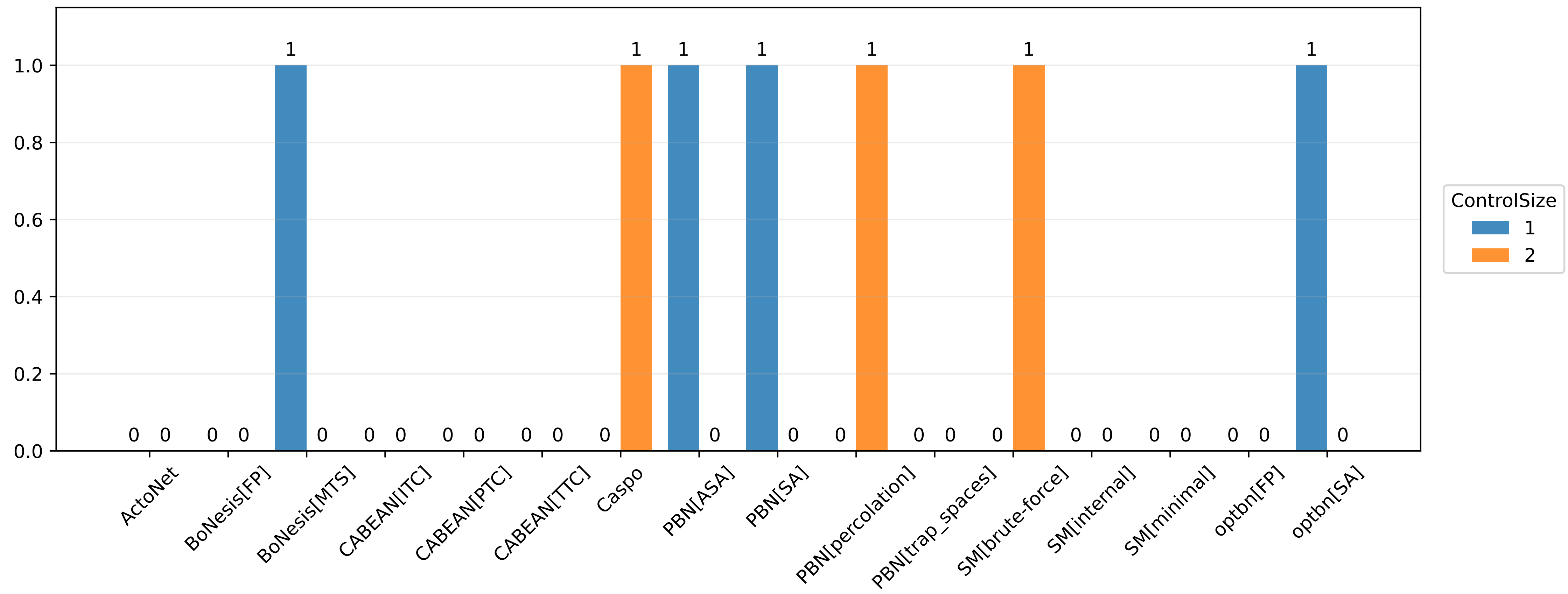

### _score_histogram.pdf

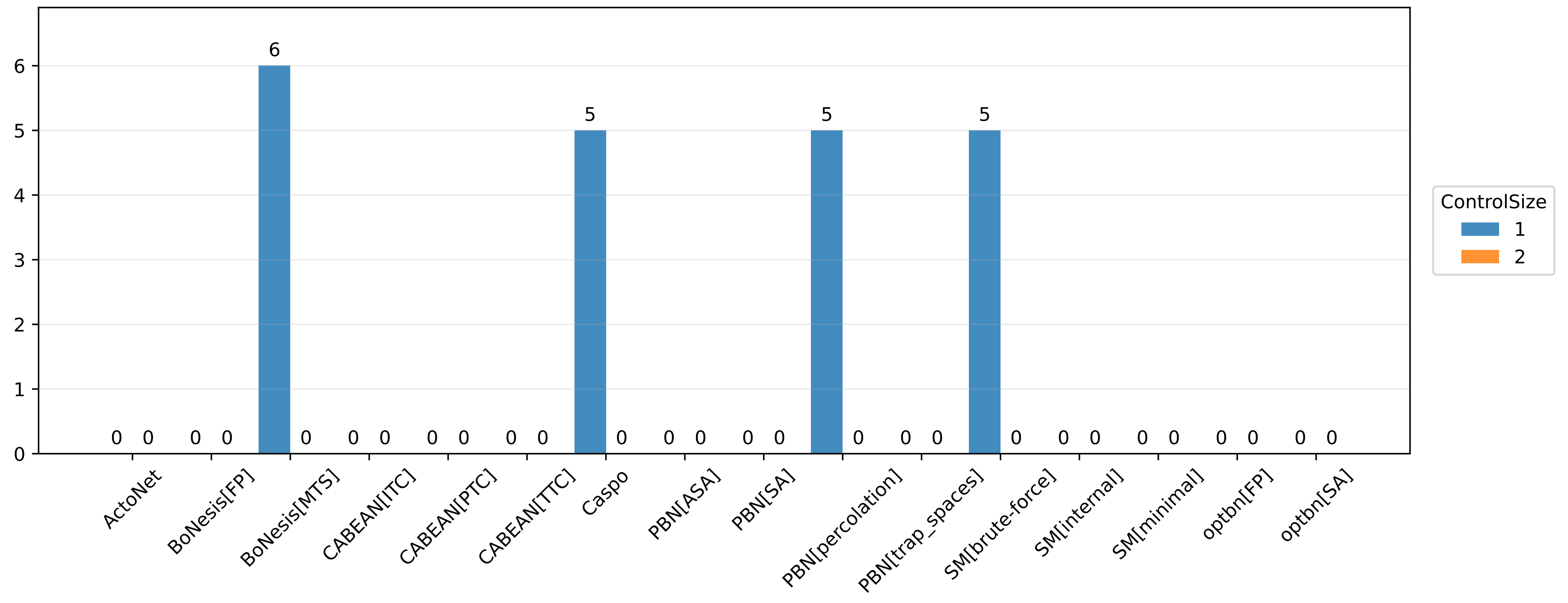

### _score_histogram.pdf

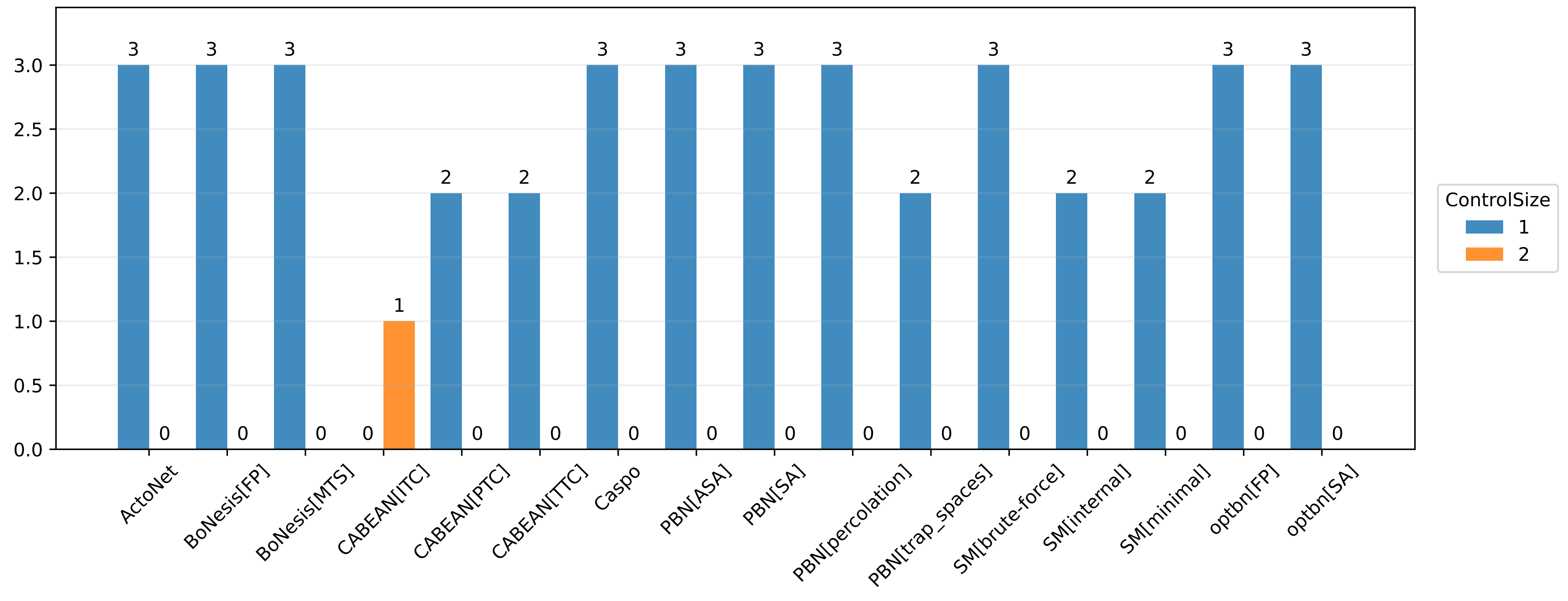

### _score_histogram.pdf

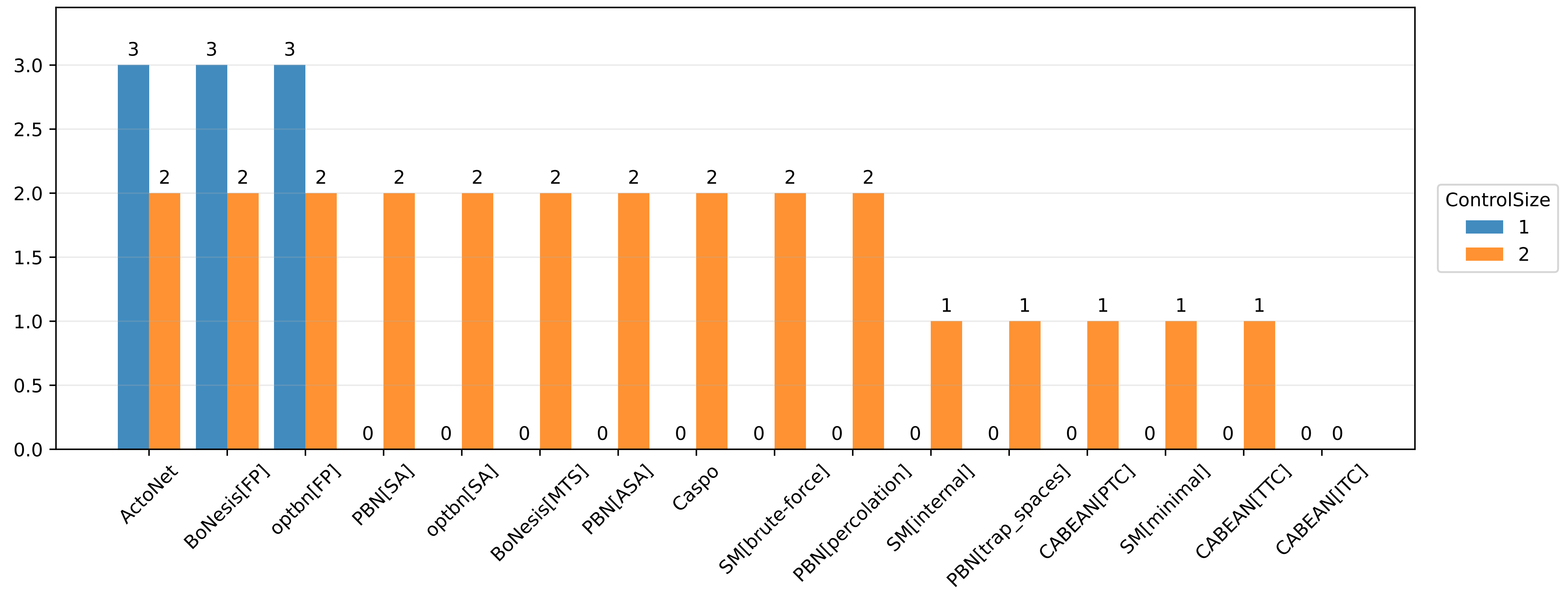

### _score_histogram.pdf

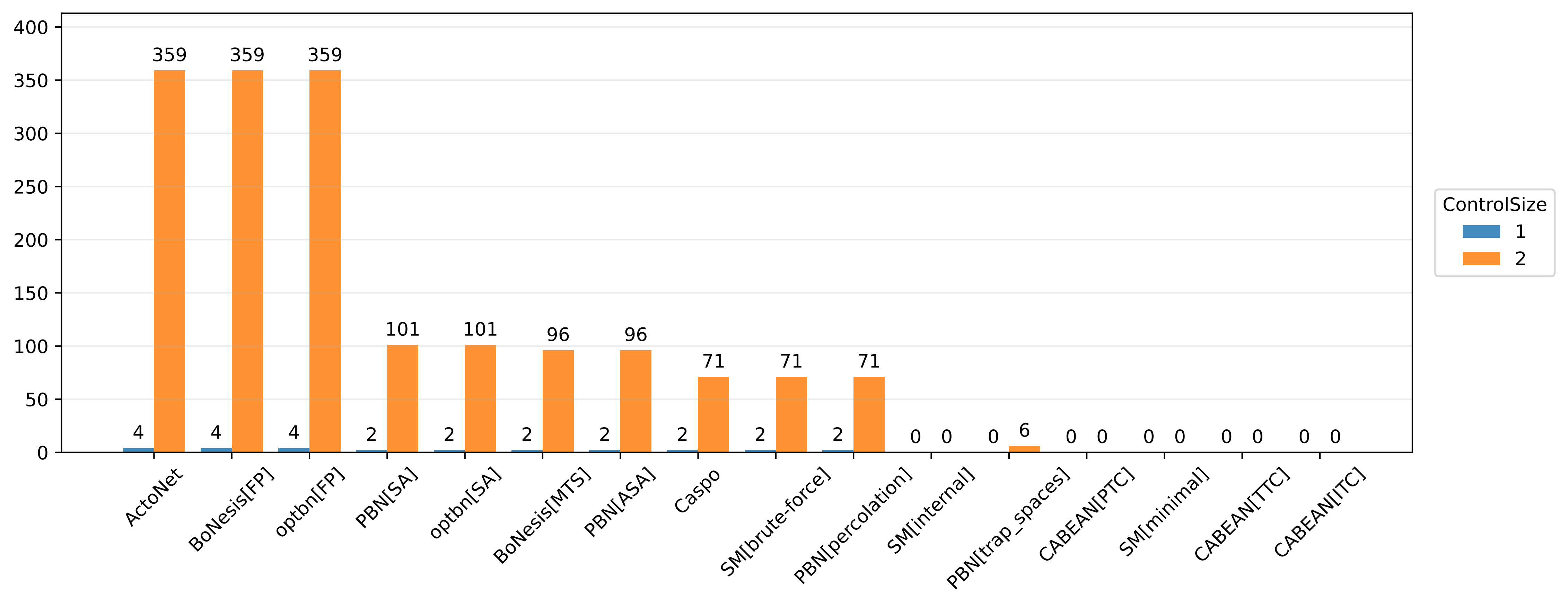

### _score_histogram.pdf

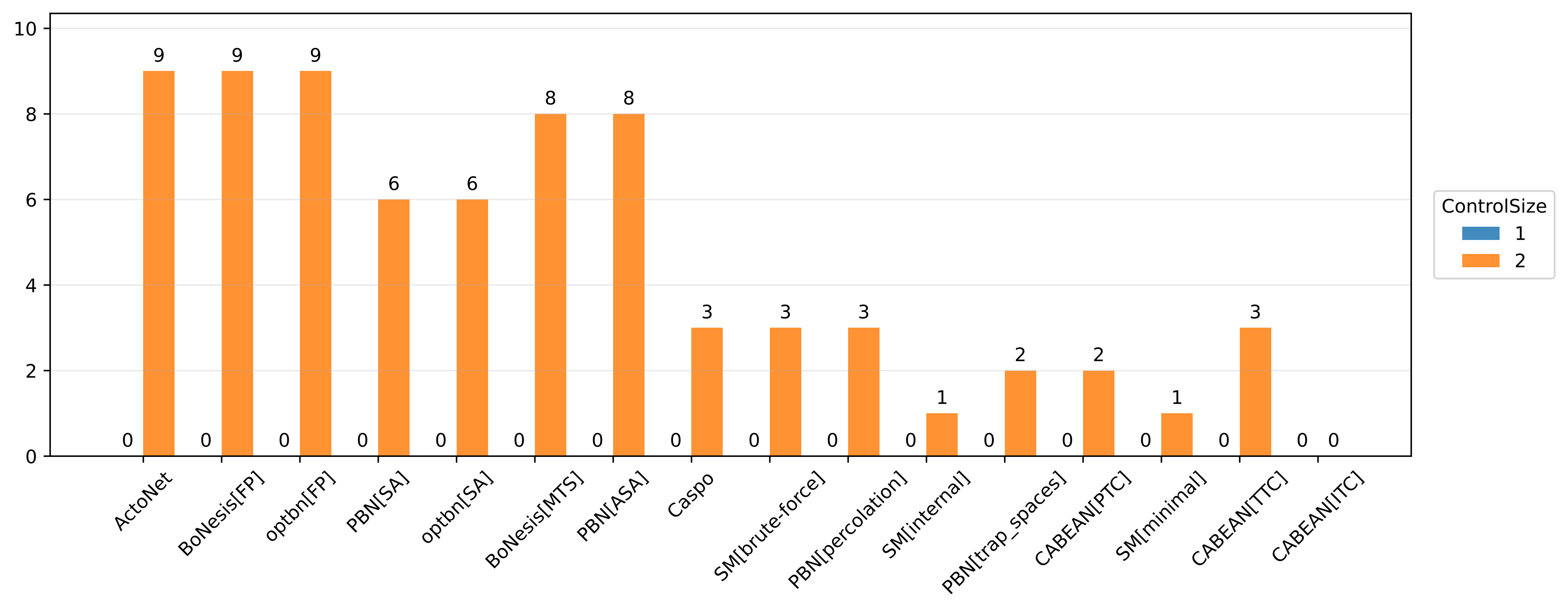

### _score_histogram.pdf

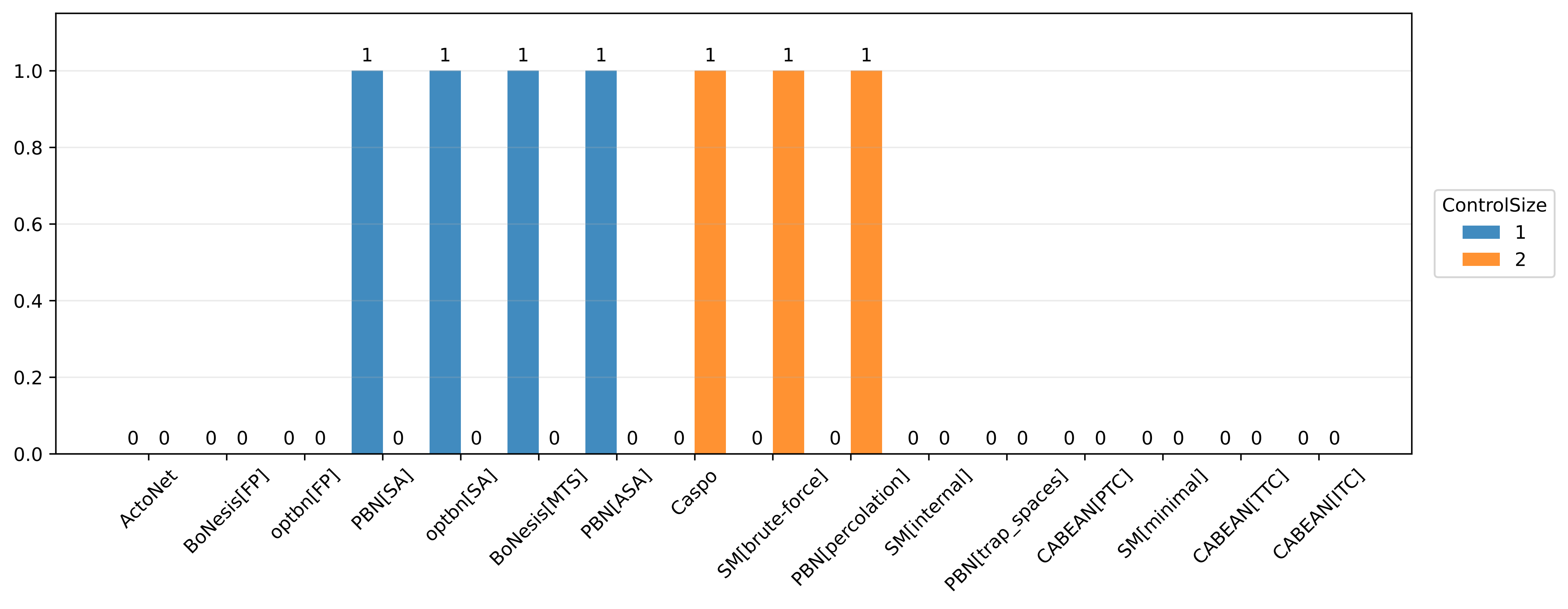

### _score_histogram.pdf

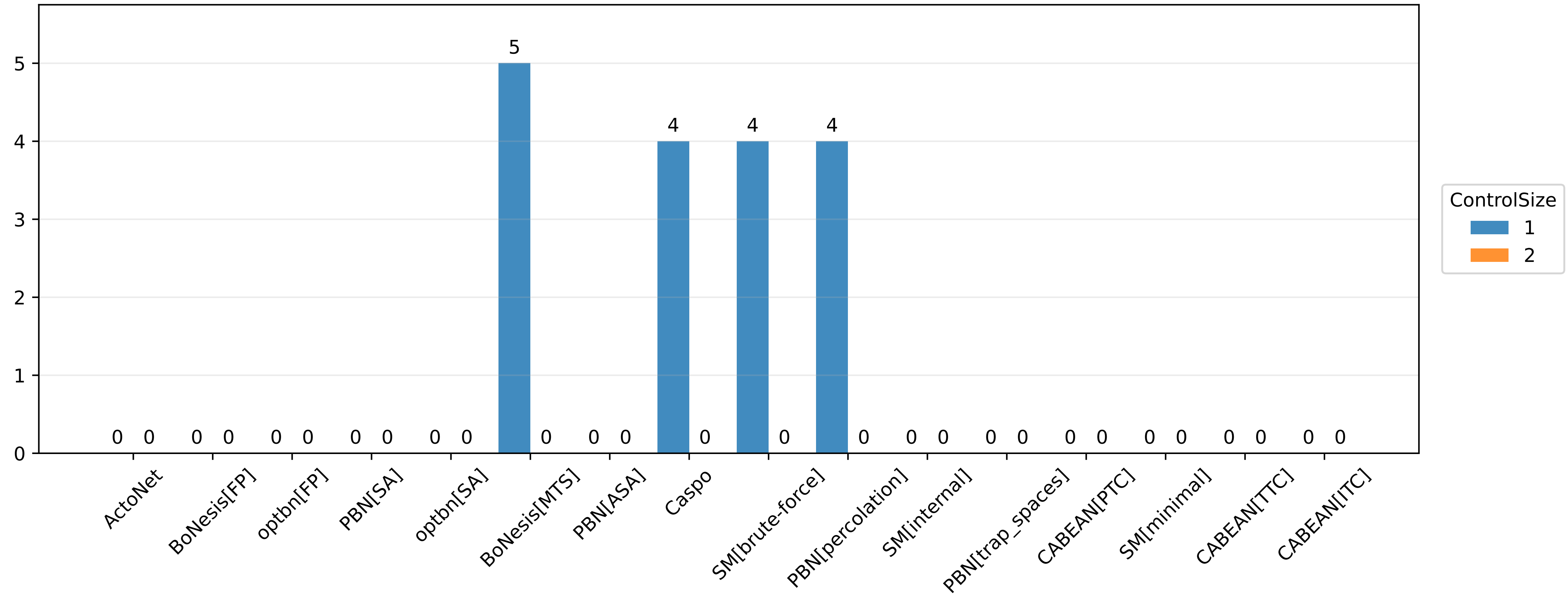

### _score_histogram.pdf

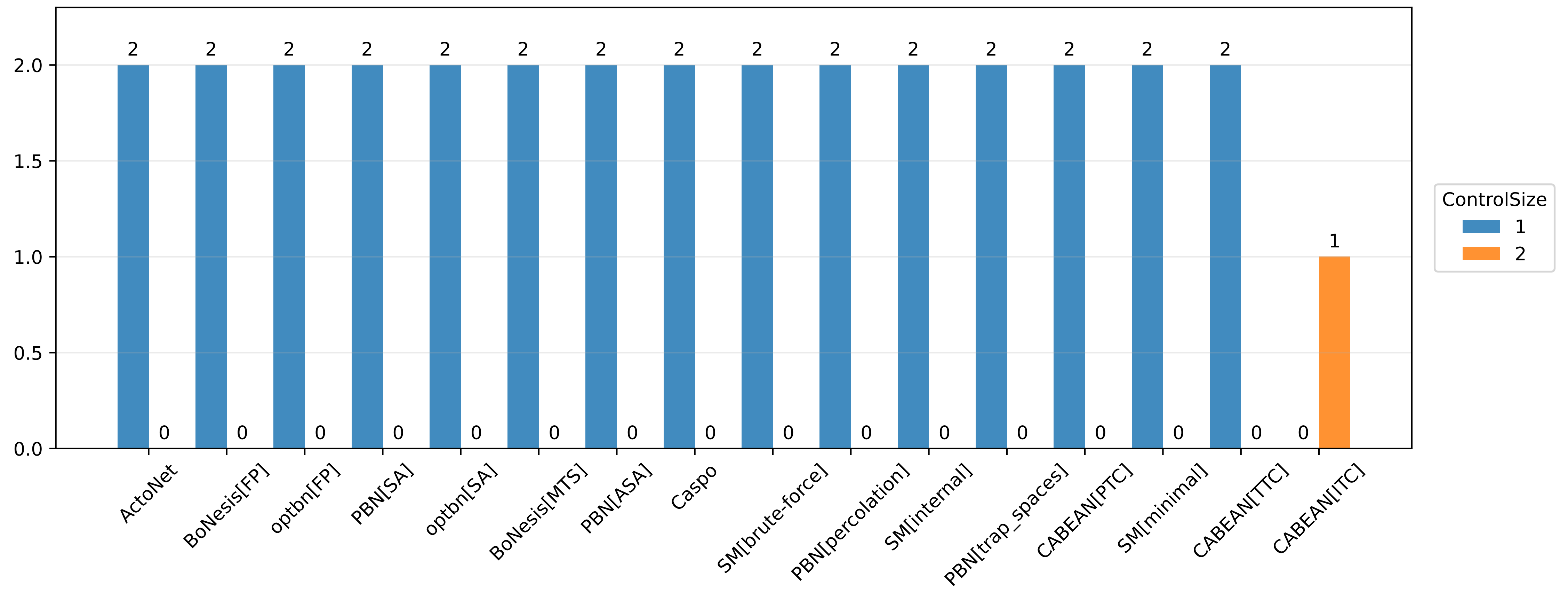

### _score_histogram.pdf

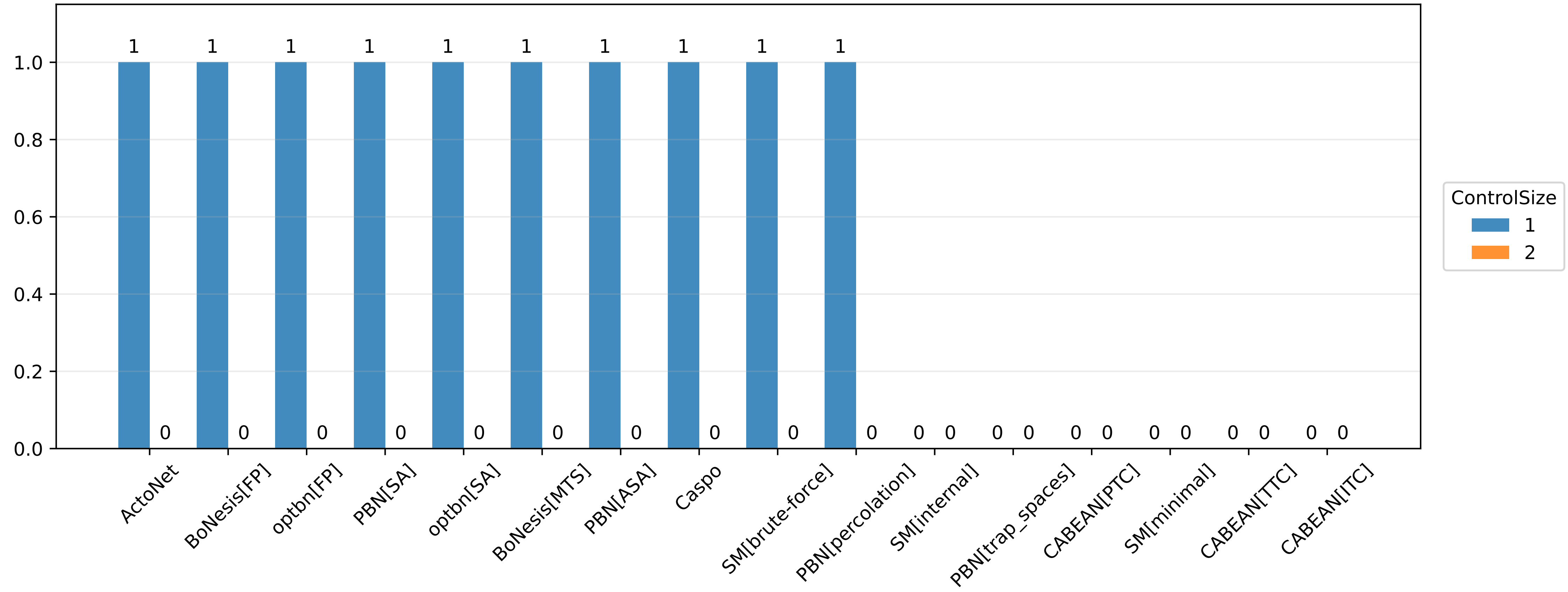

### _score_histogram.pdf

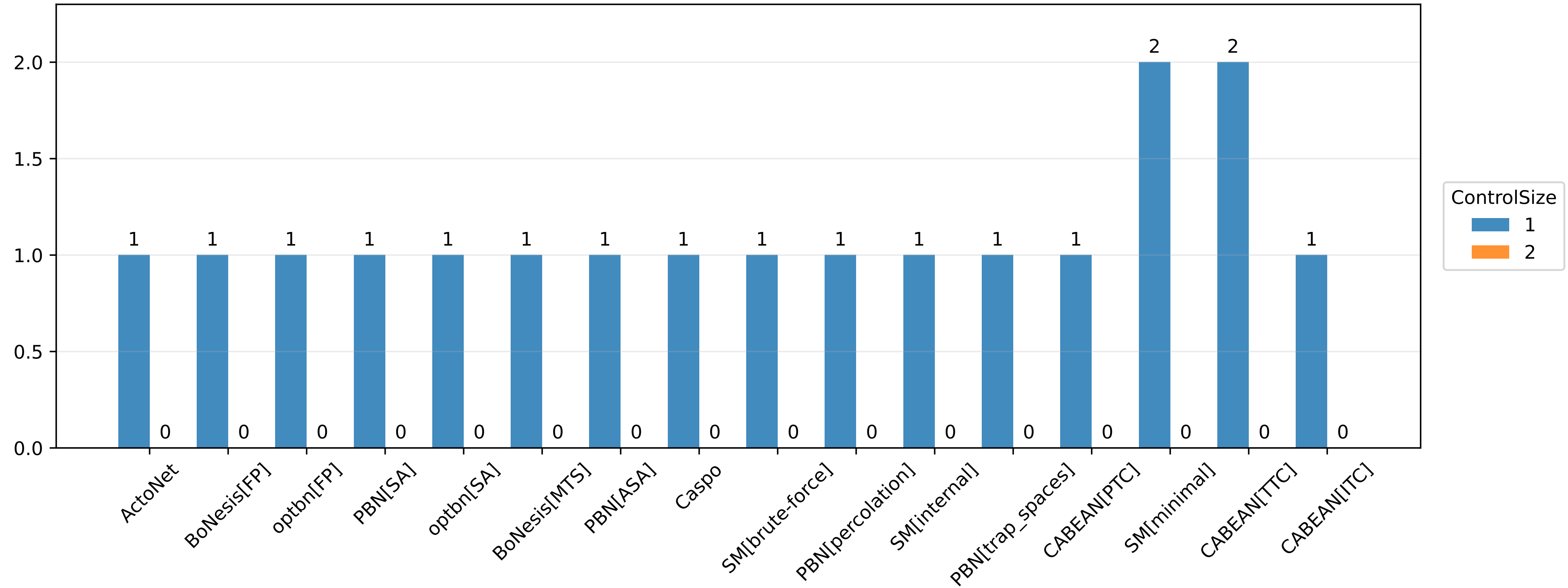

### _score_histogram.pdf

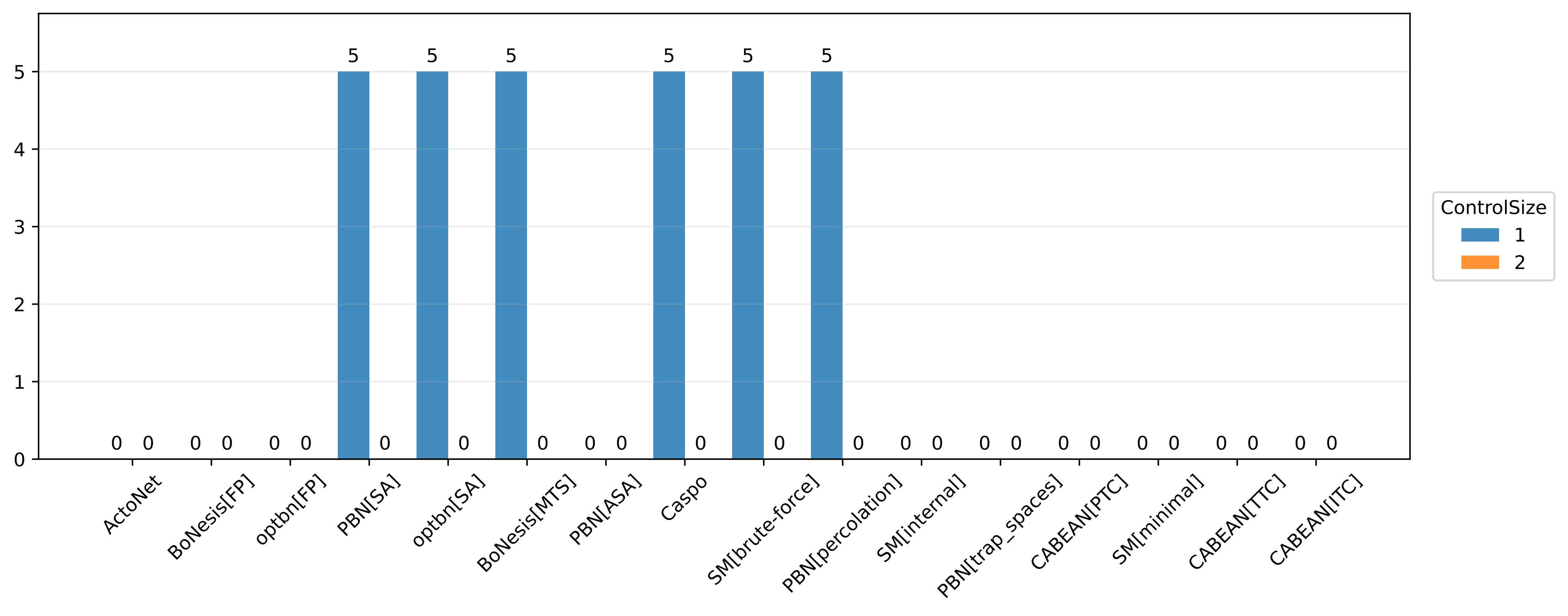

### _score_histogram.pdf

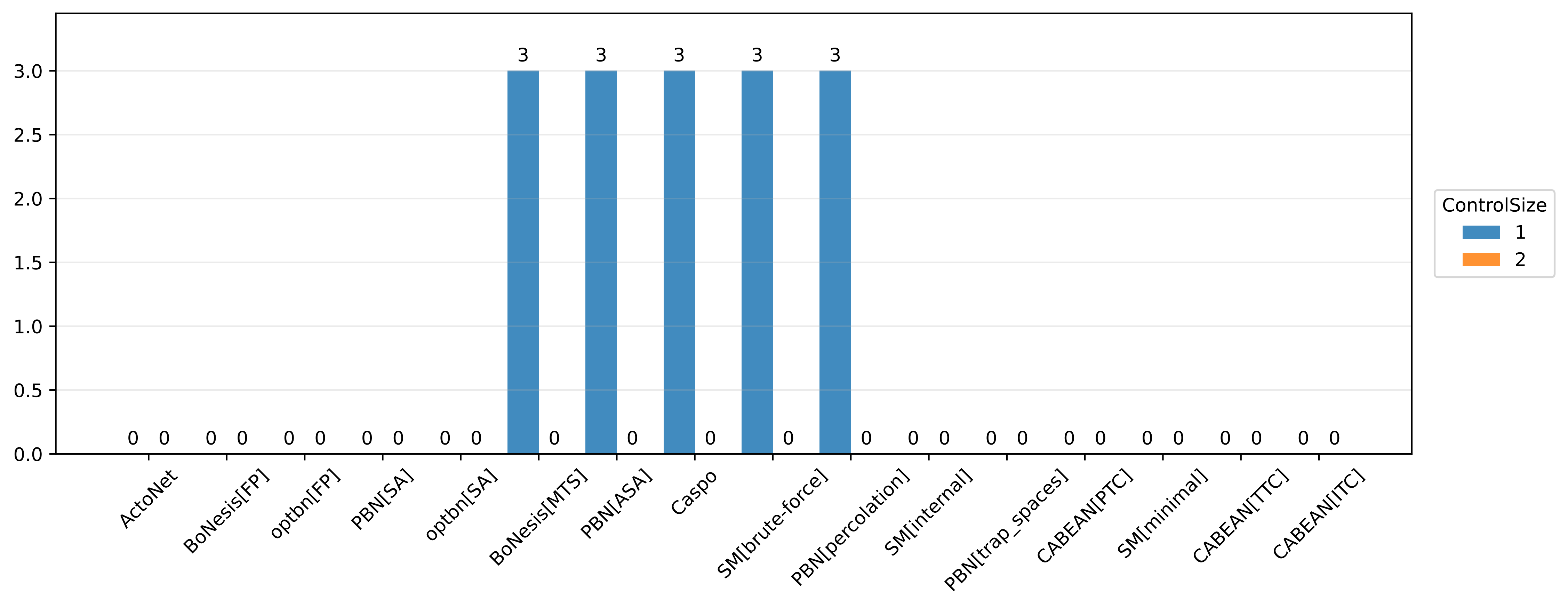

### _score_histogram.pdf

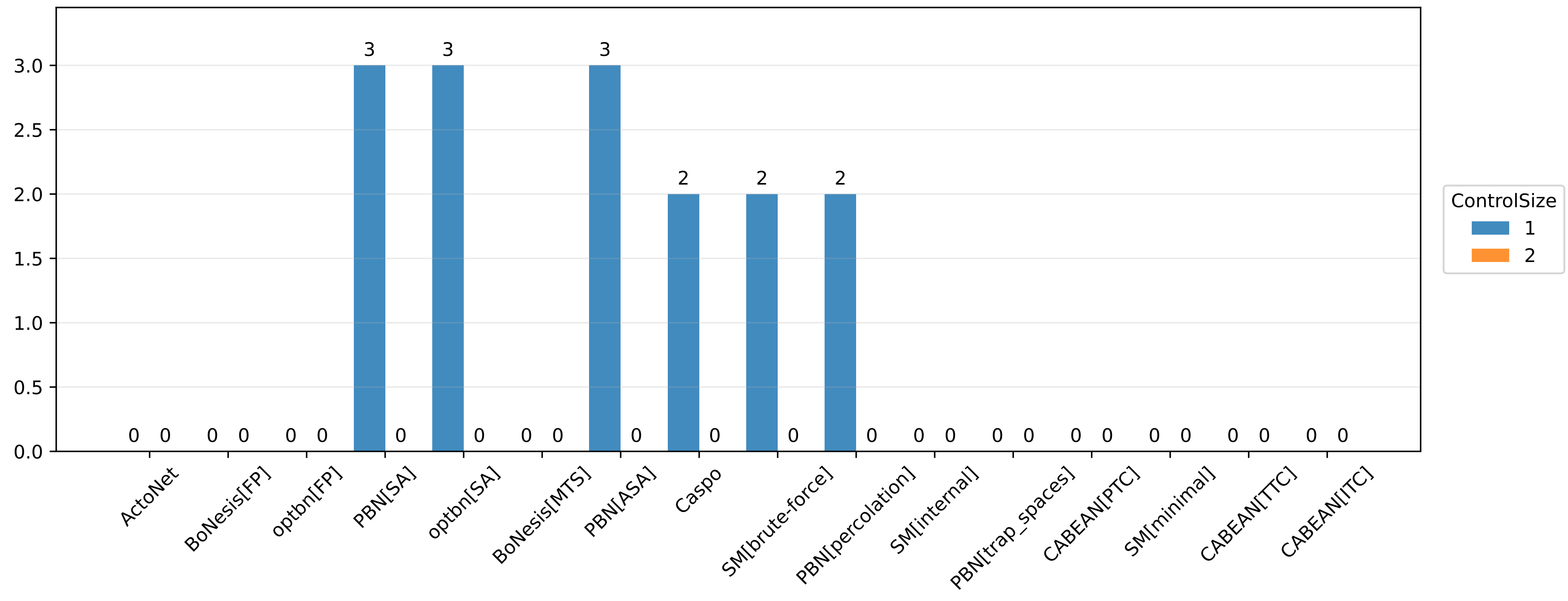

### _score_summary.pdf

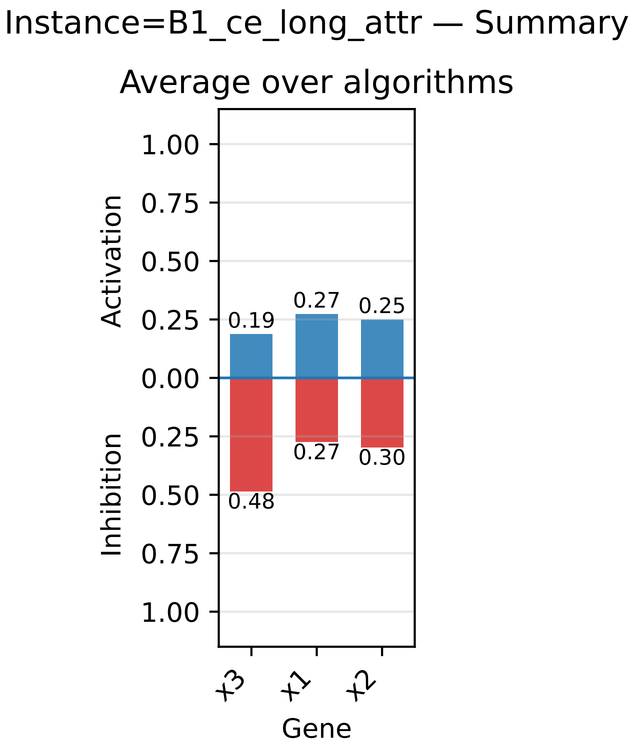

### _score_summary.pdf

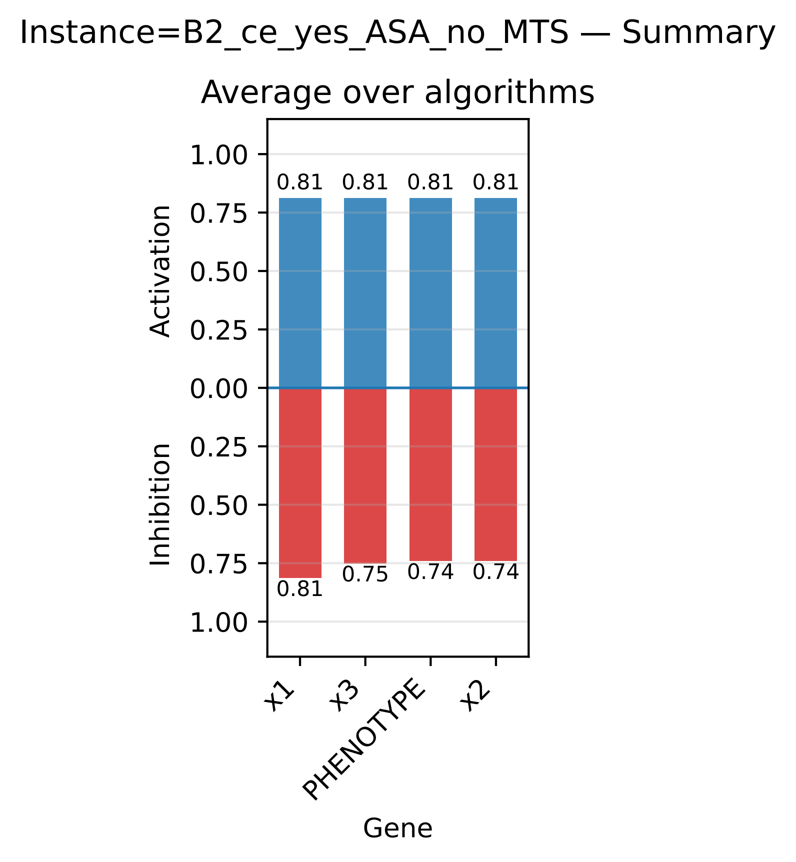

### _score_summary.pdf

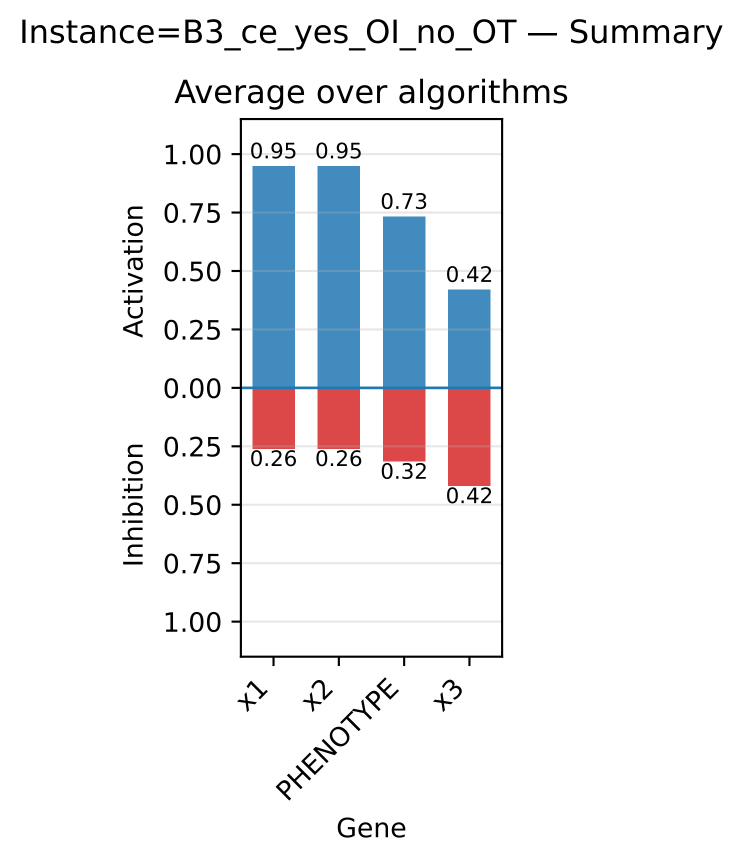

### _score_summary.pdf

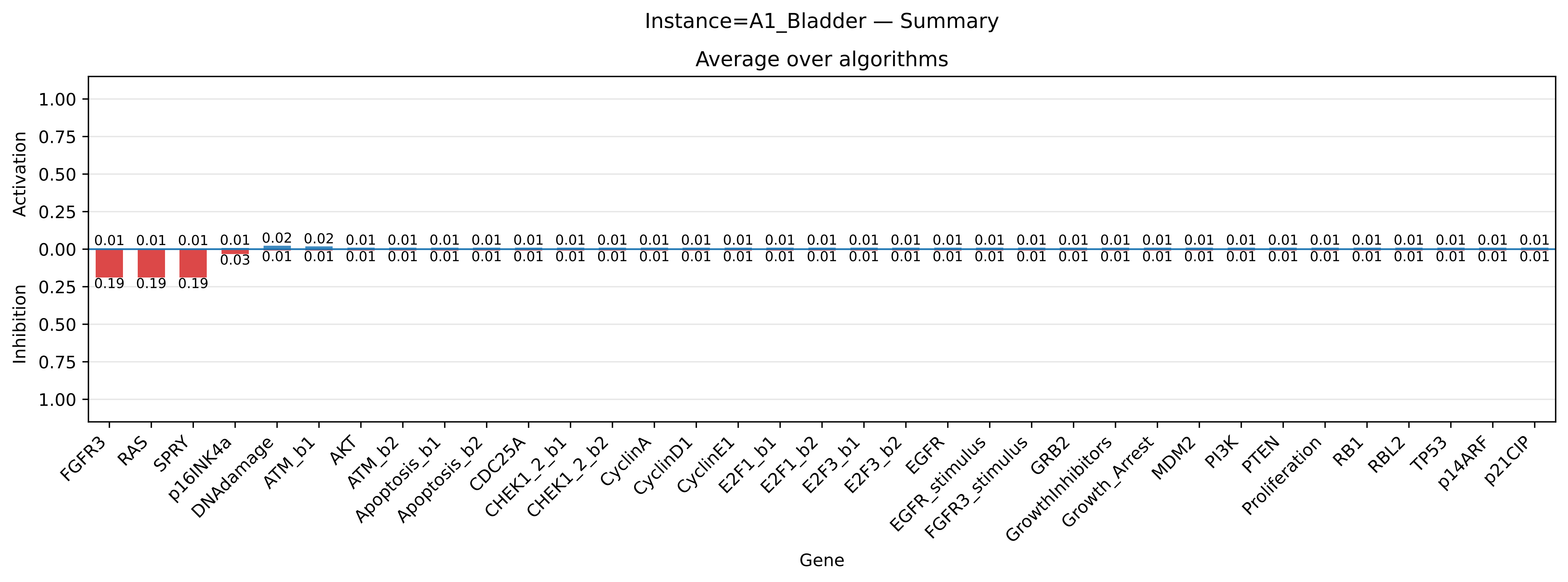

### _score_summary.pdf

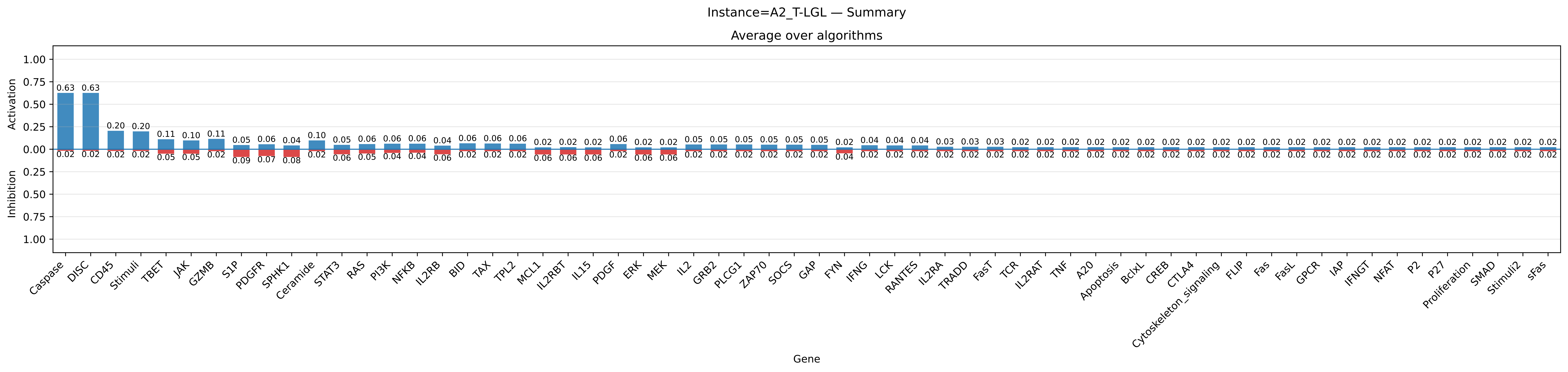

### _score_summary.pdf

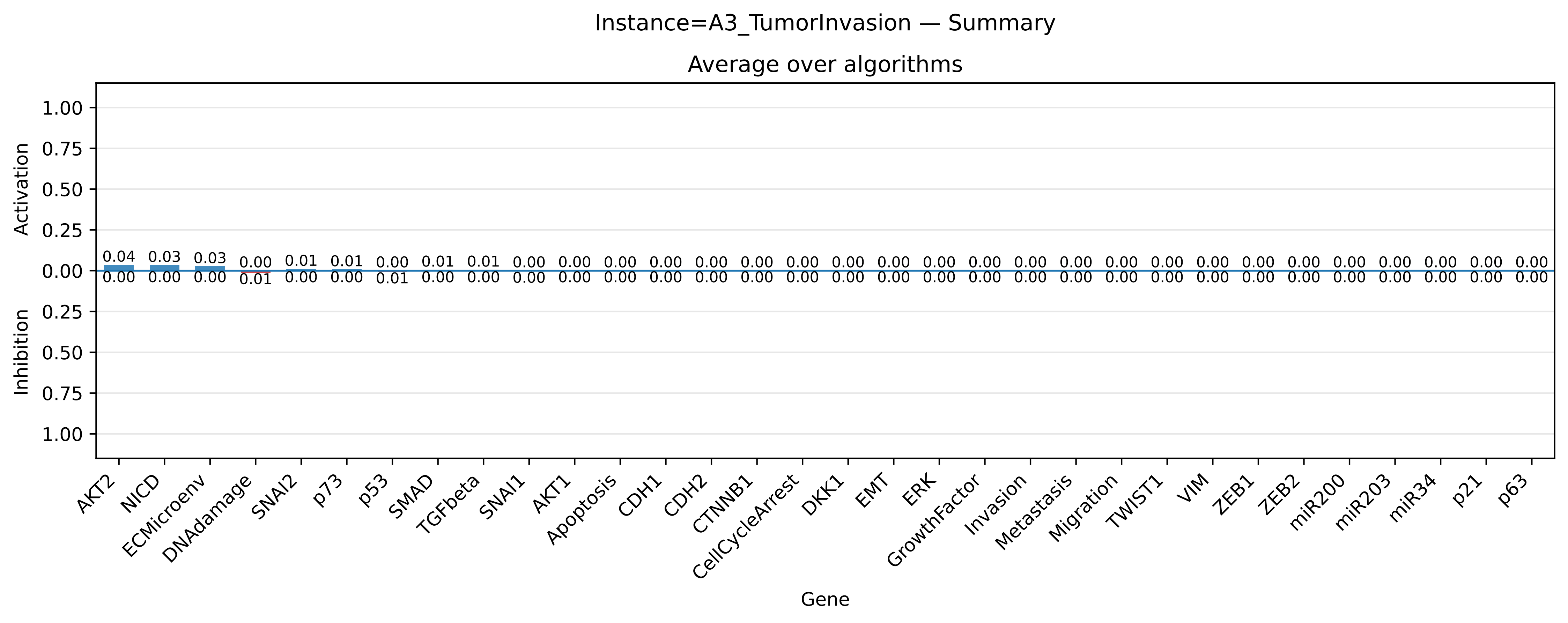

### _score_summary.pdf

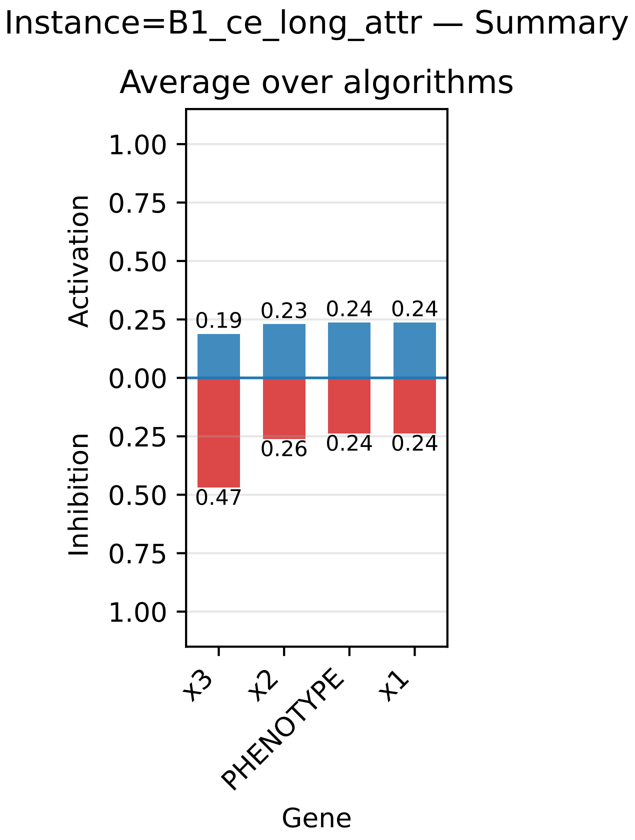

### _score_summary.pdf

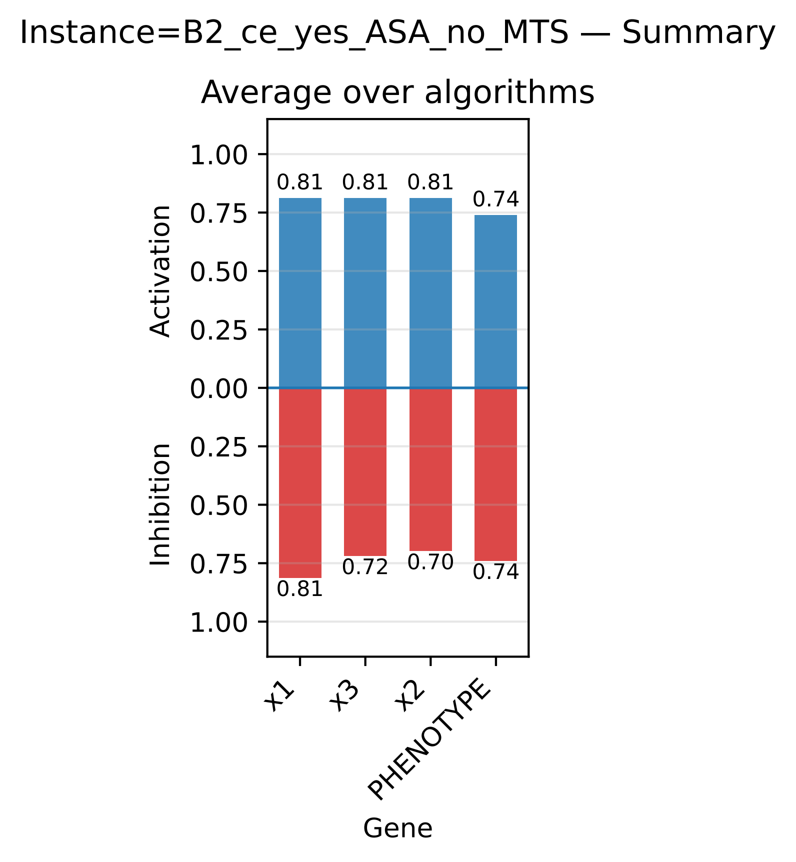

### _score_summary.pdf

Instance=B3_ce_yes_OI_no_OT — Summary

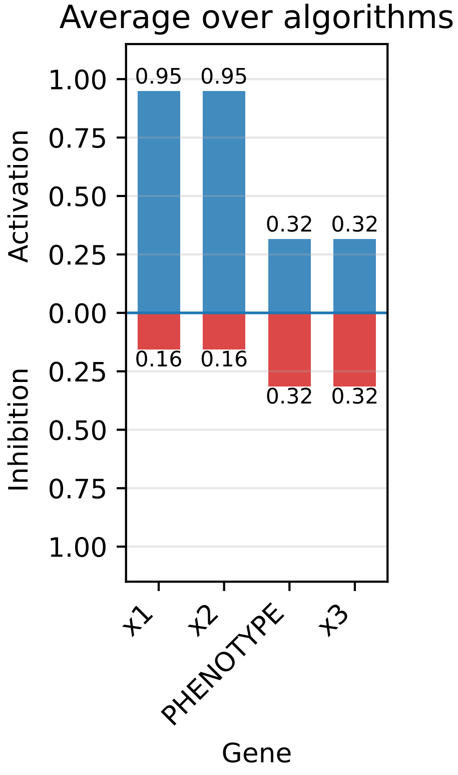

### _score_summary.pdf

#### Instance=B4_ce_yes_P_no_R — Summary

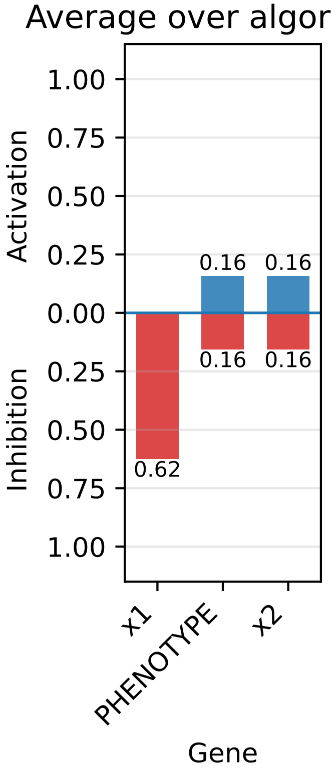

### _score_summary.pdf

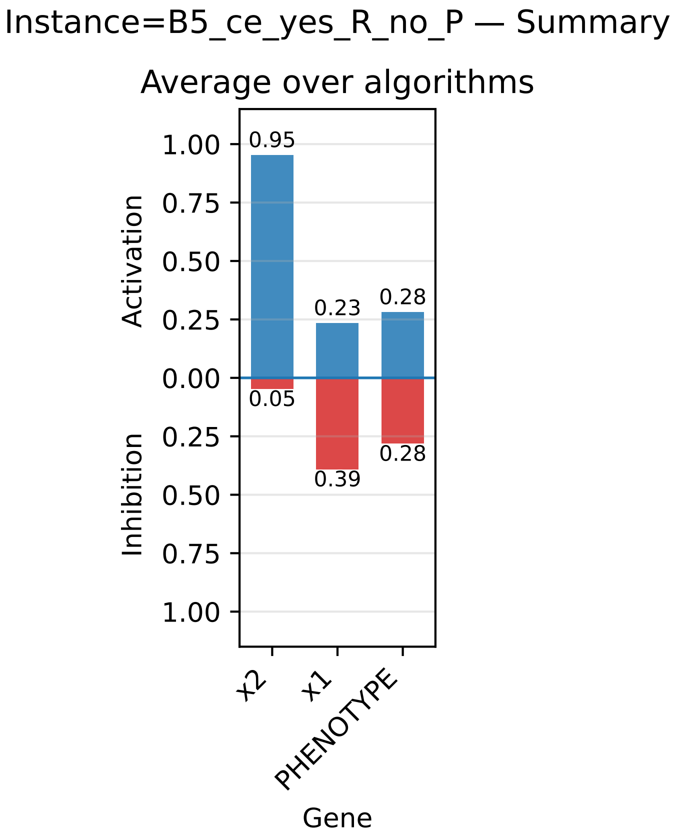

### _score_summary.pdf

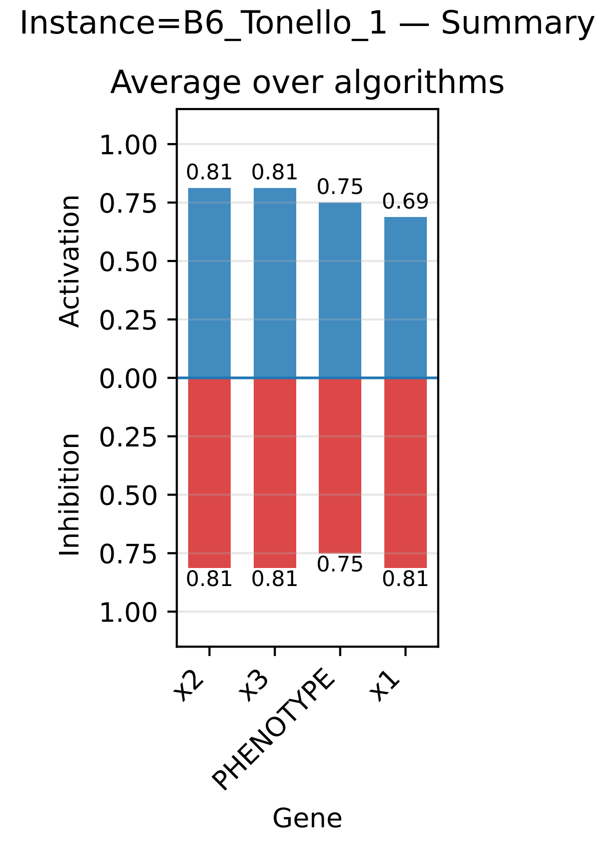

### _score_summary.pdf

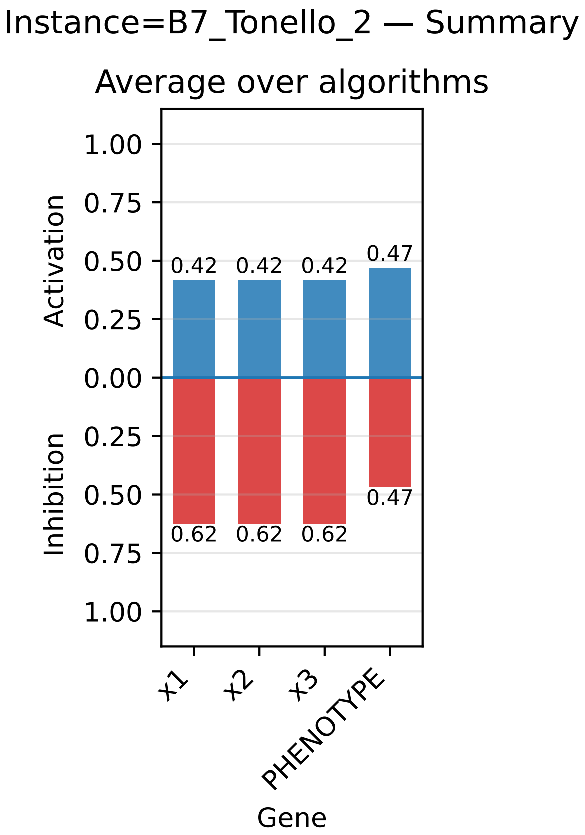

### _score_summary.pdf

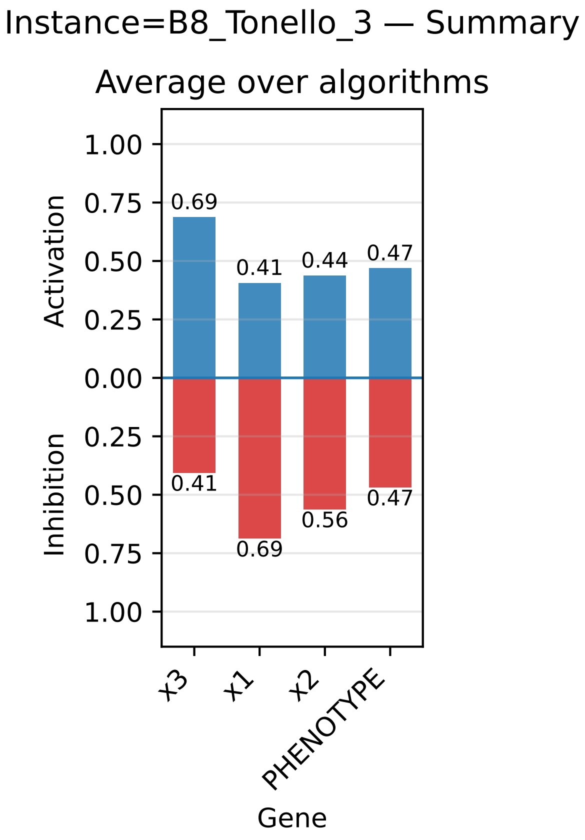
